# Single-cell full-length transcriptome of human lung reveals genetic effects on isoform regulation beyond gene-level expression

**DOI:** 10.64898/2026.03.27.714873

**Authors:** Bolun Li, Thong Luong, Elelta Sisay, Jinhu Yin, Zixuan Eleanor Zhang, Maryam Vaziripour, Ju Hye Shin, Yongmei Zhao, Bao Tran, Jinyoung Byun, Yafang Li, Chia Han Lee, Maura O’Neill, Thorkell Andresson, Yoon Soo Chang, Steven Gazal, Maria Teresa Landi, Nathaniel Rothman, Erping Long, Qing Lan, Christopher I Amos, Anny Xiaobo Zhou, Tongwu Zhang, Jin Gu Lee, Jianxin Shi, Nicholas Mancuso, Jun Xia, Haoyu Zhang, Eun Young Kim, Jiyeon Choi

**Affiliations:** Division of Cancer Epidemiology and Genetics, National Cancer Institute, National Institutes of Health, Bethesda, MD, USA; Department of Genetics, Perelman School of Medicine, University of Pennsylvania, Philadelphia, PA, USA; Center for Genomic and Precision Medicine, Texas A&M Health, Houston, TX, USA; Department of Internal Medicine, Yonsei University College of Medicine, Seoul, Republic of Korea; NCI Sequencing Facility, Frederick, MD, USA; Department of Internal Medicine, University of New Mexico, Albuquerque, NM, USA; NCI CCR Protein Characterization Laboratory, Frederick, MD, USA; Institute of Basic Medical Sciences, Chinese Academy of Medical Sciences and Peking Union Medical College, Beijing, China; Division of Pulmonary and Critical Care, Department of Medicine; Department of Genome Sciences, University of Virginia, Charlottesville, VA, USA; Department of Thoracic and Cardiovascular Surgery, Yonsei University College of Medicine, Seoul, Republic of Korea; Center for Genetic Epidemiology, Department of Population and Public Health Sciences, Keck School of Medicine, University of Southern California, Los Angeles, CA, USA

## Abstract

Genetic regulation of splicing uniquely contributes to trait-associated genome-wide association studies (GWAS) signals. However, quantitative trait loci (QTL) analysis using short-read sequencing of bulk tissues fails to capture full-length and cell-type-specific isoforms. Here, we present an isoform-level lung cell atlas from 129 never-smoking Korean women using single-cell long-read RNA-sequencing, identifying abundant unannotated and cell-type-specific isoforms. Isoform-level signatures of 37 lung cell types display a larger difference and therefore improve cell-type classification compared to gene-level expression. Notably, isoform-QTLs (isoQTLs) detect unannotated and/or cell-type-specific isoforms with independent genetic regulation from expression-QTL (eQTL), supported by enriched splicing functional elements. IsoQTLs nominate susceptibility isoforms from previously unexplained lung function and cancer GWAS loci, via eQTL-independent signals. We highlight a potentially functional novel variant of *PPIL6* in multiciliated cells underlying lung cancer risk through alternative splicing. This isoform-level resource advances our understanding of cell-type-specific isoform regulation and its contribution to lung traits and diseases.

## Introduction

Transcript isoform variation creates functional diversity in protein isoforms or influences total gene expression levels through the turnover of less stable isoforms ^1^. Alternative RNA splicing and isoform composition dynamics could play an essential role in organogenesis, and misregulation of this process could lead to various human diseases^2,3^, including cancer and other chronic diseases. However, the full diversity of isoform structures in different tissues and cell types is not completely understood due to the dearth of tissue-based and cell-type-specific full-length transcriptome datasets in diverse populations.

Consistent with the importance of isoform functionality, it is increasingly evident that unique genetic contribution to splicing regulation underlies complex diseases. Although tissue-based studies identified comparable numbers of splicing junction usage quantitative trait loci (splicing QTL or sQTL) and expression QTLs (eQTLs), they showed limited overlap of credible set variants. Consistently, sQTLs contributed to explaining various trait-associated GWAS signals independent from eQTLs^3^. For a typical sQTL mapping using short-read RNA-sequencing (RNA-seq) data, either isoform expression is estimated using probabilistic models^4,5^, or relative splice junction usage is treated as a proxy phenotype. However, the molecular effects of these junction-based sQTLs can be elusive; mapping them to a specific isoform is challenging due to shared junctions and incomplete reference annotation.

Addressing these limitations, recent full-length transcriptomes of human tissues using long-read RNA-seq reported hundreds of thousands novel isoforms accounting for 26%-77% of total isoforms^6–8^. For example, an isoform-level transcriptome using 14 tissues from GTExexhibited tissue-specificity of isoforms and included a substantial proportion of novel isoforms, with a subset being validated at the peptide level ^6^. Moreover, emerging single-cell long-read RNA-seq studies in brain tissues demonstrated isoform specificity across brain cell types ^7,9^, including disease-relevant novel isoforms ^7^, and observed an overlap of published sQTLs with cell-type-specific disease gene ^10^.

Although single-cell eQTL datasets of several human tissue types has highlighted prevalent cell-type-specific genetic regulation contributing to trait-associated GWAS loci ^11–13^, single-cell isoform-level QTL datasets are lacking due to high cost and sparsity of single-cell long-read RNA-seq. Rare cases of single-cell sQTL studies either used single-cell 5’ short-read RNA-seq^14^ or leveraged a full-length transcriptome reference to interpret junction-based sQTLs ^8^, where isoform estimation is still vulnerable to errors especially for longer and complex isoforms ^8^. Consequently, whether cell-type-specific isoforms are different from gene-level observation and whether isoform QTLs contribute to GWAS signals of complex traits independent from eQTLs have not been systematically investigated.

Lung cancer is a common and complex disease including histological types originating from distinct epithelial cell types. Although smoking is a well-known risk factor, GWAS have identified more than 60 risk loci, including those distinct to histological types and populations^15–17^. Lung adenocarcinoma (LUAD), the most common histological type, mainly arises from alveolar type 2 (AT2) of lung alveoli ^18–20^, with alveolar type 1 (AT1), club, and alveolar transitional cells/states also being implicated^21,22^. These cell types have relevance to the regeneration of alveoli that contributes to broader lung function traits and chronic lung diseases ^23^. Indeed, genetic control of lung function traits, including those defining chronic obstructive pulmonary disease (COPD), can contribute to lung cancer risk ^24^. Large-scale lung function GWAS identified >1,000 loci in multi-ancestry studies ^25,26^, and genetic correlation as well as overlap of GWAS findings between lung cancer and lung functions or COPD have been observed ^24,27^. Although previous bulk-tissue based lung eQTLs partially characterized lung cancer and lung function GWAS loci, the contribution of isoform diversity and sQTLs ^28^ as well as single-cell QTLs ^11,12^ have been less explored, especially in non-European populations.

In this study, we generated a single-cell full-length transcriptome of tumor-distant normal lung tissues from 129 Korean never-smoking women to investigate the genetic effects on transcript isoform regulation (Fig. 1). This isoform-level lung cell atlas revealed the signatures of isoform expression across 37 lung cell types. We mapped isoform-level QTLs (isoQTLs) across 33 cell types and further deciphered potential splicing relevant functions of isoQTLs distinct from eQTLs. By integrating isoQTLs with lung cancer and lung function GWAS, we uncovered the context-specific susceptibility isoforms and their contribution to GWAS signals independent from eQTLs.

**Figure 1.**
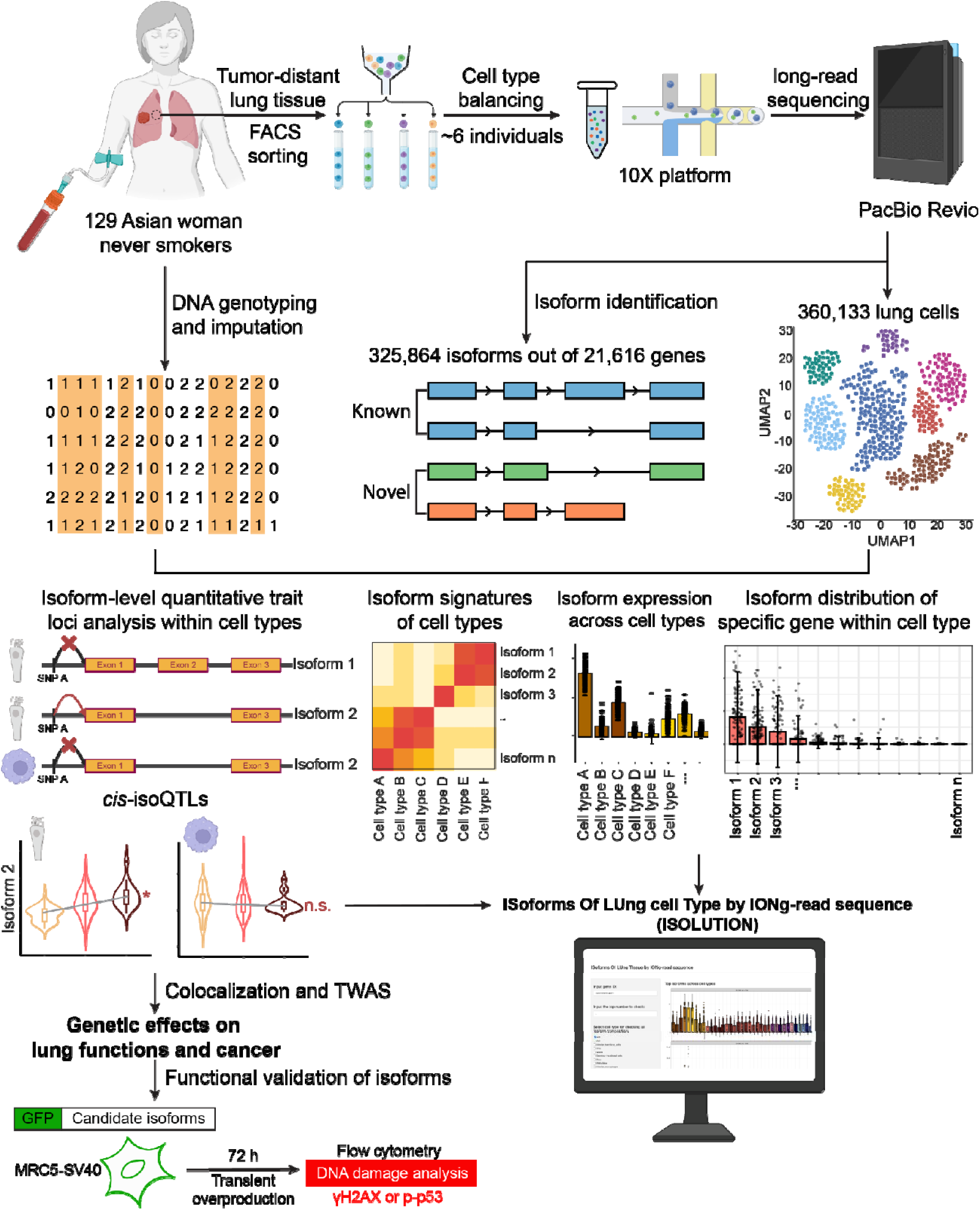
Schematic diagram of the study design. Workflow of single-cell long-read sequencing includes sample collection, cell type balancing with epithelial cell enrichment, single-cell capture, and long-read RNA-sequencing. DNA genotyping from matched blood samples and imputation were performed in parallel. Isoform identification and cell type annotation were followed by the main analyses, including cell type signature isoform detection, cell-type-specific isoQTL detection, and identification of target isoforms for lung function and cancer GWAS loci. The function of lung cancer susceptibility isoforms was experimentally validated by DNA damage assay. Open access website ISOLUTION (https://appshare.cancer.gov/ISOLUTION) features isoform structures, isoform expression across cell types, and isoQTL results from this study. FACS: fluorescence-activated cell sorting.

## Results

### Full-length transcriptome uncovers isoform diversity in 129 lung tissues

To profile full-length single-cell transcriptomes of lung tissues from 129 individuals (**Supplementary Table 1**), we adopted PacBio single-cell multiple amplicon sequencing (MAS-seq), incorporating cell type balancing and multiplexing (**Fig. 1**). Of note, we generated a cell-barcode-matched short-read-based dataset using the same cDNA libraries (Luong et al.^29^), which allowed systematic comparisons between two datatypes. Matching genotype data from blood samples was obtained using DNA genotyping followed by imputation (**Methods**). We employed cell type balancing to further reduce sampling bias and enrich epithelial cells, which include cells of lung cancer origin but are underrepresented in single-cell processing ^23,30^. For this, fluorescence-activated cell sorting (FACS) was used^31^ (**Methods**), followed by multiplexing of ∼6 individuals per single-cell batch (22 total batches).

In total, we identified 325,864 full-length isoforms from 129 lung tissues after quality control (QC)(**Methods, Fig. 2A**). These transcripts were defined in six structural categories (SQANTI3 ^32^): full-splice-match or FSM, incomplete-splice-match or ISM, novel-in-catalog or NIC, novel-not-in catalog or NNC, fusion gene, genomic, and antisense (**Fig. 2B**). Among these, the known isoforms (FSM) accounted for 82% of total full-length reads, indicating the total gene levels could be mainly represented by the known isoforms (**Fig. 2C**). The detected known isoforms accounted for 34.6% of the annotated transcripts (55,329) from GENCODE v32 (**Supplementary Fig. 1A**). Notably, 83% of the total isoforms were novel transcripts not annotated in GENCODE v32 (representing all the categories except FSM in **Fig. 2D**). The high proportion of novel isoforms is consistent with the findings in recent bulk-tissue long-read sequencing datasets (26-77%)^33,34^. For 20.0% (4,323/21,616) of genes, only one isoform was detected, which were mostly annotated isoforms (80% of 4,323 genes) (**Fig. 2E**). For 80% of 21,616 genes, multiple isoforms were detected, where the numbers of novel isoforms significantly correlated with those of known isoforms (correlation coefficient: 0.44, *p*-value < 2.2e-16), similar to previous studies ^6^. Leveraging our sample size, we surveyed the isoform distribution across 129 individuals (median 55,872 per individual). We observed known isoforms exhibited bimodal distributions, where a substantial proportion (7.8%) was widely expressed in all individuals. Novel isoforms were mainly specific to smaller subsets of individuals compared to the known ones (**Fig. 2F**).

**Figure 2.**
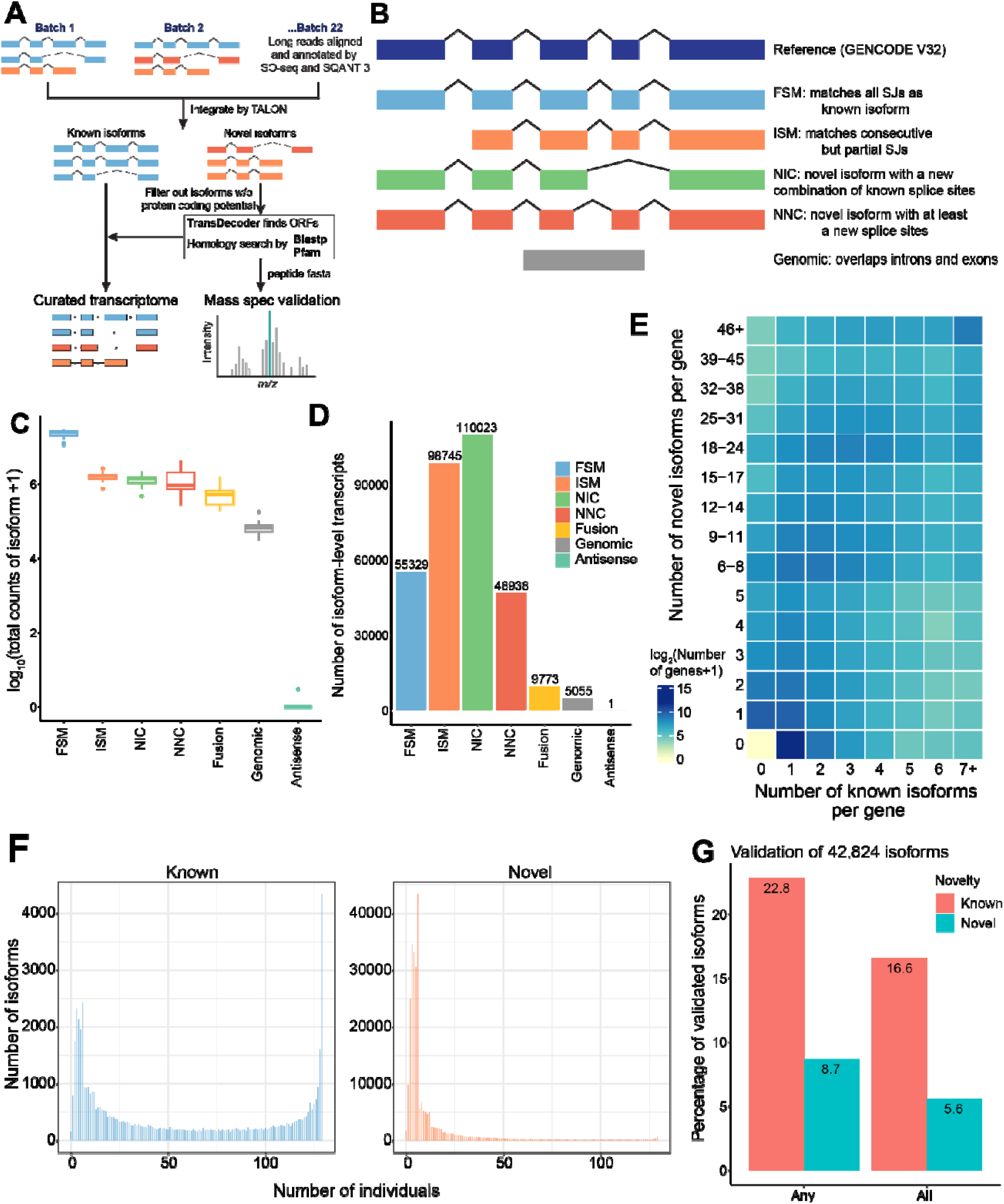
Identification and characterization of transcript isoforms in lung cells. **(A)** The workflow of transcriptome integration across single-cell batches and quality control (QC) for isoform curation. (**B)** Schematic plots of structural categories of isoforms according to SQANTI3. For an isoform not FSM or ISM, it is defined as NIC if including a novel splice junction(s) composed of known donor and acceptor sites, or NNC if including a novel splice junction(s) including at least one novel donor or acceptor site. (**C**) Abundance of isoforms from each structural category. Each data point represents the summed raw counts of all single-cell batches for each isoform that were log_10_ transformed after a pseudo-count of 1 was added. Center lines show the medians; the box indicates the middle of 50% of the data; whiskers extend 1.5 times the interquartile range from the 25th and 75th percentiles; outliers are represented by dots. (**D)** Numbers of post-QC isoforms from each structural category annotated according to SQANTI3. (**E)** Number of known and novel isoforms per gene. The color in the heat map shows the log_2_-transformed number of genes. **(F)** Distribution of known (top) or novel (bottom) isoforms detected across the number of individuals. (**G)** Percentage of validated isoforms detected in any or all 4 samples analyzed by mass-spectrometry. The colors show the novelty of isoforms.

To gain further insights into novel isoforms, we estimated an isoform’s potential to be a target of nonsense-mediated decay (NMD) based on the presence of a premature termination codon located >=50 nucleotides upstream of the last splicing junction. We found that the proportions of isoforms sensitive to NMD were higher in novel isoform categories than in known ones (3.4% in known or FSM vs 16.8% in all novel classes), where NIC (new combination of known splice sites) and fusion isoforms had the highest proportions (20.7% and 22.3%, respectively) (**Supplementary Fig. 1B**).

To validate the detected isoform diversity in lung cells, including the prevalence of novel and individual-specific isoforms, we took multi-layered approaches.

First, we sought replication of our isoforms in the GTEx custom full-length transcriptome using 14 types of bulk tissues including lung ^6^. Given a larger sample size and a higher total isoform number in our data, we replicated 35.8% (31,521) of the GTEx isoforms. Among the known isoforms, our data detected a higher proportion of annotated transcripts (34.6% in GENCODE v32 vs. 12% in GENCODE v26 ^6^). For novel isoforms, 49.8% of the replicated isoforms were annotated as novel in both datasets, including 729 novel isoforms validated by mass-spectrometry in GTEx.

Second, we further validated novel isoforms at the peptide level by liquid chromatography-tandem mass spectrometry (LC-MS/MS) proteome profiling of bulk lung tissues from a subset of our samples (n = 4). Of 42,824 qualifying isoforms (expressed in >20% individuals, **Methods**), 22.8% of known isoforms (5,495/24,127) and 8.7% of novel isoforms (1,639/18,697) were validated with matching peptides (**Supplementary Table 2, Methods**), which is consistent with proteomic validation of isoforms identified by bulk long-read RNA-seq (approximately 10% across tissue types)^6^. Major subsets of these mass-spec validated isoforms (73.1% and 63.8%) were detected in all 4 samples (**Fig. 2G**).

Third, we cross-validated splice junctions of our isoforms in the barcode-matched single-cell short-read dataset with a higher sequencing depth (**Methods**). Among 346,748 splice junction sites, 31.2% were validated in our short-read data, including 18.7% of novel junctions. Conversely, 56.4% of splice junctions detected in short-read data by Leafcutter were matched to a full-length isoform, allowing the functional interpretation of the junction in the isoform context (**Supplementary Fig. 1C**).

Fourth, we performed additional sequencing to assess whether the observed individual-specific isoforms are due to the sequencing coverage of our dataset (2 SMRT cells per batch). For this, we sequenced two representative libraries using 4 SMRT cells on PacBio Revio. Saturation analysis indicated that the detection of known genes and isoforms was saturated with a limited increase beyond 2 SMRT cells (vertical dashed lines in **Supplementary Fig. 2A** left and middle panels), indicating a sufficient depth at this cutoff. However, there was no clear saturation for total isoform detection due to the contribution of novel isoforms (**Supplementary Fig. 2B** right panel). Nevertheless, doubling of sequencing depth from 2 to 4 SMRT cells only increased the number of isoforms by 13.0% and 14.6%, indicating an attenuated effect on isoform detection (**Methods, Supplementary Fig. 2B**). Notably, 17.0% and 18.0% of these isoforms were from those exclusively observed in one of the 22 batches at the 2 SMRT cell coverage (**Supplementary Fig. 2C**). These data indicated that the observed individual-specific isoforms in our dataset might be detected and validated in multiple individuals in different batches with increased sequencing depth. At single-cell levels, we observed 37% and 35% increase in the per-cell isoform detection by doubling sequencing depth (**Supplementary Fig. 2D**). A steeper increase in per-cell isoform detection compared to batch level suggested that pseudo-bulk approaches will be needed for cell-type-specific analyses.

Collectively, the validation supported the reliability of our full-length transcriptome of normal lung tissues that could be used for direct assessment of cell-type-specific isoform-level expression.

### Isoform-level lung cell type signatures are more specific than gene-level

Our barcode-matched dataset enabled us to investigate whether isoform expression could provide additional cell-type-level information not captured in total expression. To characterize cell type signatures of isoform expression, we first annotated cell types using the gene-level expression matrix to align with the established markers of lung cell types ^35^ (**Methods, Supplementary Fig. 3**). In total, we identified 37 lung cell types in four major lung cell categories: epithelial, immune, endothelial, and stromal cells (**Fig. 3A and Supplementary Fig. 4A**). Reflecting our design of epithelial cell enrichment and cell type balancing, epithelial cells accounted for 45% of total cells (**Fig. 3B**). Compared to the independent cell type annotation of our short-read dataset ^29^, we found a cell-to-cell annotation match for 84.3% cells, with few rare cell subtypes missing in the long-read data (**Supplementary Fig. 4B**).

**Figure 3.**
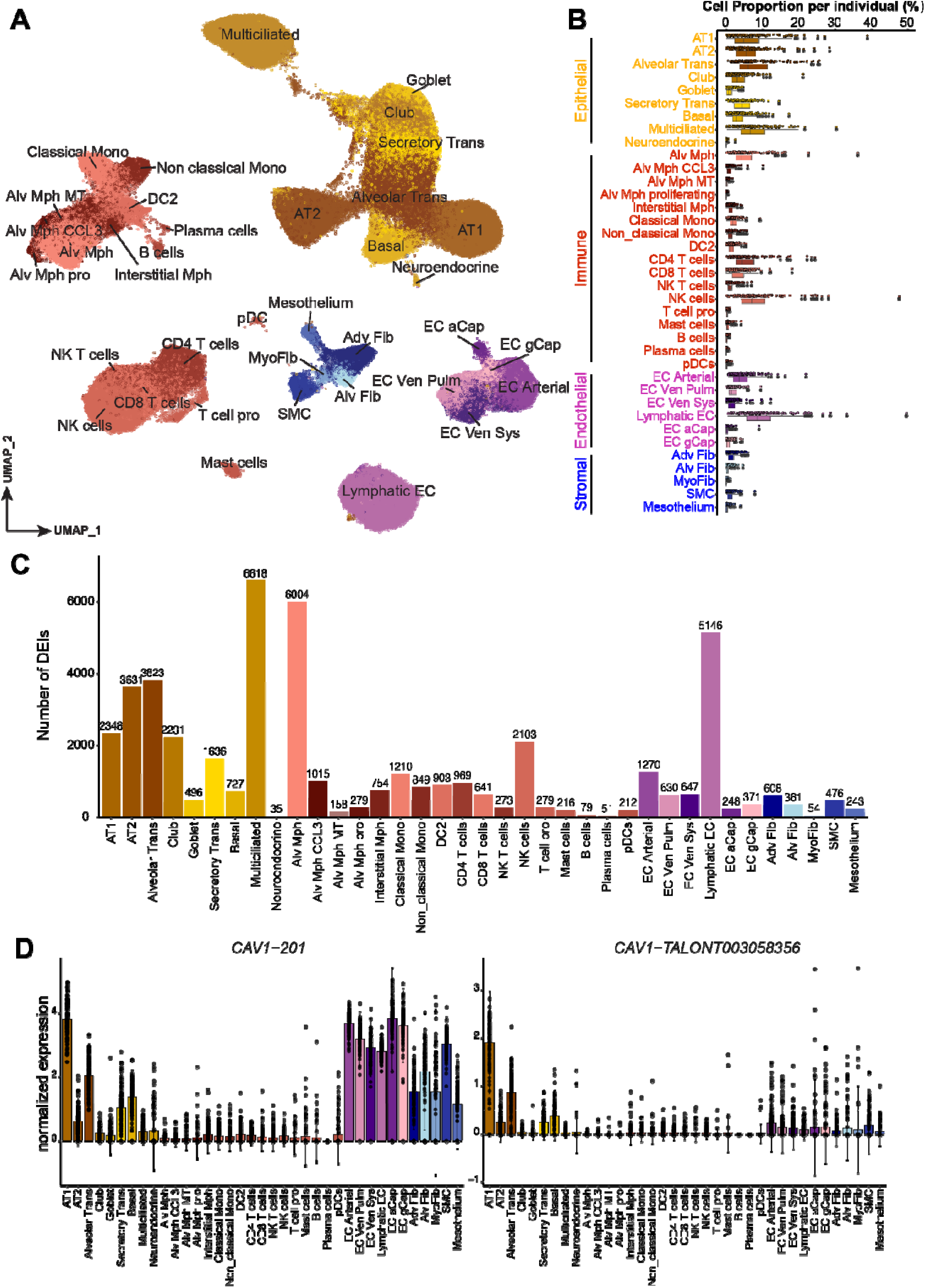
Isoform-level lung cell atlas and cell type signature isoforms. **(A)** Uniform manifold approximation and projection (UMAP) plot depicts 37 lung cell types from 129 individuals. (**B**) The box plots with scattered points show the proportion of each lung cell type in each individual. Center lines show the medians; the box indicates the middle of 50% of the data; whiskers extend 1.5 times the interquartile range from the 25th and 75th percentiles; outliers are represented by dots. Each point on the top of the box represents the proportion of cell type within an individual. (**C**) Number of differentially expressed isoforms (DEIs) across 37 lung cell types. DEIs are defined as isoforms with log2FC > 1 and adjusted *p*-value < 0.05 in differential expression analysis. (**D**) Normalized expressions of *CAV1-201* and *CAV1-TALONT003058356* at the individual level across lung cell types. Each dot represents the isoform expression of each individual within each cell type. The error bars show the mean of normalized expression ± standard deviation. Alveolar Trans: alveolar transitional cells; Secretory Trans: secretory transitional cells; Alv Mph: alveolar macrophages; Mono: monocytes; pDC: plasmacytoid dendritic cells; EC: endothelial cells; aCap: capillary aerocytes; gCap: general capillary; EC Ven Pulm: pulmonary venous ECs; EC Ven Sys: systemic venous ECs; Adv Fib: adventitial fibroblast; Alv Fib: alveolar fibroblast; SMC: smooth muscle cell; MyoFib: myofibroblast.

To detect cell type signatures of isoform expression, differential expression analysis was performed within each cell type (**Methods**). To address the data sparsity at the single-cell level (**Supplementary Fig. 5A**) and exclude batch-specific isoforms (**Fig. 2F**), we pseudo-bulked isoform expression into individual levels and retained relevant abundant isoforms for each cell type (**Methods**). We nominated differentially expressed isoforms (DEIs, defined as isoforms with log_2_(fold change) > 1 and adjusted *p-*value < 0.05) of each cell type against all the other cell types. Out of 42,824 tested isoforms, we identified 19,595 DEIs from 10,135 genes across 37 lung cell types (**Fig. 3C**), with 5,719 (29.2%) being novel isoforms. Nearly all of these 10,135 genes (89.9%) were also differentially expressed in the same cell type at the gene level (adjusted *p-*value < 0.05, **Methods**). Consistently, 50.8% of these DEIs were the most-abundant isoforms of their corresponding genes, suggesting that DEIs captured the representative isoforms of cell-type signature genes. Similarly, the DEIs were well-represented in gene-level cell type markers that we used for cell type annotation, where the most abundant isoforms from 74 (77.9%) markers were detected as DEIs, and 89.2% of them were known isoforms.

To assess the cell-type specificity of isoform signatures, we first compared the fold changes of cell-type-specific expression between isoform- and gene-levels. We found that 13,163 DEIs (67%) displayed a larger fold change than the corresponding genes in at least one cell type (**Supplementary Fig. 6A**). These data suggested that a substantial subset of isoforms are more cell-type-specific than their corresponding genes, which could be leveraged in cell type annotation as isoform-level markers. For example, caveolin-1(*CAV1*) is commonly used as an AT1 cell marker ^36^ but is also expressed by multiple other cell types within and outside the epithelial cell group at the gene level (**Supplementary Fig. 6B**). For the most abundant isoform, 201, this pattern also persisted (**Fig. 3D and Supplementary Fig. 6B**). Notably, we detected a novel isoform of *CAV1*, *TALONT003058356*, as a significant DEI, which was expressed by 97.7% of individuals and demonstrated a higher specificity to AT1 (**Fig. 3D and Supplementary Fig. 6B**). This specificity was maintained at a higher sequencing coverage, indicating that the isoform specificity could not be simply attributed to the sparsity or lower expression but are more likely to reflect biological distinction (**Supplementary Fig. 7A-B**).

To further investigate whether isoform signatures contribute to cell type clustering and annotation, we compared clustering performance between highly variable isoform and gene signatures. Benchmarking of clustering performance for cell type annotation demonstrated that using top-ranking gene-level features outperformed an equal number of top-ranking isoforms in dataset-wide comparisons. However, top-ranking isoform features outperformed their matched genes, by gaining additional isoform information for the same set of genes (**Supplementary Fig. 8A-D**). Furthermore, at the cell-type-level comparisons, 8 of 37 cell types displayed a better performance of isoform-level features (**Supplementary Fig. 8E**). These data suggested that isoform-level information for a subset of cell types could contribute to cell clustering and cell type annotation even when using gene-level ground truth based on conventional marker genes.

Collectively, our data demonstrated that cell-type-specific isoforms could capture distinct cell-type differences beyond average gene expression in a subset of lung cell types.

### Single-cell lung isoQTLs highlight cell-type-specific isoform regulation

Leveraging our sample size of 129 individuals from a similar genetic background, we performed isoQTL mapping on 33 qualifying cell types (**Methods**). To address the data sparsity, we pseudo-bulked the single-cell counts at the individual level using the same criteria as differential expression analyses (**Fig. 4A**).

**Figure 4.**
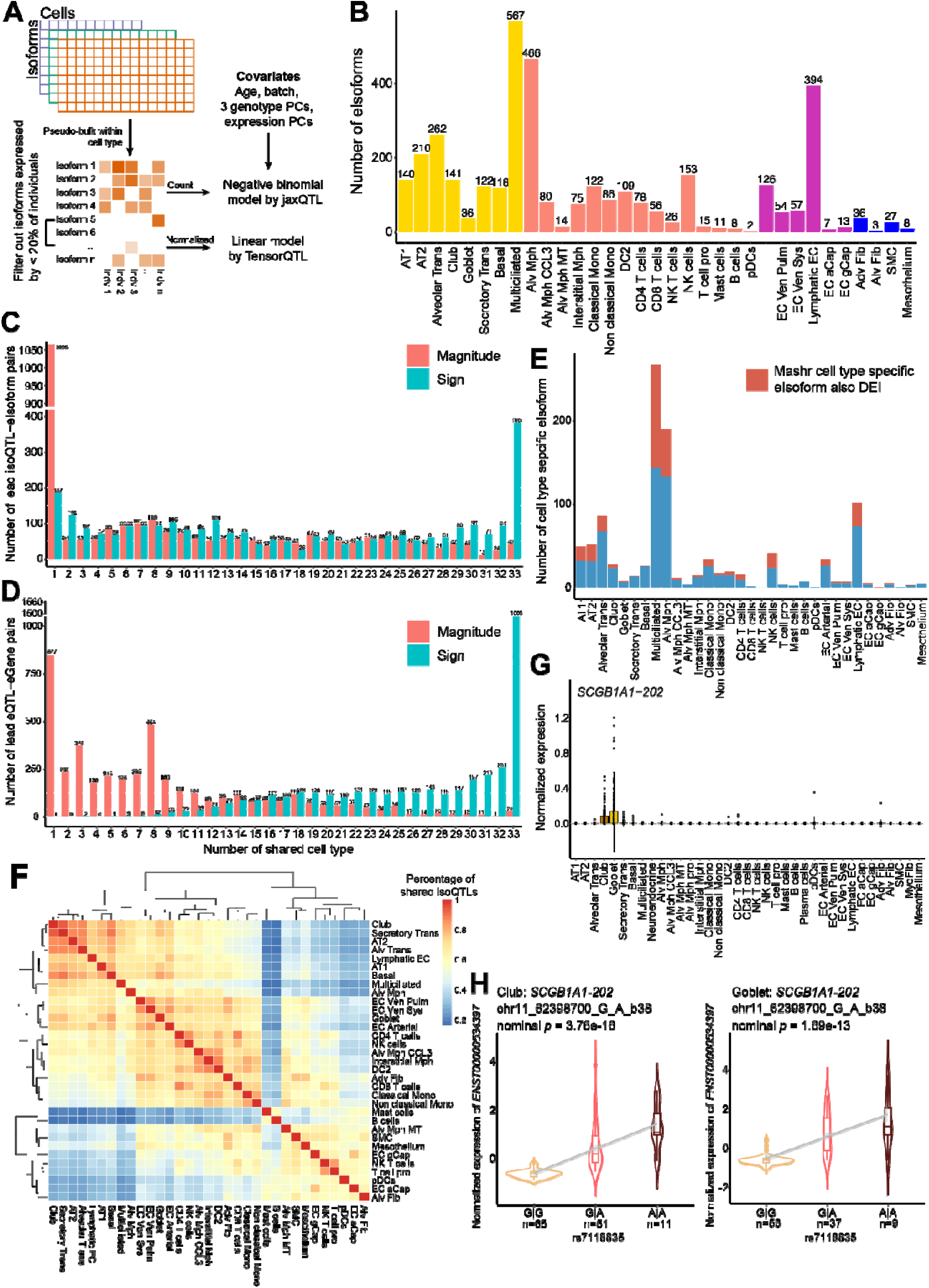
isoQTLs of lung show cell-type specificity and lineage-specific sharing. (**A**) Schematic of pseudo-bulk isoQTL mapping process. Isoform filtering criteria and covariates for negative binomial regression are shown. (**B**) Number of eIsoforms identified by isoQTL analysis across 33 cell types. The bars are colored by cell categories (yellow for epithelial, red for immune, purple for endothelial, and blue for stromal). (**C**) Number of lead isoQTLs shared based on magnitude or sign across cell types after Mashr harmonization. Sharing magnitude is defined as the effect size of an isoQTL being within a factor of two of the top isoQTL (isoQTLs with the maximum absolute value of effect size) across the indicated number of cell types. Sharing sign is defined as an isoQTL sharing the same allelic direction with the top isoQTL across the cell types. (**D**) Number of lead eQTLs ^29^ shared based on magnitude or sign across cell types after Mashr harmonization as described in **Methods.** (**E**) Number of cell-type-specific eIsoforms across 33 cell types. The proportions also detected as DEIs for the same cell type are colored in red. (**F**) Pairwise isoQTL sharing based on magnitude among cell types. In pairwise comparison, isoQTLs significant in at least one cell type were included. The color in the heat map represents the percentage of shared isoQTLs in the paired cell types. (**G**) Normalized expressions of *SCGB1A1-202* at the individual level across lung cell types. Each dot represents the isoform expression of each individual within each cell type. The error bars show the mean of normalized expression ± standard deviation. (**H**) Association between normalized expression of *SCGB1A1-202* and the genotype of lead isoQTL rs7118835, in club (left) and goblet cells (right). The lead isoQTL is shown at the top with chromosome, position, reference allele and alternative allele based on hg38 assembly. Center lines show the medians; the box indicates the middle of 50% of the data; whiskers extend 1.5 times the interquartile range from the 25th and 75th percentiles; outliers are represented by dots; density of normalized expression is represented by the width of violin shape.

We applied a negative binomial (NB) regression model for isoQTL mapping using jaxQTL^37^, which was shown to be powerful for sparse datasets and well calibrated on permuted data (**Supplementary Fig. 9** and details in **Methods**). Benchmarking between this NB model by jaxQTL and a linear model using TensorQTL revealed that the NB model outperformed the linear model. Namely NB model identified 27.5% higher eIsoform numbers (isoforms with at least one significantly associated variant in at least one cell type), especially among lower-expressed isoforms (**Supplementary Fig. 10C**). The NB model also replicated 78.9% of the eIsoforms identified by the linear model with a consistent effect direction (**Supplementary Fig. 10A-B**). Across 33 cell types, we identified 2,016 unique eIsoforms from 1,562 unique isoGenes (genes with at least one eIsoforms in at least one cell type) (**Supplementary Table 3**). Among 2,016 eIsoforms, 428 isoforms (21.2%) were validated by our mass-spectrometry data with 14.3% being novel. isoQTL detection was positively correlated with the numbers of tested isoforms (Pearson *r^2^* = 0.75, **Supplementary Fig. 11A-B**) as well as the total numbers of cells in that cell type (Pearson *r^2^*= 0.61, **Supplementary Fig. 11C**). The numbers of cells and tested isoforms contributed to the power of isoQTL detection more than the numbers of individuals in our sample size (likelihood ratio test; number of cells: *p-*value = 4.76e-05, number of individuals: *p-*value = 0.2739, **Supplementary Fig. 11D**). These data suggested that increasing cell numbers by reducing the multiplexing complexity and/or by increasing sequencing coverage could improve the power of isoQTL detection in a given sample size.

To evaluate the specificity and sharing pattern of isoQTLs across lung cell types, we used mash ^38^ to harmonize the allelic effect sizes and account for differences in power. Of 2,842 lead isoQTL-eIsoform pairs from all cell types, 37.5% (1,065 pairs) were specific to one cell type based on both allelic effect direction and size (less than 2-fold) (**Fig. 4C**). Notably, isoQTLs showed a higher cell-type specificity than eQTLs from the barcode-matched short-read dataset ^29^, where 19.4% (4,372 pairs) were specific to a single cell type, based on a similar harmonization by mash (*p-*value = 3.93e-65 by χ2 test, **Fig. 4D**). Among the cell-type-specific isoQTL-eIsoform pairs, 31.64% were also a DEI in the matching cell type, suggesting that cell-type-specific expression drives specific isoQTL detection for this subset (**Fig. 4E**). For example, *PPIL6-207* (the most abundant isoform of Peptidylprolyl Isomerase Like 6; **Supplementary Fig. 12A-B**) was a DEI only in multiciliated cells (**Supplementary Fig. 12C**) and identified as an eIsoform only in this cell type for the lead isoQTL, rs1258822 (**Supplementary Fig. 12D**). While more isoQTLs shared the same allelic direction (13.5% of isoQTL-eIsoform pairs shared directions in all 33 cell types, **Fig. 4C**), we also observed isoQTLs displaying opposite allelic directions between different cell types. For example, *NAPRT-204* was widely expressed by multiple cell types, but an isoQTL, rs896962, displayed opposite allelic effects on *NAPRT-204* expression between alveolar macrophages and multiciliated cells (**Supplementary Fig. 13**).

Beside cell-type-specific isoQTLs, we also observed lineage-preferential sharing of isoQTLs consistent with isoform expression patterns. Pair-wise isoQTL effect sharing across cell types indicated a higher level of sharing among cell types of the same category beyond similarity in cell cluster sizes, suggesting isoQTL effect reflects biological distinctions of the cell type sub-groups (**Fig. 4F**). For example, isoform 202 of *SCGB1A1*, a marker gene of secretory epithelial cell lineage, was specifically expressed in club and goblet cells and identified as eIsoforms only in these two secretory cell types, where the allelic effect of lead variant, rs7118835, was shared in the same direction and similar sizes (**Fig. 4G-H**).

In summary, we detected isoQTLs from 33 lung cell types and observed higher cell-type specificity of genetic regulation of isoform levels compared to gene level.

### isoQTLs reveal genetic regulation of isoforms distinct from average expression

To investigate whether our isoQTLs represent unique genetic regulation of isoforms not captured in previous datasets, we first compared our isoGenes with sGenes (genes with at least one significant sQTLs) of lung tissues. While 41.1% of isoGenes were replicated at the gene level as sGenes in GTEx (v8) lung tissues (n = 515), 921 isoGenes were new to our dataset (**Fig. 5A**). Among them, 54.7% were only identified in a single cell type, suggesting that cell-type specificity not captured in bulk tissues might explain this subset. Moreover, 29.4% of the 921 isoGenes were for novel transcript isoforms, suggesting that detection of novel isoforms helped identify additional isoGenes not captured in short-read-based sQTLs (**Fig. 5B**). Additionally, we asked whether some of these 921 isoGenes were not detected in GTEx (∼85.3% European) due to allele frequency differences with East Asian populations (EAS). Among 1,319 lead variants of 921 isoGenes,11.9% (158/1,319) showed minor allele frequency (MAF) < 0.01 in EUR (1000 Genomes Project Phase III), which might not have been tested in the GTEx sQTL analysis, whereas this proportion was lower (7.7%, 103/1,334) in isoGenes replicated in the GTEx sGenes (*p* = 0.0002987, χ2 test) (**Supplementary Table 4**). Moreover, for the isoGenes that are matched to GTEx sGenes, 74.7% (847/1,134) of independent lead isoQTLs were not found in GTEx sQTLs and 50.5% (428/847) among them were in low LD (R^2^ < 0.1) with the GTEx lead SNP or monoallelic in the EUR population (**Methods**). These data indicated that our dataset-specific isoGene findings compared to mainly-European bulk sQTLs are driven by cell-type-specific, novel, or East Asian-specific isoQTLs.

**Figure 5.**
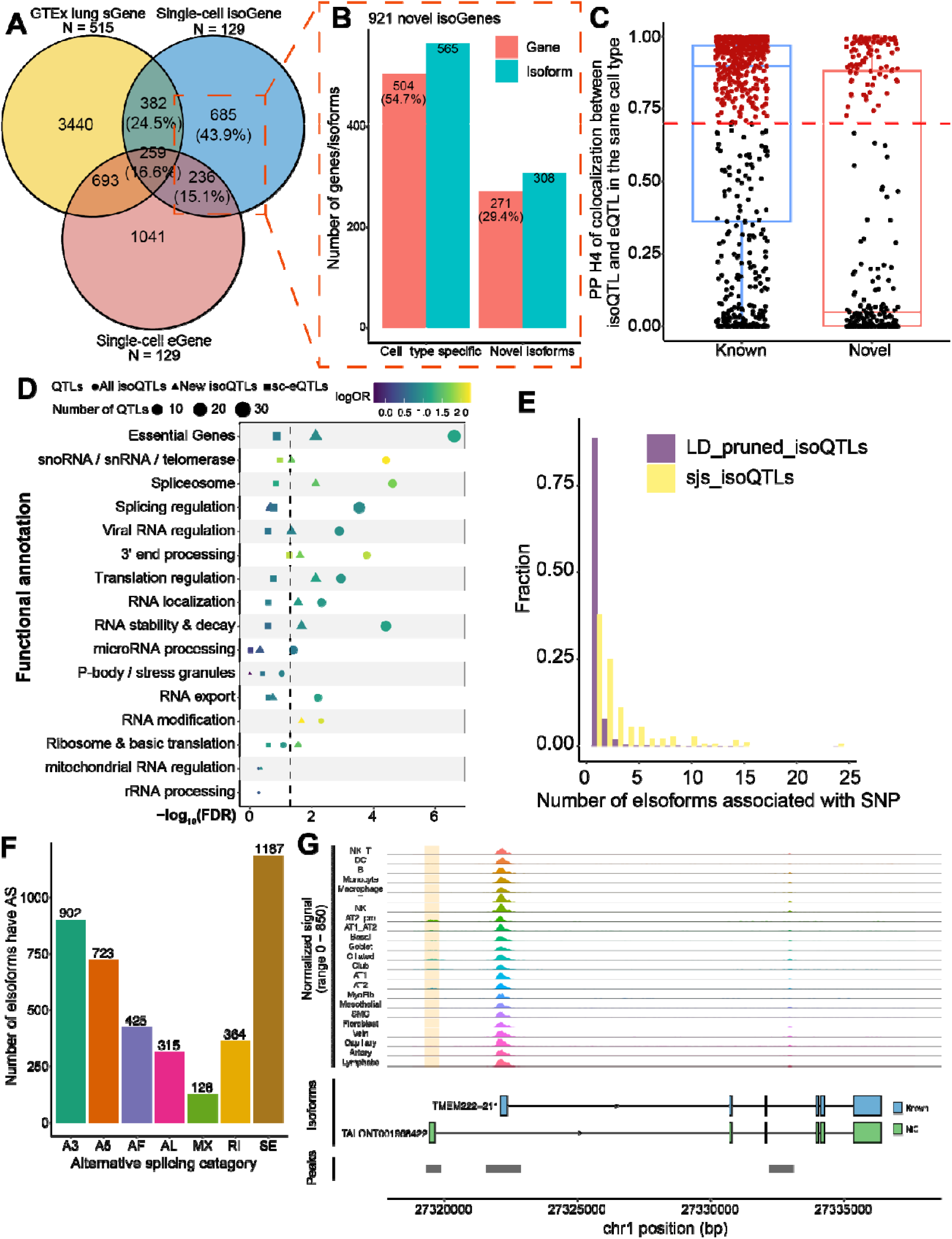
Functional annotation of isoQTLs. (**A**) Venn diagram depicts the number of gene-level overlaps among isoGenes of this study, sGenes of GTEx lung tissues (version 8), and eGenes of barcode-matched short-read single-cell eQTL data. Percentages of isoGenes are shown in parentheses. (**B**) Number of genes or isoforms that are cell-type-specific or novel (not annotated) out of 921 dataset-specific isoGenes not overlapping sGenes in GTEx lung dataset. (**C**) Posterior probabilities of colocalization between an eIsoform and a matched eGene in the same cell type. Each dot represents an eIsoform, and those above the red dashed line (PPH4 = 0.7) are colored in red and considered as significantly colocalized eIsoforms. Center lines show the medians; the boxes are colored for known or novel isoform groups, indicating the middle of 50% of data; whiskers extend 1.5 times the interquartile range from the 25th and 75th percentiles. (**D**) Enrichment of LD-pruned isoQTLs in the binding motifs of RNA-binding proteins (RBPs). The size of dots represents the number of isoQTLs located in the motifs, and the color shows the log(odds ratio). The motifs are annotated according to the functions of their RBPs. The shape of dots represents the set of QTLs, where circles represent all LD-pruned isoQTLs, triangles represent the LD-pruned isoQTLs associated with dataset-specific isoGenes (i.e., 921 isoGenes not identified as a sGene in GTEx lung), and squares represent all LD-pruned single-cell short-read eQTL. The dashed line indicates the FDR of 0.05. (**E**) Distribution of the numbers of eIsoforms associated with all LD-pruned isoQTLs or splice junction site-overlapping isoQTLs (sjs-isoQTLs). (**F**) Number of alternative splicing (AS) events identified by SUPPA2 for 2,016 eIsoforms. Alternative 3′ (A3) or 5′ splice site (A5), alternative first (AF), or last exon (AL), retained intron (RI), skipped exon (SE) and mutually exclusive exons (MX). (**G**) Single-cell ATAC-seq peak signals of canonical promoter and potential alternative promoter for *TMEM222* in normal lung tissues from Long *et al*. The potential alternative promoter peak (highlighted in beige) aligns with the transcription start site of the novel eIsoform (estimated as alternative first exon and colored in green).

Next we asked whether isoQTLs represent genetic signals distinct from average gene expression captured by eQTLs. Leveraging the barcode-matched short-read data ^29^, we first observed a relatively low level of sharing between single-cell isoGenes and eGenes (31.7% of isoGenes are also eGenes) (**Fig. 5A**). Among the overlaps, 59.2% of isoGenes were detected in the same cell types. isoQTL-eQTL colocalization analysis of this subset demonstrated that 53.6% isoQTLs (250/466 eIsoforms) colocalized with eQTLs (194 eGenes) (posterior probability of the hypothesis H4 by coloc ^39^, PPH4 > 0.7) (**Fig. 5C, Supplementary Table 5**). Of these colocalized eIsoforms, 81.2% were the known isoforms in GENCODE v32, and 58.8% were the most abundant isoforms of the corresponding genes, suggesting that they mainly contribute to the total gene levels and potentially contribute to eQTL signals. However, 46.4% of the tested isoQTLs did not colocalize with eQTLs, suggesting genetic effects independent from gene expression regulation. Notably, a higher proportion of isoQTLs among novel isoforms did not colocalize with eQTL compared to that of known isoforms.

To understand the functional basis of unique isoQTL findings, we assessed their enrichment in motifs of RNA-binding proteins (RBPs) grouped into functional categories. For this, we used the RBP motifs identified by enhanced crosslinking and immunoprecipitation followed by sequencing (eCLIP-seq) in the ENCODE project ^40^. We found that isoQTLs were significantly enriched in the functional categories relevant to splicing regulation and other post-transcriptional processing, including “spliceosome”, “splicing regulation”, “3’ end processing” and “RNA stability & decay” (**Fig. 5D**). For example, seven LD-pruned isoQTLs overlapped motifs of RNA helicase and spliceosomal protein, AQR, suggesting their potential regulatory effects on splicing by affecting intron-binding of spliceosomes ^41^ (**Supplementary Fig. 14**). In addition, the isoQTLs for the 921 isoGenes not found in the GTEx sGenes displayed a similar enrichment pattern, further validating the robustness of our dataset-specific isoQTLs. In contrast, eQTLs from the barcode-matched short-read data showed no significant enrichment in any of the RBP motif categories (**Fig. 5D**). These data indicated post-transcriptional regulatory mechanisms underlying isoQTLs that are distinct from eQTLs.

We then asked if isoQTLs affect conserved splicing junction sites (sjs), which might have a direct functional impact rather than tag the effect of functional variants in LD. For this, we identified 60 sjs-isoQTLs located in the first and last 2bp of an intron of eIsoforms. We found that 64.5% of these sjs-isoQTLs were associated with more than one eIsoforms, a significant enrichment compared to 11.3% LD-pruned isoQTLs (**Fig. 5E**; χ2 test *p* = 1.282e-10). These data indicated that sjs-isoQTLs were more likely to be associated with multiple eIsoforms, potentially due to the reciprocal effect on multiple alternative isoforms sharing the same splice junction. For example, a COVID-19 severity-associated splice acceptor variant ^42^, rs10774671, in *OAS1 ^43,44^* was identified as a sjs-isoQTL in our dataset, where a reciprocal allelic effect of three isoforms was more pronounced in lung alveolar macrophages (**Supplementary Fig. 15**). Our sjs-isoQTL also replicated an EAS-biased sQTL, rs11064437, in *SPSB2,* reported in a blood sQTL study using short-read data ^14^. We demonstrated that the affected novel splice junction can now be mapped to a novel isoform, *TALONT0006364782*, providing a better understanding of the reciprocal isoform regulation of this gene (**Supplementary Fig. 16**).

Beyond reciprocal splicing, mechanisms such as intron retention (RI) and alternative first exon splicing (AF) were inferred from isoQTL functional classes, informed by full-length isoforms (**Fig. 5F**) using SUPPA2. Namely, 364 eIsoforms exhibited intron retention with significant enrichment in NMD sensitivity (Fisher’s exact test *p* = 0.001287, odds ratio: 2.24), suggesting the potential connection to RNA degradation ^45^. In addition, AF events indicated alternative transcription start sites (TSSs), potentially attributed to cell-type-specific alternative promoter usage. By integrating our single-cell ATAC-seq data of similarly processed lung tissues^43^, we found that TSS of 142 eIsoforms with AF events overlapped cell-type-specific non-canonical promoter ATAC peaks (**Methods**). Moreover, 94% of these eIsoform-peak pairs were detected in a matched cell type. For example, we detected an epithelial cell-specific peak near the TSS of a novel isoform of *TMEM222* (TALONT001958422), which was upstream of the multi-cell type promoter peak for the main isoform, *211* (**Fig. 5G**). The data suggested cell-type-specific isoform regulation by potential alternative promoter usage. Together these findings supported the unique functional contribution of isoQTLs through alternative mechanisms beyond regulating total gene expression levels.

### GWAS integration highlights new target genes in lung epithelial cells

Having established distinct genetic regulation underlying isoQTLs, we investigated how they contribute to GWAS signals relevant to lung tissues. For this, we first performed colocalization of isoQTL with lung cancer GWAS in different histological types (overall lung cancer, LUAD, lung squamous cell carcinoma or LUSC, small cell lung cancer or SCLC) from diverse populations ^16,17^ and multi-ancestry GWAS of two lung function traits (forced expiratory volume in 1 second or FEV_1_ and FEV_1_ over forced vital capacity or FEV_1_/FVC) ^26^(**Fig. 6A**). Among 48 lung cancer-related GWAS loci, we observed colocalization (PP.H4 > 0.7) for 8 eIsoforms from 6 loci (**Fig. 6B, Supplementary Table 6**). These included 5 isoforms previously not colocalized with lung cancer at the gene level in bulk tissues: *HSPA1B* in 6q21.32, *HLA-A* in 6q21.33, *RSPH4A* in 6q22.1, *CHMP3* in 2p11.3, and *PPIL6* in 6q21 ^16,17^ (**Fig. 6B, Supplementary Table 6**). Among 346 lung function loci, we observed colocalization for 17 eIsoforms from 12 FEV_1_ loci and 27 eIsoforms from 18 FEV_1_/FVC loci (23 lead SNPs) (**Fig. 6B**, **Supplementary Tables 7-8**). Among them, 28 isoforms were previously not colocalized at the gene level using bulk tissue. These included *CTSS-TALONT003792695* in alveolar macrophages for FEV_1_, *RUNX1-202* in alveolar transitional cells for FEV_1_/FVC, and *EIF3E-211* in alveolar macrophages for both traits. Overall, among 49 eIsoforms colocalized with lung-relevant GWAS signals (**Fig. 6C**), 69.4% were from previously non-colocalized genes^16,17,26^.

**Figure 6.**
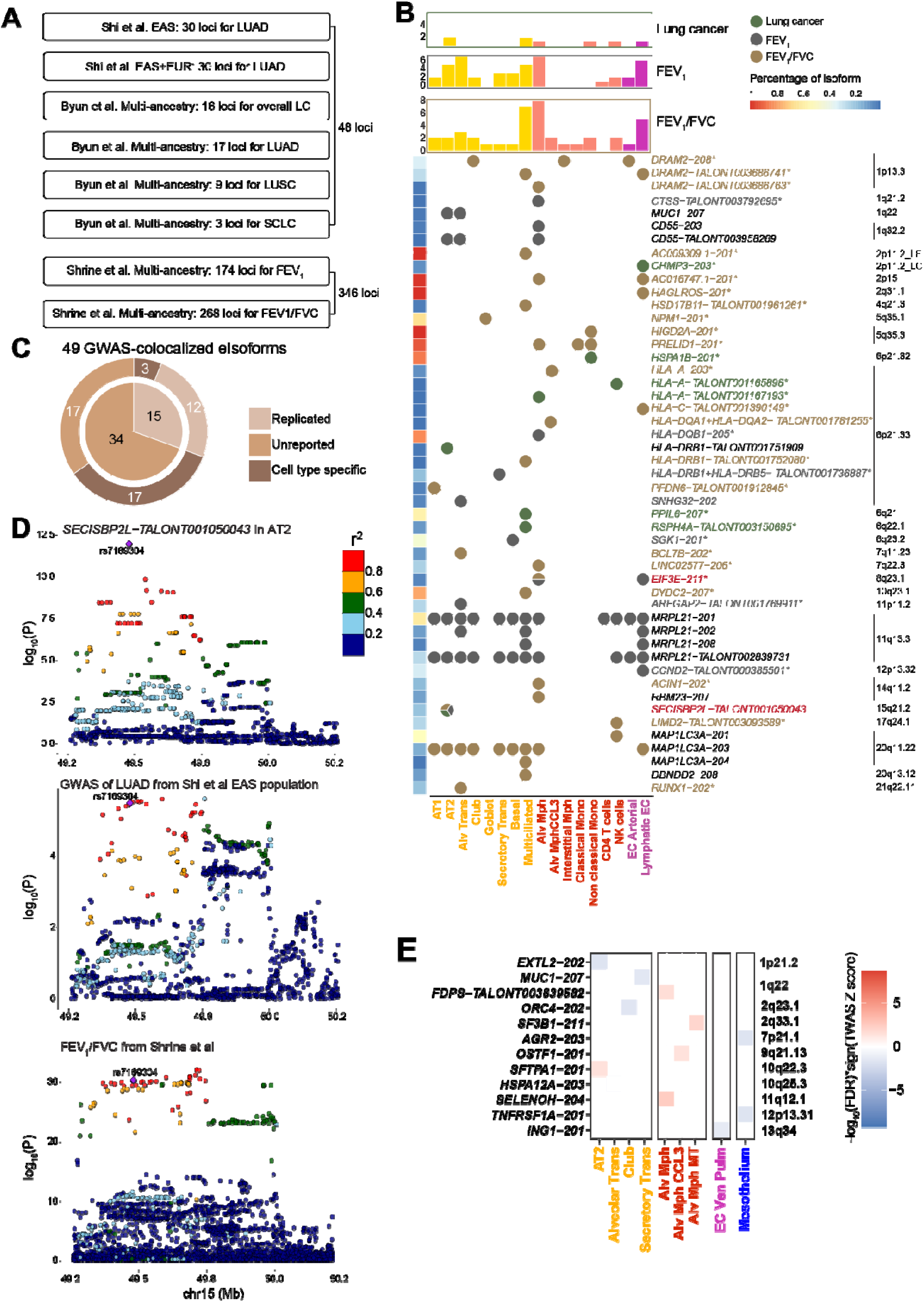
GWAS integration with lung function and cancer subtypes. (**A**) GWAS summary statistics of lung cancer subtypes and lung function traits used for colocalization. (**B**) Colocalization of 49 eIsoforms for three traits (i.e., lung cancer, FEV_1_ and FEV_1_/FVC) across 17 lung cell types. The color of the circles indicates the associated traits (dark green for lung cancer, dark grey for FEV_1_ and bronze for FEV_1_/FVC). Colocalized eIsoforms from genes not previously reported by bulk lung tissue colocalization are shown with an asterisk, whereas the color shows the specific trait. The bar plots at the top panel show the number of colocalized eIsoforms across cell types. The bar is colored by cell categories (yellow for epithelial, red for immune, and purple for endothelial). (**C**) Pie chart presents the fractions of GWAS-colocalized isoforms from replicated or unreported genes. The outer circle presents the cell type specificity of isoQTL-eIsoform pairs determined by Mashr harmonized effect size magnitude across cell types. (**D**) isoQTL colocalization with lung adenocarcinoma (LUAD from Shi et al EAS population) and FEV1/FVC at the *SECISBP2L* locus in AT2 cells. SNPs are color-coded based on the LD R^2^ (1000 Genomes, EAS, phase 3) with the lead isoQTL of *TALONT001050043* (purple diamond). (**E**) Summary of TWAS results in loci not reaching genome-wide significance by LUAD GWAS of East Asian population. The color in the heat map shows the signed -log_10_FDR from TWAS (red: increased expression correlated with risk, blue: decreased expression correlated with risk). LUAD: lung adenocarcinoma, LC: lung cancer, LUSC: lung squamous carcinoma, SCLC: small cell lung carcinoma, LF: lung function.

Collectively, more than half of the colocalized eIsoforms were either cell-type-specific (by mash-adjusted effect size) or DEIs, indicating substantial contribution of cell-type specificity to the new findings (**Fig. 6C**). Most of the colocalization events were observed in alveolar macrophages, multiciliated, alveolar transitional, lymphatic EC, and AT2 (**Fig. 6B)**. These cell types have relevance to lung function, especially the pathogenesis of COPD ^30,46,47^ as well as lung cancer. Among them, *SECISBP2L-TALONT001050043*, an AT2-specific eIsoform, was colocalized with LUAD and two lung function traits in AT2 cells (**Fig. 6D**), which are progenitor cells for alveolar regeneration, and a LUAD origin. Consistent with previous genetic correlation and causal effect of reduced FEV_1_ and/or FEV_1_/FVC with lung cancer risk^24^, lower expression of *SECISBP2L-TALONT001050043* in AT2 cells were correlated with impaired lung functions (both FEV_1_/FVC and FEV_1_) and increased risk of lung cancer.

Notably, multiciliated cells have been implicated in lung function ^47,48^ but an underappreciated cell type in the lung cancer context. Two lung cancer susceptibility isoforms from unreported genes, *PPIL6-207* and *RSPH4A-TALONT003150695,* were multiciliated-specific eIsoforms. Among them, *RSPH4A* encodes a radial spoke head protein of cilia, relevant to respiratory-tract mucociliary clearance ^49^, and high-penetrance germline mutations in this gene were found in primary ciliary dyskinesia ^50^.

To improve power in susceptibility isoform detection using ancestry-matched GWAS data, we next performed a transcriptome-wide association study (TWAS) for LUAD from East Asian populations ^17^. In total, we identified 74 significant isoforms from 23 loci (**Supplementary Table 9**). Of note, 47.8% of the TWAS loci (11 loci with 12 isoforms) were new signals outside the significant GWAS loci, nominating new susceptibility genes at the gene level: *EXTL2, MUC1, FDPS, ORC4, SF3B1, AGR2, OSTF1, SFTPA1, HSPA12A, SLELNOH*, and *ING1* (**Fig. 6A and 6E**). Consistently, a large proportion of the TWAS isoforms (54 isoforms, 73%) were from these new loci or unreported genes from the known GWAS loci by previous TWAS using bulk lung tissue eQTL/sQTL ^17,51–53^. Among the previously reported susceptibility genes, *HLA-A* and *SECISBP2L* were consistently identified by both colocalization and TWAS of LUAD, albeit for different isoforms and/or cell types (**Supplementary Fig. 17B and Fig. 6B**).

TWAS implicated 26 lung cell types from all four categories for LUAD susceptibility. Of note, 82.4% (71% when excluding HLA genes) of TWAS isoforms were from epithelial and immune cell categories, consistent with previous implications of these cell categories in lung cancer ^31^. Similar to colocalization, 23% of TWAS isoforms were from DEIs of the matching cell types (**Supplementary Fig. 17C**). In addition, we noted isoforms of epithelial-lineage-specific genes, such as *MUC1-207* (epithelial cells)^54^ and *SFTPA1-201* (alveolar epithelial cells) ^55^, where GWAS signal did not achieve genome-wide significance. Among them, *MUC1-207*, encoding mucin-1, was significant in secretory transitional cells (**Fig. 6E**) and also colocalized with FEV_1_ in AT2 and alveolar transitional cells (**Fig. 6B**). In these cell types, lower *MUC1-207* levels were correlated with lower FEV_1_ and increased LUAD risk (**Supplementary Fig. 18A-B**), suggesting a potential functional connection. Protein structure prediction indicated that the 207 isoform lacked the extracellular mucin domain ^56,57^, a hallmark of all mucins, suggesting a unique function different from the canonical isoform (**Supplementary Fig. 18C**).

Collectively, isoQTLs identified previously undetected target genes from lung cancer and functions GWAS loci, especially in lung epithelial cell types, supporting common mechanisms among lung function and cancer.

### isoQTLs contribute to GWAS signals independently from eQTLs

Having identified previously unappreciated genes by bulk tissue and/or eQTL findings, we investigated whether isoQTLs contribute to GWAS signals independent from eQTLs. Among 49 colocalized eIsoforms for lung function or cancer GWAS signals, most (61%) were known isoforms, and 49% were relatively abundant isoforms (>14.4% abundance of that gene, **Methods**). Similar to colocalized isoforms, TWAS-significant isoforms were mainly of known (58.1%) and relatively abundant isoforms (47%) (**Supplementary Fig. 17D-E**), which was more pronounced among 12 isoforms from novel TWAS loci (91% known and 75% abundant isoforms). Given this observation, we evaluated whether the prevalence of known and abundant isoforms is due to isoQTLs mainly contributing to GWAS signals through total gene expression levels. For this, we asked whether GWAS-colocalized isoQTLs colocalize with eQTLs (**Fig. 5C**). Among 27 eIsoforms with matching eGenes in short-read-based eQTL in the same cell type, 48.1% did not colocalize with eQTLs. Collectively, 71% of the GWAS-colocalized eIsoforms were either not detected as an eGene in the same cell type or not colocalized with a matching eQTL, indicating an independent contribution of isoQTLs (**Fig. 7A**).

**Figure 7.**
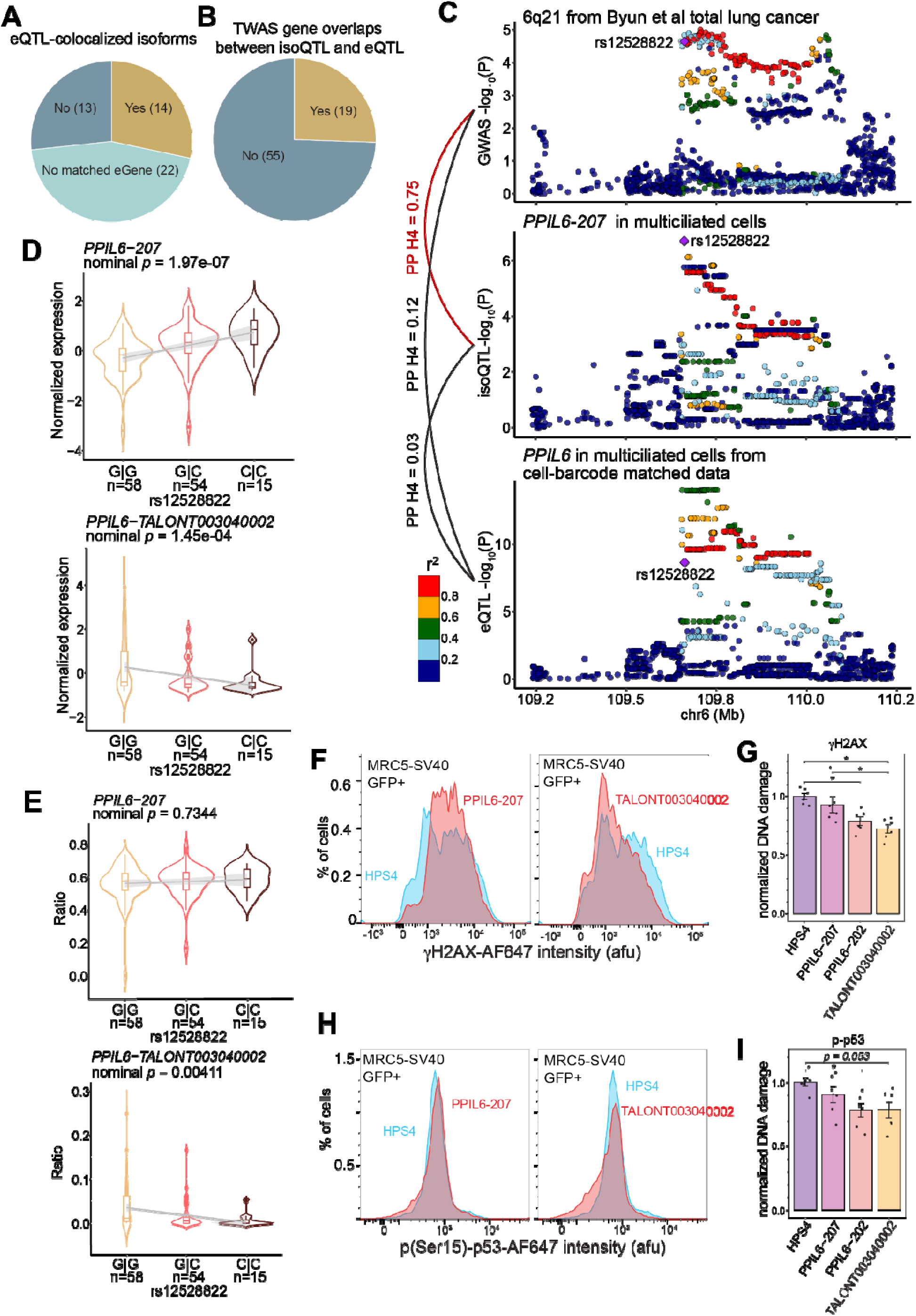
Distinct contribution of isoQTLs to GWAS signals. (**A**) Pie chart presents the fraction of GWAS-colocalized eIsoforms with an eQTL-colocalizing isoQTL for a matched eGene in the same cell type. (**B**) Pie chart presents the fraction of TWAS significant isoforms overlapping TWAS genes from cell-barcode matched short-read eQTL data (**C**) isoQTL or eQTL colocalization with lung cancer GWAS at the *PPIL6* locus in multiciliated cells. SNPs are color-coded based on the LD R^2^ (1000 Genomes, EAS, phase 3) with isoQTL lead SNP, rs12528822 (purple diamond). (**D**) Association between the genotype of lead isoQTL rs12528822 and the normalized expression of *PPIL6-207* (top panel) and *TALONT003040002* (bottom panel). (**E**) Association between the genotype of lead isoQTL rs12528822 and the proportion of *PPIL6-207* (top panel) and *TALONT003040002* (bottom panel) over all *PPIL6* isoforms. *P*-value is calculated by linear regression with the same covariates used in isoQTL mapping adjusted. Allele C of rs12528822 is the risk-associated allele for lung cancer. The violins and boxes are colored by genotype, and the grey line shows the trend of association. Center lines show the medians; the box indicates the middle of 50% of data; whiskers extend 1.5 times the interquartile range from the 25th and 75th percentiles; outliers are represented by dots; density of normalized expression is represented by the width of violin shape. (**F**, **H**) Representative histograms of DNA damage marker γH2AX (**F**) or p-p53 (**H**) for overproduced PPIL6 isoforms or HPS4 in GFP^+^ transfected cells from the same experimental batch. (**G,I**) Bar plots showed the DNA damage level (i.e., γH2AX or p-p53) normalized to the median intensity of GFP^+^ HPS4-overproducing MRC5-SV40 cells. Error bars show the mean±standard error. Black dots show the individual level of normalized value of two replicates from three experiments (n = 6). One-way ANOVA is used to test the differences across groups, * represents adjusted *p-*value < 0.05 by post hoc Tukey’s test. All summary statistics are in **Table S11**.

These eQTL-independent isoforms were mainly from previously not colocalized genes (83%). Similarly, we observed that 74.3% of TWAS isoforms were unique to the isoQTL dataset without a matched TWAS gene in any cell type in our cell-barcode-matched short-read data (**Fig. 7B**). These findings supported unique contributions of isoQTLs to GWAS signals that are largely independent from eQTLs.

Among eQTL-independent isoforms, *PPIL6-207* was a multiciliated cell-specific eIsoform and DEI (**Supplementary Fig. 12**) and colocalized with the overall lung cancer signal in a multi-ancestry GWAS (**Fig. 7C**). Intriguingly, *PPIL6* eQTL in the same cell type from the short-read dataset ^29^ did not colocalized with *PPIL6-207* isoQTL or lung cancer GWAS signals, although *PPIL6-207* was the most abundant isoform (**Supplementary Fig. 12**). Consistently, from the GTEx lung tissue, *PPIL6* eQTL did not colocalize with the lung cancer GWAS signal ^16^, and sQTL was not reported. These data indicated that cell-type- and isoform-specific genetic signals independent from total expression level contributed to this GWAS signal. To understand the biological mechanisms underlying this distinct isoQTL signal, we first investigated isoform dynamics of *PPIL6* in multiciliated cells. In addition to *PPIL6-207* (canonical isoform), *PPIL6-202,* and two novel isoforms (*TALONT003040002* and *TALONT003040049*) were also eIsoforms (**Supplementary Fig. 12A-B**). Among them, only *PPIL6-202* isoQTLs colocalized with short-read *PPIL6* eQTL, suggesting that the effect of this isoform is mainly through contributing to the total expression. Notably, *TALONT003040002* isoQTLs displayed the opposite allelic direction to the other isoforms (**Fig. 7D-E and Supplementary Fig. 19A**), suggesting a potential reciprocal regulation. We further performed isoform proportion QTL (**Methods**) for all qualifying *PPIL6* isoforms and observed that only *TALONT003040002* showed a significant QTL (*p* = 0.0041), suggesting that the proportion of this novel isoform could have a meaningful impact on the *PPIL6* isoform structure (**Fig. 7D-E, and Supplementary Fig. 19A**). To gain insights into the functional implications of this novel isoform, we performed protein structure prediction, as it was predicted not to be regulated by NMD. Compared to the two known isoforms (207 and 202), the novel isoform showed low predicted local distance difference test (pLDDT) scores for the cyclophiline-like domain of PPIL6, which implied an intrinsically disordered region or insufficient domain information (**Supplementary Fig. 19B-D**). A previous study reported that a knockdown of total *PPIL6* levels using siRNAs targeting a common region of multiple isoforms increased the endogenous DNA damage in immortalized lung fibroblasts ^16^. To further investigate the potential function of this novel isoform relevant to lung cancer risk, we performed cell-based DNA damage assays ^16,58^ of 202, 207, and TALONT003040002. Using the same cell line system, we overexpressed these three isoforms (**Supplementary Table 10**) and assessed their effect on the endogenous DNA damage as evidenced by γH2AX ^59^ and phospho-p53 (p-p53) ^60^ levels (**Methods**). Compared to the control gene, HPS4 *^58^*, overproduction of 202 and the novel isoform significantly decreased γH2AX (adjusted *p* = 0.018 and 0.0018, respectively) (**Fig. 7F-G and Supplementary Fig. 20A-B**). The 202 isoform also displayed significantly decreased p-p53 levels (adjusted *p* = 0.049), whereas the novel isoform showed a non-significant trend (adjusted *p* = 0.053) (**Fig. 7H-I and Supplementary Fig. 20C**). Notably, the 207 isoform did not show any effect on endogenous DNA damage in both assays and further showed a significant difference from the novel isoform in γH2AX levels (adjusted *p* = 0.022). The observed cellular function of the novel isoform suggested that the novel isoform might contribute to the GWAS-colocalizing isoQTL signal of the canonical 207 isoform through reciprocal allelic proportion change. Consistent with this notion, the lung cancer protective allele is associated with lower expression of the 207 but higher expression of the novel isoform, which aligns with lower endogenous DNA damage in the cell line data upon overexpression (**Supplementary Figs.19-20**).

Collectively, our data demonstrated the independent genetic effect of cell-type-specific isoQTLs contributing to lung-related GWAS signals that were unexplained by eQTLs. To share our full-length lung cell atlas and cell-type-level isoQTL data, we established an online tool, “ISOLUTION: ISoform QTL of LUng cell Types by lONg-read sequence” (https://appshare.cancer.gov/ISOLUTION). ISOLUTION provides: 1) full transcript structures of post-QC isoforms; 2) isoform expression patterns across 37 lung cell types and 129 individuals; 3) searchable summary statistics of eIsoforms and isoQTLs, and 4) eQTL summary statistics and expression data based on the barcode-matched short-read dataset.

## Discussion

In this study, we present an isoform-level lung cell atlas and deploy an open-access web interface, ISOLUTION, for accessing the data across cell types and individuals. From this atlas, we identified 270,536 novel transcript isoforms and characterized cell-type signatures of isoform expression in lung tissues from never-smokers. Supported by the multi-layered validation including proteome profiling, cell-barcode matched short-read data, and single-cell multi-omics data, our dataset provides a reference for annotating transcript isoforms and exploring isoform functions in lung cell types. The sample size of our dataset enabled the cell-type-specific isoQTL detection, and systematic comparisons with cell barcode-matched short-read eQTL dataset uncovered the independent genetic effects of isoQTLs beyond the total gene expression level. Integration of isoQTLs with lung cancer subtypes and lung function GWAS identified new susceptibility isoforms and highlighted contributions of isoQTLs to the GWAS signals distinct from eQTLs.

Our study design demonstrated the feasibility of harnessing single-cell long-read RNA-seq at scale to detect isoform diversity. Previous bulk tissue transcriptomes using long-read sequencing reported surprisingly large proportions of novel isoforms that are tissue-specific albeit in limited sample sizes ^6^. A recent study of GTEx tissues observed a strong effect of ancestry on the inter-individual variation of alternative splicing and the effect of sex on expression variation ^61^. Considering added technical variability by sequencing batches, our design including 129 individuals of similar genetic background, same sex, and without smoking effect, in 22 cell-type balanced batches helped reduce the dataset-wide variability for reliable isoform detection. Notably, doubling the sequencing depth enabled the detection of initially single-batch-specific isoforms in additional batches. These data, together with the peptide-level validation of novel isoforms, suggested that a substantial proportion of batch-specific novel isoforms could be further validated at a higher sequencing coverage.

The isoform-level lung cell atlas allowed a systematic investigation of the lung cell type signatures of isoform expression. Single-cell full-length transcriptomes exploring isoform specificity have been limited to brain tissues^7,9,10^ and tumor tissues^62^ with tens of samples from human and mice. Beyond anecdotal examples ^9^, our systematic analysis observed isoform-level signatures showing larger cell-type differences than gene level across 37 lung cell types, including more distinct marker genes at the isoform level. Moreover, we investigated the contribution of isoform information to cell clustering and cell type annotation. Although gene-level clustering outperformed isoform-level in dataset-wide comparisons, isoforms better distinguished a subset of individual lung cell types, consistent with the previous findings that isoforms could separate excitatory neuron subtypes ^7^. Given that current cell type annotation and iterative clustering rely on established marker genes and conformity to existing reference datasets, ground truth was set based on gene-based annotation, potentially favoring gene-level clustering and annotation. Further investigations will be needed on how to integrate gene-level and isoform-level data to maximize the benefit of both features in distinguishing cell types and cell subtypes in single-cell analyses.

Isoform-level QTL analysis uncovered the genetic effects on isoform regulation that are distinct from total gene expression levels, including those contributing to lung traits and diseases. Taking advantage of our cell-barcode matched short- and long-read datasets, we determined that only a subset of isoQTLs contributed to the regulation of total gene-level expression (**Fig. 5C**). Intriguingly, isoQTLs were enriched for the functions relevant to splicing and other post-transcriptional regulation, whereas eQTLs were not. These findings are consistent with the observations in GTEx tissues, where no overlap was observed between junction-based sQTL and eQTL credible sets in 39.4%–54.9% matched genes across tissue types. Our data extended this finding to cell-type-resolved full-length isoforms by demonstrating that eQTLs and isoQTLs are relatively matched for detected cell types, but a substantial proportion are not colocalized, with more distinct signals coming from novel isoforms. Colocalization and TWAS of lung cancer and lung functions GWAS persistently demonstrated that isoQTLs uniquely contribute to GWAS signals. The majority of GWAS-colocalized isoQTLs were independent from eQTLs, and most of TWAS findings did not show matched genes in eQTL data. Consistent with our findings, a recent isoTWAS study using bulk-tissue short-read data found only 50 overlapping genes between 353 isoTWAS and 306 TWAS significant genes in the context of lung cancer ^28^. These data suggested that isoQTLs can contribute to some of the GWAS heritability previously unexplained by eQTL and other mechanisms.

The example of *PPIL6-207* for lung cancer provides one of the potential mechanisms underlying independent contribution of isoQTLs. Although *PPIL6-207* displayed a unique GWAS colocalization that *PPIL6* eQTL in the same cell type did not, it was paradoxical as *PPIL6-207* is the canonical and most abundant isoform. A potential explanation could be through a reciprocal isoform regulation with a novel isoform, *TALONT003040002,* via alternative splicing. Although this relatively low-expressing novel isoform did not colocalize with the GWAS signal, the proportion of this isoform among all the *PPIL6* isoforms was uniquely associated with the *PPIL6-207* isoQTL lead SNP. This data suggested that allelic alternative splicing effect captured by the main isoform could be partially tagging the effect of the novel isoform. The observed cellular function of this novel isoform in reducing the endogenous DNA damage and apparent null function of the main isoform, 207, conform with this notion. Moreover, the allelic direction of lung cancer risk allele being associated with lower levels of the novel isoform is consistent with heightened DNA damage leading to tumorigenesis. Given that another isoform, *PPIL6-202*, also reduced DNA damage in lung cells but mainly contributed to total expression and eQTL signal, it is likely that the novel isoform is not the sole player in the context of endogenous DNA damage. The precise mechanism of *PPIL6* isoform dynamics mediated by isoQTLs and how their function in multiciliated cells could contribute to lung tumorigenesis need to be explored further.

GWAS integration highlighted common genetic regulation underlying lung cancer and lung functions in alveolar epithelial cells and multiciliated cells. Previous genetic correlation and causal effect studies as well as overlap of GWAS loci and target genes implicated shared genetic contribution to lung function traits and lung cancer, especially adenocarcinoma. In addition to consolidating these findings, our data provided isoform and cell-type contexts for known genes and identified new genes that are common in both phenotypes. For example, lower expression of the *SECISPB2L* isoforms was correlated with impaired lung function and increased lung cancer risk in AT2 cells, facultative progenitors for alveolar regeneration and LUAD origin. *SECISPB2L* encodes SECIS binding protein 2-like that has been reported to display an inhibitory activity on lung cancer cell proliferation ^63^. It is possible that an increased cell proliferation might lead to a compromised homeostasis in lung repair process and favors tumorigenesis.

*MUC1* is elevated in epithelial cancers, including lung cancer, especially adenocarcinoma, and is a well-known tumor antigen^57^. Although it was identified in lung function GWAS, the locus has not reached genome-wide significance in lung cancer GWAS. In our data, lower levels of *MUC1-207* were correlated with increased risk and impaired lung function in AT2 and transitional cells. This isoform encodes a mucin 1 protein lacking all the mucin repeats forming the extracellular domain that can function as both barriers and signaling molecules ^56,57^. For example, lack of the domain interaction with NFκB p65 can impact pathogen clearance via interference of pro-inflammatory cytokine expression^56,64^. Other domains such as cytosolic tail might also have roles in tumorigenesis via phosphorylation or beta-catenin binding^57^. In addition to alveolar epithelial cells, we also observed that isoforms in multiciliated cells were frequently colocalized with both lung function and cancer GWAS, albeit with different genes. The roles of multiciliated cells in lung function and associated conditions have been known in the context of cilia dysfunction or loss followed by mucociliary clearance dysfunction leading to COPD ^47,65^. However, multiciliated cells were less studied in lung cancer development compared to the known cell types of lung cancer origin. Our data revealed potential connections of respiratory dysfunction involving multiciliated cells that could contribute to the cellular environment promoting lung tumorigenesis.

In conclusion, we provided a single-cell long-read sequencing dataset of lung tissues with a cell-barcode-matched short-read dataset, which has a broad implication for investigating isoforms and alternative splicing and facilitating the understanding of genetic effects of GWAS signals in the context of cell types and lung diseases.

### Limitations of the study

We recognize that our study has several limitations. First, even though we used twice the standard read-depth (2 SMRT cells) for each single-cell batch, the isoform-level expression profiles at the single-cell level are still sparse. Increasing sequencing depth up to 4 SMRT cells could reduce the sparsity to a degree, but it also proportionally increases the number of novel isoforms with lower abundance that need to be further validated. Reassuringly, we found that the cell type specificity of isoform expression did not result from the sparsity, by increasing the sequencing depth. Second, although our scale enabled us to perform QTL mapping, the isoQTL findings were limited by the detection power due to a modest sample size. Allele-specific isoform expression and alternative splicing would be worthwhile to explore more genetic effects in future studies. In terms of isoform identification, we performed in-sample proteomics validation with 11% validated at the peptide level, but the functions of both known and novel highlighted isoforms and their roles in lung biology and diseases need further experimental validation.

## Methods

### Human subjects and tissue collection

Sample collection for this study and the companion short-read eQTL study ^29^ was based on the same process as our published study ^31^. We surgically collected tumor-distant normal lung parenchyma tissues from 129 patients mainly diagnosed with primary lung adenocarcinoma. Tumor-distant normal samples were obtained from the periphery of the resected lobe, more than 2 cm away from the tumor edge, which is unlikely to present tumor phenotype based on guidelines of the National Comprehensive Cancer Network. All participants were treatment-naive at the time of surgery and did not have other active lung diseases, such as interstitial lung disease or infectious lung disease. Among the 129 patients, 102, 12, and 7 were diagnosed with stage I, II, and III lung adenocarcinoma, respectively. Two patients were diagnosed with adenocarcinoma in situ (pTisN0) and another two patients were found to have pleural seeding metastasis during surgery and were diagnosed with stage IV disease. Four patients underwent surgical resection for diagnostic and therapeutic purposes due to suspected lung cancer but were confirmed to have benign lesions (nodular dense fibrosis, multifocal interstitial fibrosis, chronic granulomatous inflammation, and lymphoplasma cell proliferation, respectively). All 129 participants were self-reported female, ethnically Korean, and never-smokers (individuals who have smoked < 100 cigarettes in their lifetime). The sample size was determined based on a single-cell eQTL power calculation study^66^. Detailed characteristics of the patients included in the study are in **Supplementary Table 1**. Patient tissues were obtained under a protocol approved by Yonsei University Health System, Severance Hospital, Institutional Review Board (IRB 4-2019-0447, 4-2022-0706), and informed consent was obtained from each patient prior to surgery. All samples were collected primarily to study risk factors and prognoses of respiratory diseases. The donors consented to the secondary use of their samples for comprehensive research purposes, including providing data containing personally identifiable information such as sex, age, and the medical center. All experiments were performed following applicable regulations and guidelines.

### Genotyping, QC and imputation

Genomic DNA was extracted from the buffy coat using the QIAamp DNA blood kit (Qiagen, cat. 51185). Genotype data were generated using the Illumina Infinium Global Screening Array (v2) according to the manufacturer’s protocol. Genotype data were exported in PLINK format (v 1.9.0) and filtered to include SNPs with sample call rate > 95%, genotype cell rate > 95%, Hardy-Weinberg equilibrium *p*-value > 10^-6^, and minor allele frequency (MAF) > 1%. A/T & G/C SNPs with MAF > 0.4, or SNPs with more than 20% allele frequency difference compared to 1000 Genome phase 3 or not in 1000 Genome phase 3 reference were further excluded. The genotype data with 354,170 SNPs after QC were separated by chromosome and submitted to the Michigan Imputation Server using 1000 Genome (phase 3 v5) EAS population as reference for imputation. Minimac4 (v1.0.2) and Eagle (v2.4) were run on Michigan Imputation Server ^67^. The imputed VCF data was lifted over to hg38 using bcftool (v1.19).

### Single-cell processing, cell-type balancing, and multi-plexing

Fresh tissue dissociation and cryopreservation conditions were optimized to maximize epithelial cell retention as described before^31^. Briefly, tissues were resected at a ∼2 cm^3^ size and submerged in MACS Tissue Storage Solution (Miltenyi Biotec cat.130-100-008) within 30 mins to be stored at 4°C for the same day dissociation. Tissues were chopped into 2mm diameter pieces and dissociated using Multi Tissue Dissociation Kit 1 (Miltenyi Biotec, cat. 130-110-201) with a reduced amount (25%) of enzyme R on gentleMACS Octo Dissociator (Miltenyi Biotec, cat. 130-096-427). Red blood cells were subsequently removed using Red Blood Cell Lysis Solution (Miltenyi Biotec cat.130-094-183). Single-cell suspension of each sample was frozen using 90% FBS and 10% DMSO and stored in liquid nitrogen until further processing except for the time of transfer in dry ice.

Prior to single-cell capture, cell-type proportions balancing and sample multi-plexing were performed as described before ^31^. Briefly, we performed FACS sorting of four major lung cell type categories and balanced their ratio to enrich epithelial cells and reduce immune cell proportions before multi-plexing samples from ∼6 individuals per single-cell batch. Thawed single-cell suspensions were filtered using 70 μm pre-Separation Filters (Miltenyi Biotec, cat. 130-095-823) and labelled using antibodies against EPCAM (ThermoFisher Scientific, cat. 25-9326-42), CD31 (BioLegend, cat. 303116), and CD45 (BioLegend, cat. 304006), and stained with DAPI (ThermoFisher Scientific, cat. D21490). DAPI^-^ live single-cells were sorted on BD FACSAria Fusion Flow Cytometer (BD Biosciences) using three gates: EPCAM^+^CD45^-^ (designated “epithelial”), EPCAM^-^CD45^+^ (designated “immune”), and EPCAM^-^CD45^-^ (designated “endothelial or stromal”) (gating strategy described in Long et al ^31^). For balancing, we used ∼6:3:1 ratio of “epithelial”: “immune”: “endothelial or stromal” cells for each individual sample. An equal number of remixed cells from ∼6 individuals were pooled for loading to the 10X Genomics Chromium Controller (100069, 10X Genomics, USA).

### Long-read MAS-seq single cell library construction and sequencing

cDNA libraries were prepared using the Chromium Next GEM Single Cell 3’ Reagent Kits v3.1 following the manufacturer’s guidelines (CG000315-Rev D, 10X Genomics, USA). From these cDNAs, MAS-seq (Multiplexed Arrays Sequencing) single cell libraries were constructed using the MAS-Seq for 10x Single Cell 3’ Kit (102-659-600, PacBio, USA) following the manufacturer’s procedure (102-678-600, PacBio, USA). The same set of cDNAs was used for short-read sequencing libraries in the companion study ^29^. MAS-Seq single cell libraries were sequenced on PacBio Revio system using PacBio v2.0 chemistry. MAS-seq application and MAS SMRTbell adapters and barcodes were chosen from the SMRT link run design for sequencing. Each of the 22 batches was sequenced on 2 Revio SMRT cells and achieved 9 -10 million HiFi reads. Segmentation of HiFi reads yields 138 to 162 million reads with a mean read length of 771 bp. For sequencing depth benchmarking, two of the 22 batches (SCBatch4 and SCBatch19) were sequenced on a total of 4 Revio SMRT cells and achieved 17-22 million HiFi reads and 270-335 million segmented reads. The segmented reads were used for downstream analysis.

### Processing of PacBio full-length reads

The full-length reads processing was guided by the recommended MAS-SC and ISO-seq ^68^ pipeline by PacBio (https://isoseq.how/umi/). The raw sequencing data from each run were processed by *ccs* v4.0 to generate circular consensus sequencing (CCS) reads requiring at least three full pass subreads and predicted accuracy ≥ Q20. Primers were removed and barcodes were identified by *lima*, only keeping the full-length non-chimeric (FLNC) reads containing both a 5’ and a 3’ primer followed by a poly-A tail. The UMIs and cell barcodes were clipped from the reads using the T-12U-16B design using *isoseq tag*. PolyA tail and artificial concatemers were removed by *isoseq refine*. The full-length reads from the two SMRT cells of the same single-cell batch were combined for the following steps of the analysis. To address the PCR duplication, we used *isoseq groupdedup* to generate one CCS read per founder molecule. The deduplicated reads were aligned and mapped to the reference genome (GENCODE v32) downloaded from 10X genomics (https://www.10xgenomics.com/support/software/space-ranger/downloads). The mapped reads were collapsed into unique isoforms which were classified into structural categories by *Pigeon classify,* based on SQANTI3. Simultaneously, the abundance of each isoform was counted. The isoforms were filtered out according to SQANTI3 quality control indicators, including low coverage (< 3 reads per batch), the presence of noncanonical splice junctions, the existence of intrapriming (0.6 as cutoff), and potential reverse transcriptase switching events. As described in the SQANTI paper ^69^, GT-AG, as well as GC-AG and AT-AC, were considered as canonical splice junctions, which altogether account for 99.9% of all human introns.

### Cell calling, demultiplexing, doublet detection and quality control

For single cell detection, we took advantage of our barcode-matched short-read dataset that uses the same cDNA samples to provide guidelines for cell number expectation, as this function is not available in ISO-seq pipelines. We performed a percentile method using a range of percentiles (95-99 with 1% increment) to approximate cells. The percentile method first identifies the specific percentile of UMI counts per cell and generates the UMI cutoff by a factor of 10. The cell numbers of each single-cell batch using different percentiles were compared with our short-read results, and the percentile approximating the smallest number that exceeds the short-read result was determined and selected for each batch. The *--method percentile* argument with optimized percentiles (**Supplementary Table 12**) was used for the command of *isoseq bcstats* for cell calling. Vireo ^70^ was used to demultiplex and assign cells to each individual based on genotypes. We used *bcftools mpileup* with -X pacbio-ccs argument to call variants from the aligned and de-duplicated reads from the ISO-seq pipeline ^71^. The bi-allelic variants with QUAL > 51 were retained as reference for single-cell genotyping, which was estimated by cellSNP-lite with *--minMAF 0.1 --minCount 20 ^72^*. The output single-cell genotypes from cellSNP-lite were used in vireoSNP with germline genotypes from the blood DNA of pooled individuals for each single-cell batch. A cell was assigned to the individual of the best singlet from vireoSNP ^70^ if the maximum probability for a single individual was larger than the probability of doublet. Otherwise, the cell was labelled as a doublet. Besides genotype-based doublets, we performed scrublet ^73^ to determine doublets based on gene expression. Cell by gene matrix and cell by isoform matrix using genes or isoforms expressed by more than 5 cells for each single-cell batch were loaded to Seurat. Cells with the number of detected RNA < 200 or percentage of mitochondrial RNA > 25% were dropped as low-quality or dead cells. Cells labelled as doublet by either vireoSNP or scrublet were removed as potential doublets.

### Isoform detection and characterization

We integrated the transcriptomes from 22 single-cell batches using TALON ^74^ after initial cell filtering described above. The processed full-length reads from all batches were merged to generate the combined genome reference annotation file. To further curate the transcriptomes from single-cell long-read sequencing, we used TransDecoder to filter out novel isoforms without open reading frames (ORFs) using default parameters. We utilized the Pfam and Blastp databases to search for homology to known proteins to screen ORFs with protein-coding potential. We only retained the transcripts with ORF marked as complete length for the novel isoforms classified by *Pigeon*. After these QCs, the number of isoforms was decreased from 831,469 to 325,864. As TALON does not recognize the fusion gene category, we kept the category labels for isoforms classified as fusion genes by SQANTI3.

To validate the robustness and reliability of our isoform detection, we compared the integrated and curated transcriptome with GENCODE v32 and GTEx long-read sequencing dataset of bulk tissues by using *gffcompare*. While comparing with GTEx long-read dataset, the isoforms marked as “=” in *gffcompare* meaning exactly matching at the “intron-chain”-level (**Supplementary Fig. 1A**) were considered as the replicated isoforms.

To cross-validate long-read based splice junctions with those from the barcode-matched short-read data, leafcutter ^75^ was used to first detect short-read-based splice junctions. The BAM files of the short-read data mapped to GENCODE v32 by CellRanger v7.0.1 were split into the individual level based on cell barcodes using *sinto* v0.10. According to the documentation of leafcutter, the *regtools* function was used to extract junctions from BAM files and identify intron clusters as splice junctions. The leafcutter-identified splice junctions were compared with splice junctions detected in 325,864 post-QC full-length isoforms based on the coordinates for cross validation.

For the analysis of non-sense mediated decay (NMD) sensitivity, we leveraged the coding sequence (CDS) results from TransDecoder and defined isoforms carrying a premature termination codon located 50nt or more upstream of the last splicing junction as NMD sensitive. To characterize the potential post-transcriptional functions of eIsoforms, we employed SUPPA2 v.2.4 ^76^ to define alternative splicing events, including alternative 3′ splicing (A3), 5′ splicing (A5), first exon (AF), last exon (AL), retained intron (RI), skipped exon (SE), and mutually exclusive exons (MX).

To predict the potential protein structure of isoforms, AlphaFold 3 model ^77^ was used via AlphaFold server (https://alphafoldserver.com/). The output peptide sequence from TransDecoder was used as input for prediction. The structures are colored by predicted local distance difference test (pLDDT) or predicted domain by AlphaFold 3. The domain is annotated based on Protein Data Bank (PDB) by The Encyclopedia of Domains (TED)^78^.

### Peptide-level validation of abundant isoforms

#### Digestion and Sample Processing

To validate novel isoforms identified by long-read sequencing, we randomly selected 2 individuals each from 2 randomly selected single-cell batches (NCI134, NCI135, NCI138, NCI139; 4 samples in total). Samples were resuspended in 200 uL EasyPepTM (Thermo Scientific) Buffer with 1 uL nuclease and Phos Stop. 5 ug of Trypsin/LysC with 50 uL Reduction Solution and 50 uL Alkylation solution was added to each sample and allowed to digest overnight, shaking at 500 rpm at 37°C. The next day, reactions were quenched with 50 uL of Digestion Stop Solution. EasyPep Mini plate was used for sample clean up, and samples were dried in the speed vacuum.

#### Fractionation

Off-line fraction was accomplished using high pH reversed phase separation on a Waters Acquity UPLC. Peptides were loaded onto an XBridge Peptide BEH C18 column and eluted with a 6-20% gradient of 90% acetonitrile with 10 mM Ammonium Formate pH 9.4 over 1.5 minutes, 20-70% over 68.5 minutes, and 70-90% over 5 minutes with a flow rate of 0.35 mL/min. Fractions were collected for 1 minute and were combined into 24 final fractions before being dried prior to analysis.

#### LC/MS analysis

The dried peptides were reconstituted in 0.1% formic acid and loaded onto DNV PepMap Neo C18 LC column (Thermo Scientific, CA) utilizing a Thermo Easy nLC 1200 LC system (Thermo Scientific, CA) connected to an Orbitrap Fusion Lumos. Peptides were eluted with a 5-35% gradient of 80% acetonitrile with 0.1% formic acid over 90 min, a 35-50% gradient over 30 minutes, and a 50-95% gradient over 3 min with a flow rate of 300 nL/min. The instrument was run in the TopSpeed method with the MS1 performed at 120,000 resolution over a mass range of 4000 to 1600 m/z, with an auto maximum injection time and standard AGC target. MS2 scans were acquired in the Orbitrap with a resolution of 15,000, normalized collision energy set at 30, with an auto maximum injection time and standard AGC target, isolation window of 1.6 m/z and charges of 2-5 for MS2 selection.

#### Database Search and Post Process Analysis

The MS files were searched in Proteome discoverer using the Sequest node. Due to mass-spec inherently favoring abundant proteins, we searched the mass-spec data against the isoforms expressed in > 20% individuals in each cell type. This criteria was the same as those for DEI detection and isoQTL mapping analyses where isoforms were required to be relatively abundant and detected in more than 2 single cell batches, thus avoiding isoforms biased in a single batch. The peptide fasta of these qualifying isoforms was generated by TransDecoder. The search was performed using full trypsin digestion with a max 2 missed cleavages, Min peptide length 6, MS1 mass tolerance of 20ppm, MS2 tolerance of 0.02 Da, variable modification of oxidation on methionine and static modifications of carbamidomethyl on cysteine. Percolator was used for FDR analysis. The isoforms with >=2 unique peptides in any sample or detected in >=2 samples with high confidence (FDR < 0.01) were considered as validated isoforms

### Clustering and cell type annotation

After removing doublets detected by vireo and scrublet, we performed automatic cell type annotations using Azimuth and scArches as initial guidelines, followed by iterative clustering for the final cell type annotation. Gene-level expression profiles were integrated and projected to the HLCA core dataset by scANVI using pre-selected feature genes (1,841 out of 2,000). The *FindCluster* function with the Leiden algorithm and a resolution of 2.0 was used for clustering after integration, which resulted in 43 clusters. Cell clusters were classified into 4 categories of lung cell types based on canonical cell category gene expression (*i.e., EPCAM* for epithelial*, PTPRC* for immune*, CLDN5* for endothelial and *COL1A1* for stromal). We dropped clusters with > 10% of cells expressing more than one canonical category marker or > 10% of cells automatically annotated as a different cell category. To further annotate cells within the cell category, highly variable genes were re-selected within each category for re-clustering by SCTranform combined with Harmony. The Leiden algorithm was used with a resolution of 0.5 except for the immune category, where 1.0 was used. Highly expressed genes for each cluster were calculated using *FindMarkers* function and compared with HLCA marker genes ^35^. Clusters showing signatures of more than one cell type within the category were dropped as doublets. The remaining clusters were annotated according to HLCA cell type markers while cross-referencing automatic annotation results.

### Differential isoform expression analysis

For differential expression analysis, we pseudo-bulked raw counts by aggregating single-cell level matrices within cell types for individuals with more than 5 cells for that cell type. Isoforms expressed by more than 20% of individuals in each cell type were kept and tested in differential analysis. For example, in the rarest cell type, neuroendocrine, 31 individuals met the criteria of pseudo-bulking, and isoforms expressed by more than 20% of individuals should be at least detected in 7 individuals, *i.e.*, at least two single-cell batches, thus avoiding isoforms detected in only one single-cell batch from being tested. Based on these criteria, a total of 42,824 isoforms were kept for further analysis, with the minimum read counts being 13 for the lowest abundant isoform. This cutoff showed a high correlation of individual-level isoform expression (median r = 0.93) across sequencing depths in each cell type, indicating robust expression across individuals (**Supplementary Fig. 5B**). Differential expression analysis was performed with DESeq2 using a negative binomial model to compare one cell type versus all the other cell types. Significance of differentially expressed isoforms (DEIs) was claimed as adjusted *p*-value < 0.05 and log_2_ fold change > 1. Differentially expressed genes (DEGs) were identified following the same pipeline.

For FDR control, we generated the permuted pseudo-bulk expression data and performed differential analysis for each cell type. Using different cut-offs for adjusted *p-*values to claim significance (not accounting for fold change), we found that false positives were well controlled (**Supplementary Fig. 21A**) in most cell types except for three rare cell types (*i.e*., mast cells, myofibroblasts and SMC). To further check the inflation of FDR in these cell types, we performed the permutation 1,000 times and evaluated how many times one isoform is falsely claimed as a significant isoform. We considered an isoform as a potential false positive, if it was falsely claimed as significant for more than 50 times out of 1,000 permutations when using FDR < 0.05. We found that 10.5%-21.9% of isoforms were potential false positives in these rare cell types, which is higher than 5.2% in AT2, a control cell type (**Supplementary Fig. 21B**), suggesting a higher potential of detecting false positives in these three rare cell types. To further characterize the potential false positives in the three cell types, we checked the distribution of adjusted *p-*values and log2 fold changes in volcano plots (**Supplementary Fig. 21C**). We found that most (>90%) potential false positives are from the down-regulated isoforms, which might be caused by the smaller sample sizes, where many isoforms could be at a lower level compared to the rest of the dataset. Since we set the criteria for claiming DEIs at FDR < 0.05 with log_2_ fold change >1 (up-regulated genes only), we believe most of the significant findings were not affected by the potential false positive findings.

To benchmark our pseudo-bulk differential analysis method with a single-cell level method, we followed the pipeline of Patowary *et al* ^7^ to detect single-cell level DEIs. This single-cell level method identified more DEIs (24,773) than the individual-level DEIs (19,595). However, this gain seemed to be mainly from low-expressing isoforms (73% being expressed by < 1% of cells vs. 59% of individual-level DEIs) and single-batch expressed isoforms (10% vs. >=2 batch requirement for individual-level DEI), while not increasing the overall diversity of DEI detection (average 1.4 DEIs per gene across cell types vs. 1.8 per gene for individual-level DEI). Collectively, individual-level summary and stringent filtering resulted in more robust and diverse DEI detection across cell types while avoiding preferential detection of low-expressing and single-batch expressed isoforms.

### Mapping isoQTLs in each cell type

For isoQTL mapping, we employed jaxQTL^37^ for the negative binomial regression model and TensorQTL ^79^ for the simple linear model using variants within a ±1MB *cis*-window of TSS of an isoform. isoQTLs were only tested in the cell types contributed by more than 40 individuals with more than 5 cells for that cell type. Isoforms expressed by more than 20% of individuals of each cell type were included in isoQTL mapping, to ensure that the tested isoforms were not entirely from a single sequencing batch. We created the raw pseudo-bulked counts of isoforms (by sum) for the NB model and adjusted for per-individual library size as offset, whereas standardized log-transformed normalized expression levels were used in TensorQTL. The isoform levels were used as the phenotype in isoQTL analysis. For genotype data, variants with MAF < 0.05 or imputation R^2^ < 0.3 were filtered out, and the top three genotype principal components (PCs) calculated by plink (v 1.9.0) were used as covariates to account for the residual population structure. Expression PCs were calculated to account for the non-genetic hidden variables in the data. The numbers of PCs for each cell type were determined by an automatic elbow detection method in the R package *PCAforQTL ^80^.* The final covariates used for regression models are 3 genotype PCs, a customized number of expression PCs, single-cell sequencing batch (accounting for multiplexing individuals), and age. For TensorQTL analysis, PEER ^81^ was used instead of expression PCs following the same covariate selection process for each cell type. Following previous studies ^82^, the permutation-based Beta approximation method was used to calibrate isoform-level *p-*value for the correction of multiple tests of variants using 1,000 permutations in both jaxQTL and TensorQTL. Storey and Tibshirani procedure was applied for genome-wide FDR correction, and the significance threshold was set as FDR < 0.05 to claim significant isoforms (eIsoforms). To nominate all significant isoQTLs associated with eIsoforms, a nominal *p-*value threshold for each isoform was calculated based on the Beta distribution model of each isoform and the closest beta approximated *p*-value to 0.05 FDR within cell types ^82^.

To assess the type I error control of negative binomial models for identifying eIsoforms (**Supplementary Fig. 9**), we generated permuted data for each cell type by randomly shuffling expressed counts within each tested isoform with different seeds. The proportion of true null hypotheses (π_0_) was calculated by the *qvalue* function to evaluate the type I error control. By evaluating the isoform-level *p-*values calculated by permuted-based method from permuted data, we found there were few isoforms (0-0.16% of total tested isoforms across 33 cell types) with strong significant *p-*values (beta approximated *p-*value < 10^-5^) in permuted data, which showed no enrichment in highly-sparse isoforms or variants with highly imbalanced allele frequencies representatively. Due to our modest sample size, tens of thousands of tested isoforms, and the sparse expression of isoforms, these potentially inflated isoforms could be detected by chance. Considering the tiny fraction of potential inflation and well-calibration in simulation data^37^, we think the negative binomial model is overall well-calibrated for type I error.

To explore the cell type specificity and sharing of isoQTLs, we performed mash ^38^ to harmonize the effect sizes of isoQTLs across cell types. The lead isoQTLs of eIsoforms from each cell type were defined as the strong set. Considering the sparsity of the isoQTL results, the random set was determined by randomly selecting 1,000,000 isoQTLs from all tested variants in the *cis*-window. The EZ model of mash was applied using Z scores instead of effect sizes, where the effect sizes of isoform and variant pairs that were not tested in the cell types were imputed as 0. The strong set was used to learn data-driven covariance matrices and the random set was used to learn correlation among isoQTLs across cell types. After fitting the model, the posterior means of Z scores and local false sign rate (LFSR) were calculated by applying the fitted model to the strong set. The harmonized Z scores were converted to effect sizes by dividing the standard error from initial isoQTL results for cross-cell-type comparison.

To evaluate whether our isoQTLs detected genetic signals specific to Asian populations, the allele frequencies of lead isoQTLs were summarized and compared using the EAS and EUR populations from the 1000 Genomes Project (**Supplementary Table 4**). In addition, we evaluated whether independent QTL signals were identified for genes having both isoQTLs and sQTLs (GTEx v8 lung tissue). Since an isoGenes can possibly have multiple lead isoQTLs for different eIsoforms, we first determined “independent” lead isoQTLs for each gene by keeping one variant out of one LD block if there were multiple lead isoQTLs in high LD (R^2^ > 0.8). For the 847 “independent” lead isoQTLs not previously identified as any significant sQTL for the same genes in GTEx v8, we determined the isoQTLs with R^2^ < 0.1 with sQTL or mono-alleleic in the EUR population as EAS-specific QTLs.

To investigate the association between *PPIL6-207* isoQTL (rs12528822) and the composition of isoforms within *PPIL6* gene, the ratio of the isoform was calculated by dividing the summed count of PPIL6 isoforms for each individual after pseudo-bulking. The association between genotype and isoform ratios was tested by linear regression with the adjustment of the same covariates used in isoQTL mapping. With 4 eIsoforms tested, significance was claimed by *p*-value < 0.0125 (0.05/4).

### Functional annotation of isoQTLs

Since alternative splicing is regulated through the binding of spliceosome proteins and RNA-binding proteins to splice junction sites or splicing enhancers, we evaluated if the isoQTLs were: 1) directly located in splice junction sites or 2) located in the binding motifs of RNA-binding proteins. We defined the splice junction sites as the first and last 2 base pairs of introns, which are the main recognition and binding sites during the splicing process ^83^. The introns were identified and extracted from our curated single-cell long-read genome reference using the function of *talon_get_sjs*, including both novel and known introns. Significant isoQTLs were overlaid with the 4 bps of introns by matching the positions to identify sjs-isoQTLs. To assess if sjs-isoQTLs are associated with more than one isoform, we used Fisher’s exact test to compare whether the proportion of sjs-isoQTLs associated with more than one isoform is larger than that of isoQTLs not located in splice junction sites. To reduce the redundancy generated by LD, we pruned the isoQTLs using plink *--indep-pairwise 200 1 0.1*.

For the binding motifs of RBPs, we downloaded the dataset of enhanced crosslinking and immunoprecipitation (IP) followed by sequencing (eCLIP-seq) on the K562 cell line including 120 RBPs from the ENCODE project phase III ^40^. Peaks by eCLIP-seq passing the stringent cutoffs (fold enrichment ≥8 and P ≤ 0.001) as described in the ENCODE project were retained for the following enrichment analysis. Peaks were annotated into functional categories in accordance with the functions of corresponding RBPs. Genomic coordinates of motif peaks were lifted over from hg19 to hg38 for matching the position of isoQTLs. Similar to the splice site analysis, we focused on LD-pruned variants and performed Fisher’s exact test to evaluate if isoQTLs were enriched in the functionally annotated RBP binding motifs compared with non-significant variants. Addressing multiple tests for functional annotations, annotations with FDR < 0.05 were claimed as significant.

To determine the potential alternative promoters, we overlaid the single-cell ATAC-seq peaks with the TSSs of eIsoforms exhibiting alternative first exon (AF) events. The non-canonical promoter was defined if the ATAC peak is not located in the canonical TSS identified by *ChIPseeker* based on hg38 assembly.

### Colocalization

The Coloc package (v.5.2.3) was used to apply the approximate Bayes factor (ABF) colocalization hypothesis under a single causal variant assumption. To test if isoQTLs contribute to gene expression regulation, we performed colocalization analysis between eIsoforms and their corresponding genes if identified as eGenes in the same cell types in the single-cell eQTL study. In total, we tested 466 eIsoforms out of 293 genes, and the colocalization between isoQTLs and eQTLs was claimed with high confidence by posterior probability of H4 (two traits are associated with the same causal genetic variant) under ABF hypotheses (PP.H4) > 0.7. To test if eIsoforms and lung cancer or lung function share the same causal variant, we performed colocalization analysis of the significant loci in both isoQTL and GWAS and focused on the H4 hypothesis. For lung cancer GWAS, we tested 42 loci (including significant loci from total lung cancer and three predominant histological types: lung adenocarcinoma, lung squamous carcinoma and small cell carcinoma) reported in Byun *et al* multi-ancestry populations ^16^ and 30 loci for lung adenocarcinoma in East Asian population (including significant loci from discovery population and meta-analysis) as well as separate summary statistics combining East Asian discovery and external European populations reported in Shi *et al^17^.* The total number of unique loci from these two studies was 48. For each locus, we searched within the 1MB window of the GWAS lead SNP for significant isoQTLs and listed their associated eIsoforms. Then we performed colocalization using all variants overlapped between lung cancer GWAS and the listed eIsoforms in each cell type. For lung function GWAS, there were 406 significant lead SNPs associated with FEV_1_/FVC and 223 significant lead SNPs with FEV_1_ reported by the meta-analysis of multiple ancestry populations in Shrine *et al* ^26^. We performed colocalization for FEV_1_/FVC and FEV_1_ on the GWAS loci with overlaps with isoQTLs to identify lung-function-associated isoforms, where *p*-values, sample sizes and allele frequencies were used for estimating effect sizes and the variance due to the lack of these data in the downloaded summary stats. The locus are plotted using *locuscomparer* ^84^. The variants with R2 > 0.8 with the lead SNP and nominal p-values < 10-4, or credible set variants from *SuSiE ^85^* were identified as candidate causal variants in the specific locus.

### TWAS

The TWAS analysis was conducted for ancestry-matched East Asian lung adenocarcinoma GWAS summary statistics ^17^ following the documentation of FUSION^86^. First, we used the normalized pseudo-bulked expression matrix used as TensorQTL input to estimate the heritability of isoform expression within each cell type by GCTA-GREML. We then generated prediction models of each isoform based on the *cis*-SNPs (±1MB from TSS) using four models (best linear unbiased prediction, least absolute shrinkage and selection operator or LASSO, elastic-net, and top SNPs) with cross-validation performed for each model. Isoforms with heritability *p*-value < 0.05 and cross-validation *p-*value < 0.05 were retained for TWAS analysis. The imputed isoform expression from prediction models accounting for LD among SNPs calculated using the 1000 Genome EAS population was correlated with lung adenocarcinoma to identify an association between isoform and lung adenocarcinoma. To account for multiple tests, FDR corrections were performed on TWAS *p-*values, and isoforms with FDR < 0.05 were claimed as TWAS significant isoforms associated with lung adenocarcinoma.

### DNA damage assay of *PPIL6* isoforms

#### Cell lines and plasmid constructs

MRC5-SV40 (SV40-immortalized human lung fibroblasts) cells were cultured in standard Dulbecco’s modified Eagle’s Medium (DMEM, Gibco) with 10% fetal bovine serum (Gibco), 2 mM L-glutamine, 100 μg/mL penicillin, and 100 μg/mL streptomycin (Gibco). Cells were incubated at 37°C in a PHCBI incubator with 5% CO□. The cells were authenticated by American Type Culture Collection (ATCC) Short Tandem Repeat (STR) analysis and routinely checked monthly for mycoplasma contamination. Gateway entry clones for the coding sequence of *PPIL6-202*, *PPIL6-207*, and *TALONT003040002* were synthesized (**Supplementary Table 10**), verified, and cloned into pDONR plasmids, followed by subcloning into an N-terminal GFP-tagged vector (pcDNA6.2/N-EmGFP-DEST), with Gateway LR Clonase II Enzyme Mix (Thermo Fisher Scientific, 11791020). The final destination plasmids were sequence-verified by whole-plasmid sequencing at Quintara Biosciences.

#### Transfection

Transfections were performed in 6-well plates by seeding 5 x 10^4^ cells per well. Twenty-four hours after seeding, the cells were transfected using the GenJet In Vitro DNA Transfection Reagent Ver. II (SignaGen, SL100488) and 1 µg of plasmid DNA following the manufacturer’s recommendations. Each plasmid (control, and three isoforms) was transfected in duplicates in each experimental batch. Three independent batches of transfections were performed. After 24 hours of transfection, the culture medium was replaced with a fresh complete growth medium.

#### Flow Cytometric assessment of DNA damage

DNA damage was assessed by quantifying γH2AX and phospho-p53 (p-p53) expression using flow cytometry. Transfected cells were harvested 72 hours post-transfection by trypsinization, fixed with 2% formaldehyde, and permeabilized using 0.05% Triton X-100 (Sigma-Aldrich). Antibody staining was performed using anti–phospho–histone H2A.X (Ser139) antibody (clone JBW301; 1:750; Sigma–Aldrich, 05-636-25UG) and p-p53 (Ser15) mouse monoclonal antibody (clone 16G8; 1:1000; Cell Signaling Technology, 9286), and Goat anti-Mouse IgG (H+L) Cross-Adsorbed Secondary Antibody, Alexa Fluor™ 647 (1:1000 dilution; Invitrogen, A-21235). Cells were acquired and analyzed by the BD LSRII Flow Cytometer in the Texas A&M IBT Flow Core facility. Mock (transfection reagent only)-transfected cells were used to establish gating parameters for identifying GFP^+^ populations as described in Byun et al^16^. Subsequently, GFP-positive cells were gated, and the median p-p53 or γH2AX fluorescence intensities for each overexpression construct were quantified. The median of intensity was further normalized to the GFP-HPS4 control, which is a non-DNA “damage-up” protein and reflects a baseline increase associated with protein overproduction^58^. We did not observe significant variance differences across experimental batches and therefore treated all replicates equally in one-way ANOVA for comparing the difference among the groups (i.e., *HPS4*, *PPIL6-207*, *PPIL6-202*, and *PPIL6-TALONT003040002*). If significant, post hoc Tukey’s test was performed for pair-wise comparison across groups. Significance was claimed as adjusted *p*-value < 0.05 (**Supplementary Table 11**).

## Data availability

The raw sequencing data alongside the final Seurat object and genotype data will be made available on GEO and dbGaP, respectively. All code used to process the long-read isoforms, analyze single-cell data, prepare for isoQTL mapping, integrate with GWAS signals, and produce analysis for figures is available via Zenodo at https://zenodo.org/records/17915287 (ref. ^87^) and GitHub at https://github.com/NCI-ChoiLab/Lung_scLong_read_isoQTL. The GTF file of isoforms identified by long-read sequencing and integrating all single-cell batches is available on Zenodo at https://zenodo.org/records/17915287 (ref. ^87^). The GTF file for specific genes of interest, along with the isoform structures can be searched and downloaded via appshare.cancer.gov/ISOLUTION.

## Acknowledgement

This research was supported by the Intramural Research Program of the National Institutes of Health (NIH). The contributions of the NIH author(s) are considered Works of the United States Government. The findings and conclusions presented in this paper are those of the author(s) and do not necessarily reflect the views of the NIH or the U.S. Department of Health and Human Services. We thank the helpful advice and support of Dr. Pradeep Dagur at the NHLBI Flow Cytometry Core, Kristine Jones and colleagues at the NCI Cancer Genomics Research Laboratory, Dr. George Zaki at Bioinformatics and Computational Science group in Frederick National Laboratory for Cancer Research, NCI Center for Cancer Research Sequencing Facility, and Protein Characterization Laboratory. This work utilized the Biowulf cluster computing system at the NIH.

