## Supplemental Figure 1-21 for "Single-cell full-length transcriptome of human lung reveals genetic effects on isoform regulation beyond gene-level expression"

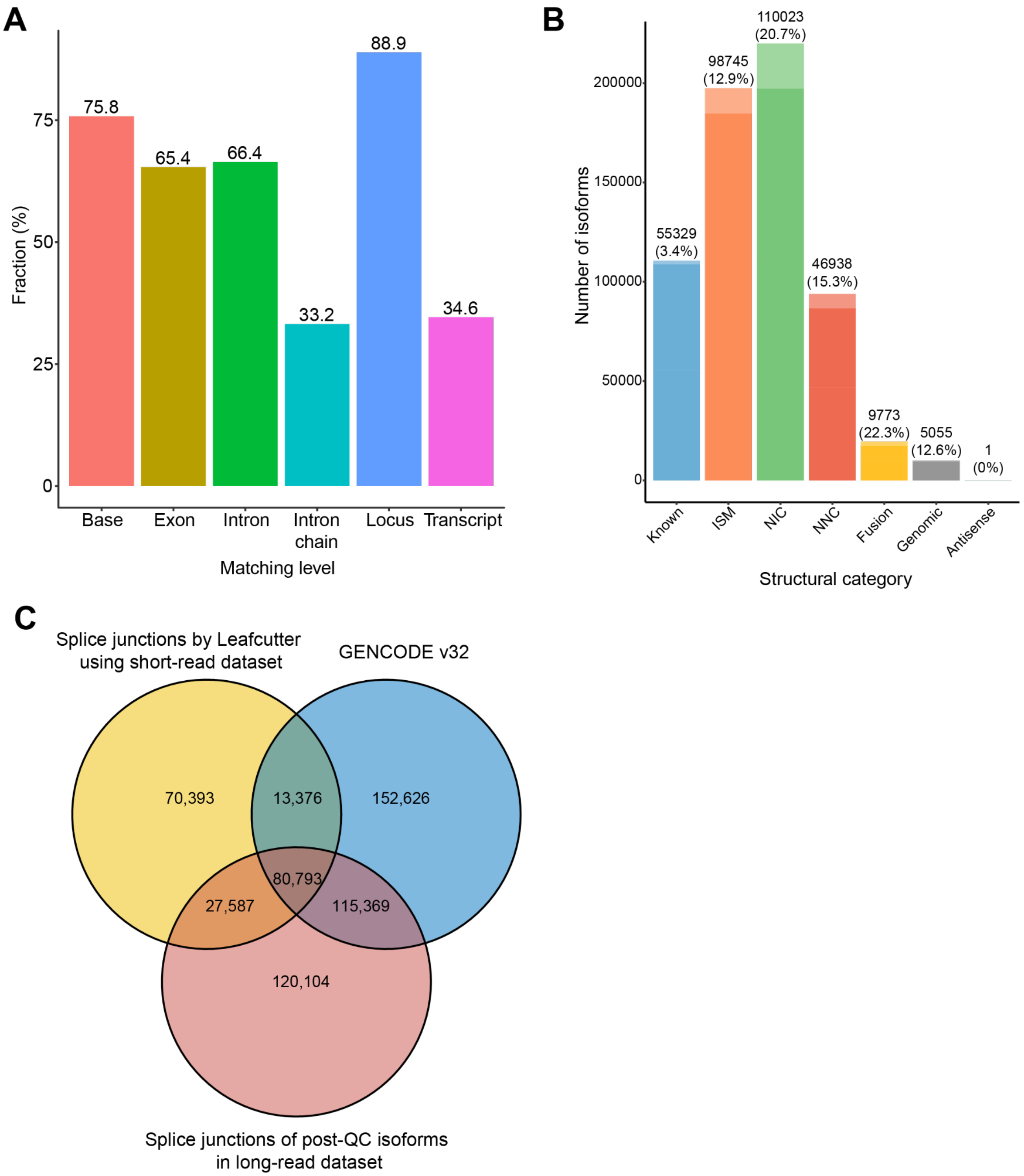


**Supplementary Figure 1. Characteristics of post-QC isoforms**

(**A**) Fraction of post-QC isoforms from our long-read dataset that agreed with the genome reference annotation (i.e., GENCODE v32) at different feature levels, including base, exon, intron, intron chain, transcript, and locus-level matching. Base level means exon bases that are reported at the same coordinate on our long-read dataset and GENCODE v32 (*i.e.*, the overlaps of exons). Exon and intron levels mean the exon/intron intervals with the same exact start-end coordinates. Intron chain level represents that all introns of long-read isoforms match with GENCODE v32. Transcript levels match the overlaps between long-read isoforms and GENCODE v32 isoforms, including isoforms with a single exon, which are not compared at intron-chain level. Locus levels match the cluster of isoforms sharing similar exons between long-read reference and GENCODE v32 reference, where at least one isoform overlapped between the two references if matched. Overall, the intron-chain and transcript levels best summarize our dataset-wide coverage of GENCODE v32 annotated isoforms. (**B**) The fractions of non-sense mediated decay (NMD)-sensitive isoforms among the novel isoforms. The fraction in each structural category is shown in the lighter shades, with the percentage shown on top of the bar plot. SJ: splice junction. (**C**) Venn diagram depicts the overlaps between splice junctions identified by Leafcutter using cell-barcode matched short-read dataset (yellow), splice junctions identified in 325,865 post-QC isoforms in long-read dataset (red), and known splice junctions identified in GENCODE v32 annotation reference.

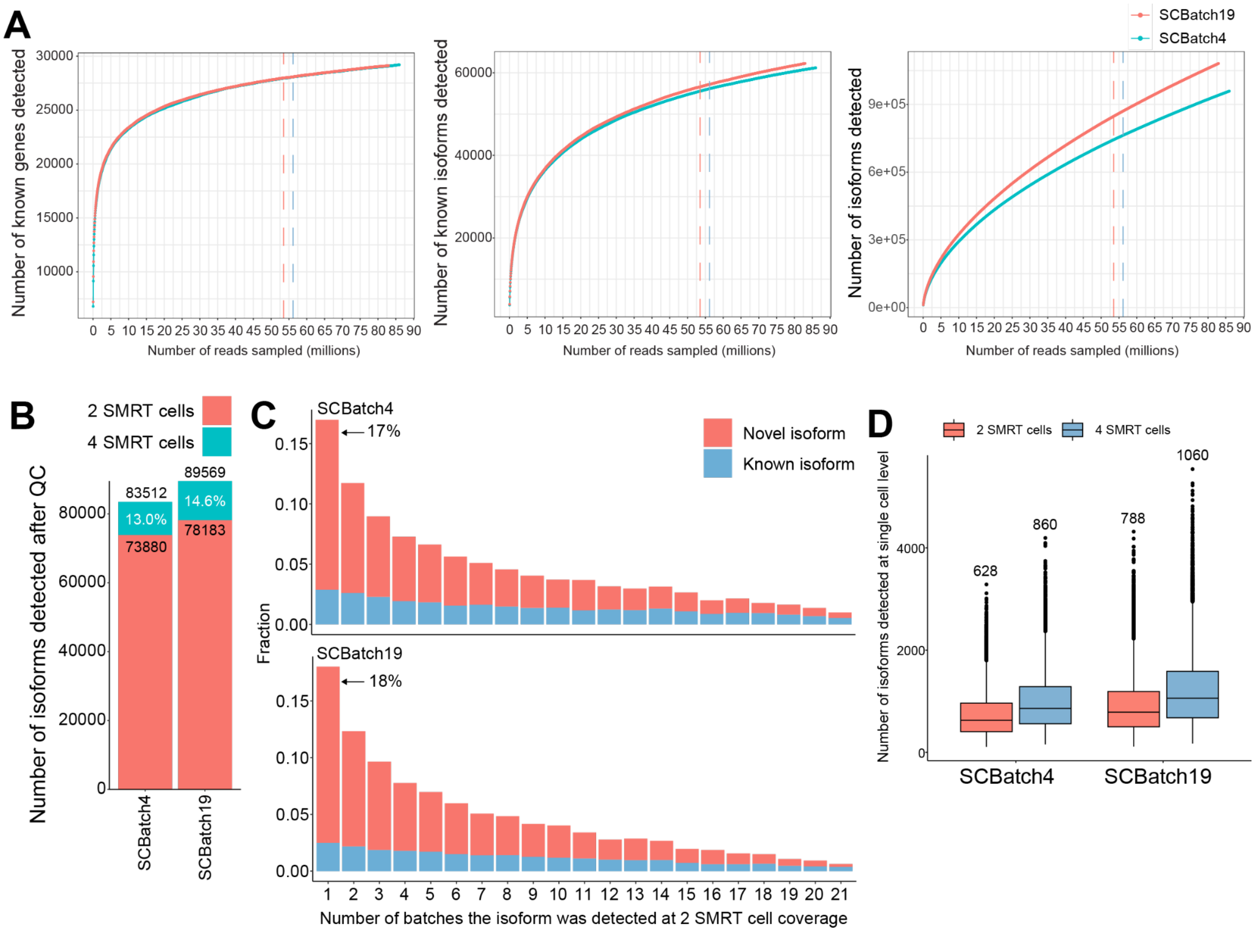


**Supplementary Figure 2. Benchmarking isoform detection with different sequencing depth**

(**A**) Saturation curves of the number of detected known genes (left), known isoforms (middle), and all isoforms (right) by randomly sampling reads. The curves are colored by single cell batches (red: SCBatch4, blue: SCBatch19). The total number of reads using 2 SMRT cells is indicated by the dashed vertical lines (red: SCBatch4, blue: SCBatch19). (**B**) The number of post-QC 325,864 isoforms detected in SCBatch4 and SCBatch19 using 2 or 4 SMRT cells. The bars are colored by different sequencing depths (red: using 2 SMRT cells, blue: using 4 SMRT cells). The percentages of isoforms only detected at 4 SMRT-cell read-depth are indicated. (**C**) The newly detected isoforms after doubling sequencing depth included a high proportion of single-batch-specific isoforms at 2 SMRT cell coverage (upper panel: 17% for SCBatch4, bottom: 18% for SCBatch19). The bar is colored according to the isoform annotation status (blue: known or FSM, red: novel). (**D**) The number of isoforms detected in each cell from sample SCBatch4 and SCBatch19 using 2 or 4 SMRT cells. The median of the number of detected isoforms is labeled. Center lines show the medians; the boxes are colored by sequencing depths (red: using 2 SMRT cells, blue: using 4 SMRT cells) and indicate the middle of 50% of the data; whiskers extend 1.5 times the interquartile range from the 25th and 75th percentiles; outliers are represented by dots.

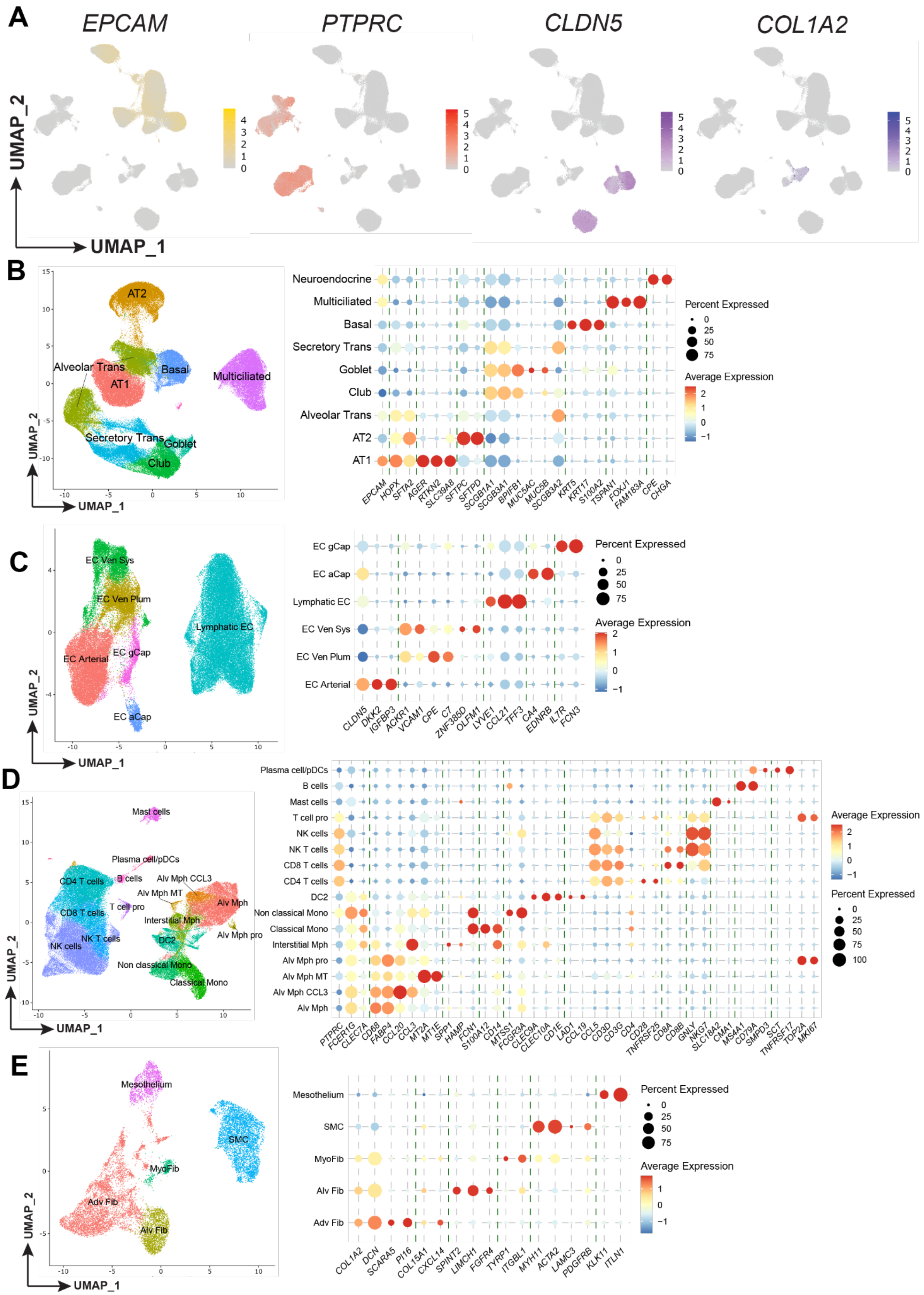
A
EPCAM
PTPRC
CLDN5
COL1A2
UMAP_2
UMAP_1
B
AT2
Alveolar Trans
Basal
AT1
Multiciliated
Secretory Trans
Goblet
Club
Neuroendocrine
Multiciliated
Basal
Secretory Trans
Goblet
Club
Alveolar Trans
AT2
AT1
EPCAM HOPX SFTA2 AGER RTKN2 SLC39A8 SFTPC SFTPD SCGB1A1 SCGB3A1 BPIFB1 MUC5AC MUC5B SCGB3A2 KRT5 KRT17 S100A2 TSPAN1 FOXJ1 FAM183A CPE CHGA
Percent Expressed
Average Expression
C
EC Ven Sys
EC Ven Plum
Lymphatic EC
EC Arterial
EC gCap
EC aCap
EC gCap
EC aCap
Lymphatic EC
EC Ven Sys
EC Ven Plum
EC Arterial
CLDN5 DKK2 IGFBP3 ACKR1 VCAM1 CPE C7 ZNF385D OLFM1 LYVE1 CCL21 TFF3 CA4 EDNRB IL7R FCN3
Percent Expressed
Average Expression
D
Mast cells
Plasma cell/pDCs
B cells
Alv Mph CCL3
CD4 T cells
Alv Mph MT
Alv Mph
T cell pro
CD8 T cells
Interstitial Mph
Alv Mph pro
NK T cells
NK cells
DC2
Non classical Mono
Classical Mono
Plasma cell/pDCs
B cells
Mast cells
T cell pro
NK cells
NK T cells
CD8 T cells
CD4 T cells
DC2
Non classical Mono
Classical Mono
Interstitial Mph
Alv Mph pro
Alv Mph MT
Alv Mph CCL3
Alv Mph
PTPRC FCER1G CLEC7A CD68 FABP4 CCL20 CCL3 MT2A MT1E SPP1 HAMP FCN1 S100A12 CD14 MTSS1 FCGR3A CLEC9A CLEC10A CD1E LAD1 CCL19 CCL5 CD3D CD3G CD4 CD28 TNFRSF25 CD8A CD8B GNLY NKG7 SLC18A2 CMA1 MS4A1 CD79A SMPD3 SCT TNFRSF17 TOP2A MKI67
Average Expression
Percent Expressed
E
Mesothelium
SMC
MyoFib
Adv Fib
Alv Fib
Mesothelium
SMC
MyoFib
Alv Fib
Adv Fib
COL1A2 DCN SCARA5 PI16 COL15A1 CXCL14 SPINT2 LIMCH1 FGFR4 TYRP1 ITGBL1 MYH11 ACTA2 LAMC3 PDGFRB KLK11 ITLN1
Percent Expressed
Average Expression

**Supplementary Figure 3. Iterative clustering and cell type annotation of single-cell long-read data**

(**A**) Canonical markers for epithelial (*EPCAM*), immune (*PTPRC* encoding *CD45*), endothelial (*CLDN5*), and stromal cells (*COL1A2*) are superimposed on the UMAP. UMAP plots depict re-clustering within epithelial cells (**B**), endothelial cells (**C**), immune cells (**D**) and stromal cells (**E**) in the left panels. The expression of marker genes used for annotation is shown in the right panels. Sets of genes used together for distinguishing specific cell subtypes were grouped as indicated by dashed green lines. For example, in epithelial cells, *SCGB1A1*, *SCGB3A1*, *BPIFB1*, *MUC5AC*, *MUC5B,* and *SCGB3A2* were used to distinguish transitional cells and secretory cells. Specifically, *SCGB1A1* and *SCGB3A1* were used to label secretory cells, and *SCGB3A2*$^{high}$ cells were considered as transitional cells, thus *SCGB1A1*$^{+}$*SCGB3A1*$^{+}$*SCGB3A2*$^{high}$ cells were annotated as secretory transitional cells. *BPIFB1*, *MUC5AC,* and *MUC5B* were reported to be more specific to goblet cells, thus *BPIFB1*$^{+}$*MUC5AC*$^{+}$*MUC5B*$^{+}$*SCGB3A2*$^{low}$ cells were annotated as goblet, and the remaining cluster of secretory cells was annotated as club cells. Additionally, the other *SCGB3A2*$^{high}$ cells were annotated as alveolar transitional cells due to the expression of alveolar marker genes (i.e., *HOPX* and *SFTA2*).

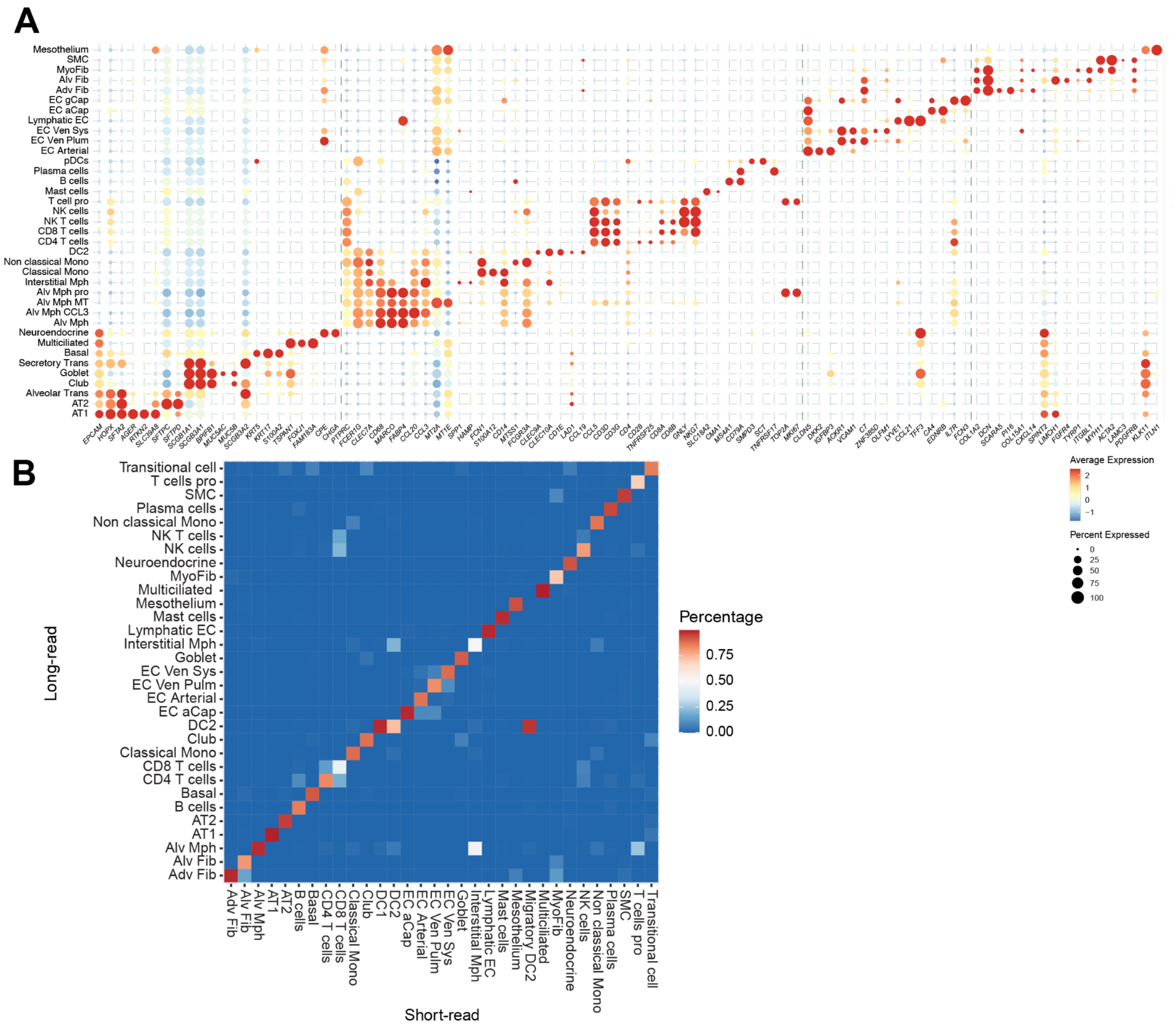


**Supplementary Figure 4. Comparison of cell type annotation between long-read and short-read data**

(**A**) Dot plot depicts gene expression levels and percentage of cells expressing marker genes used for cell type annotation. (**B**) Comparison of cell type annotation for barcode-matched cells in our short-read and long-read datasets. The color in the heat map represents the proportion of cells annotated in long-read data for cells in each cell type of short-read data. A few rare cell subtypes, such as peribronchial fibroblasts, were not found in the long-read data, which might be in part due to a lower long-read sequencing depth limiting the detection of cell-type signature genes.

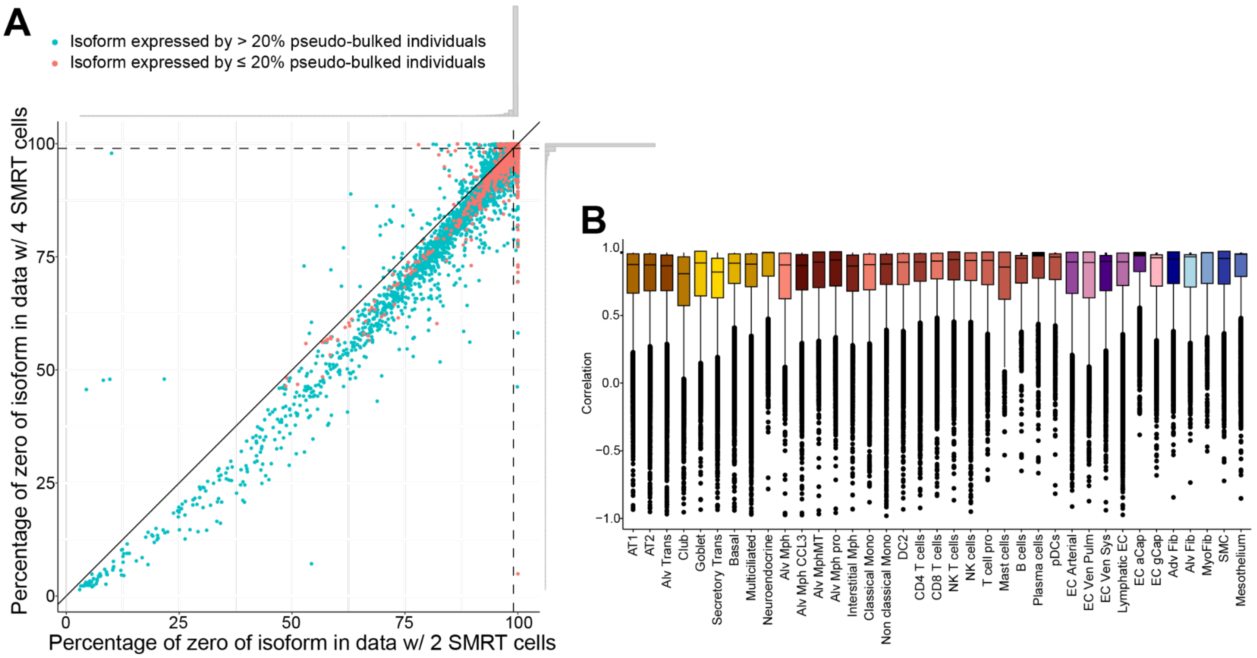


**Supplementary Figure 5. Criteria for including isoforms for DEI and isoQTL analyses addressing sparsity of isoform expression**

(**A**) Sparsity of isoform expression (*i.e.*, percentage of cells with a 0 read for the isoform across all cells). Each dot represents an isoform and is colored blue if the isoform is expressed by more than 20% of individuals after pseudo-bulking or red if expressed in less than 20% of individuals. The marginal histograms alongside the axis show the distribution of sparsity of isoform expression. At the single-cell level, the isoform expression profiles were extremely sparse, with only 3.3% of isoforms being expressed by more than 1% of cells. At an increased sequencing depth up to 4 SMRT cells, the sparsity was only modestly reduced, with 4.4% of isoforms being expressed by more than 1% of cells. The isoforms passing the filtering criteria for our main analyses (blue dots) displayed relative abundance in higher proportions of cells. (**B**) The box plots show the correlation of the isoform expression at the individual level within each cell type between 2 SMRT cell and 4 SMRT cell sequencing depths. Each dot represents the isoform expressed by > 20% of individuals and tested for DEI. By the individual level correlation, the isoform expressions based on the sequencing depth of 2 SMRT cells showed a high correlation with those of 4 SMRT cells, indicating the robustness of isoform expression at the individual level for our sequencing depth. Center lines show the medians; the boxes are colored by cell types and indicate the middle of 50% of the data; whiskers extend 1.5 times the interquartile range from the 25th and 75th percentiles; outliers are represented by dots

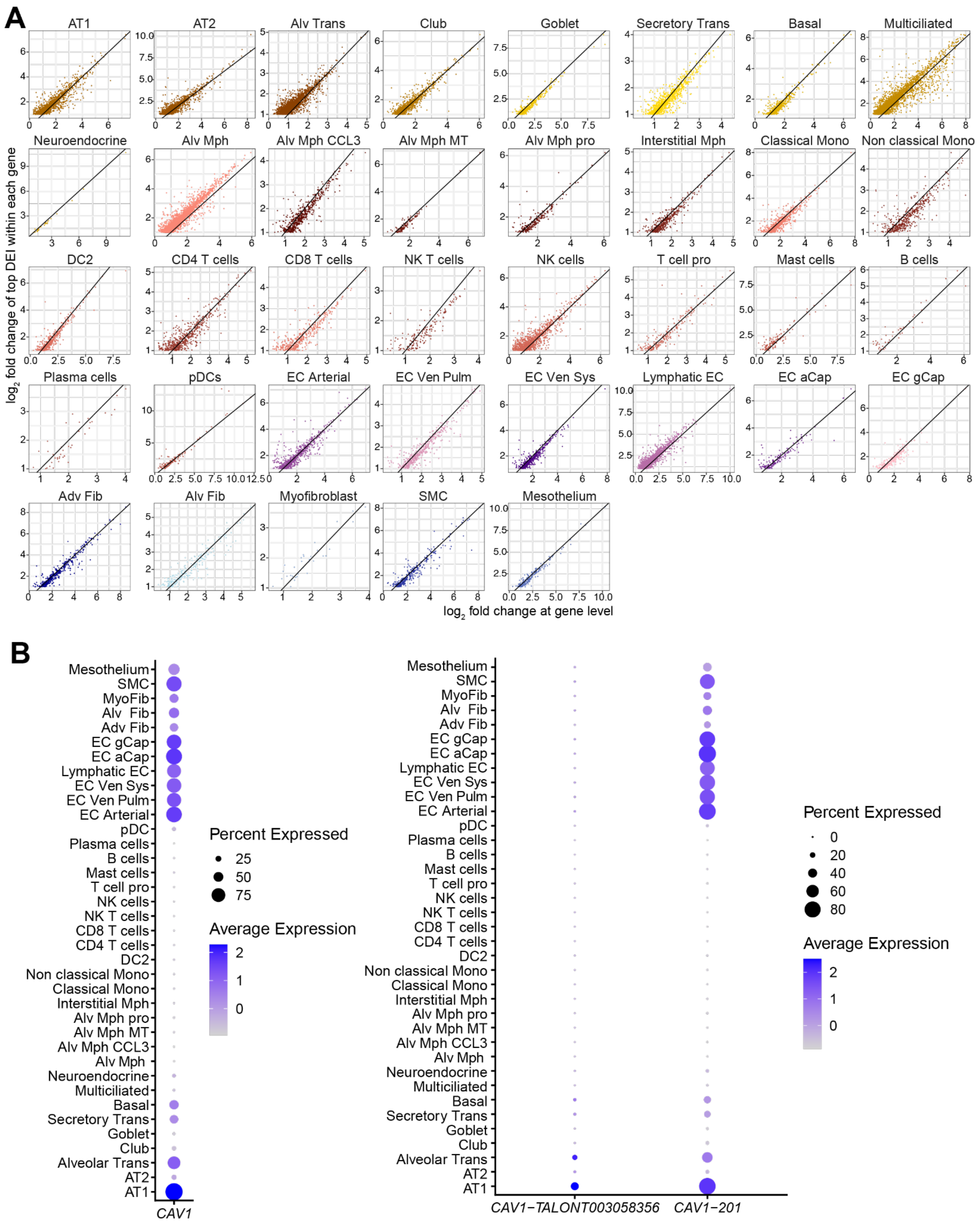


**Supplementary Figure 6. Comparisons between DEIs and DEGs**

(**A**) Comparison of $\log_2$ fold changes of DEIs with $\log_2$ fold change of the corresponding genes with adjusted p value < 0.05 from the gene-level differential expression analysis for each cell type. Each dot represents a DEI, and the black line is diagonal. The DEI with the largest fold change was matched to the gene in the cases of multiple DEIs for a gene. The isoforms over the diagonal lines indicate potentially higher specificity than gene levels for each cell type. **(B**) Dot plot depicts expression levels and percentages of cells expressing *CAV1* at the gene level (left panel) and *CAV1-TALONT003058356* and *CAV1-201* (right panel) at the isoform level.

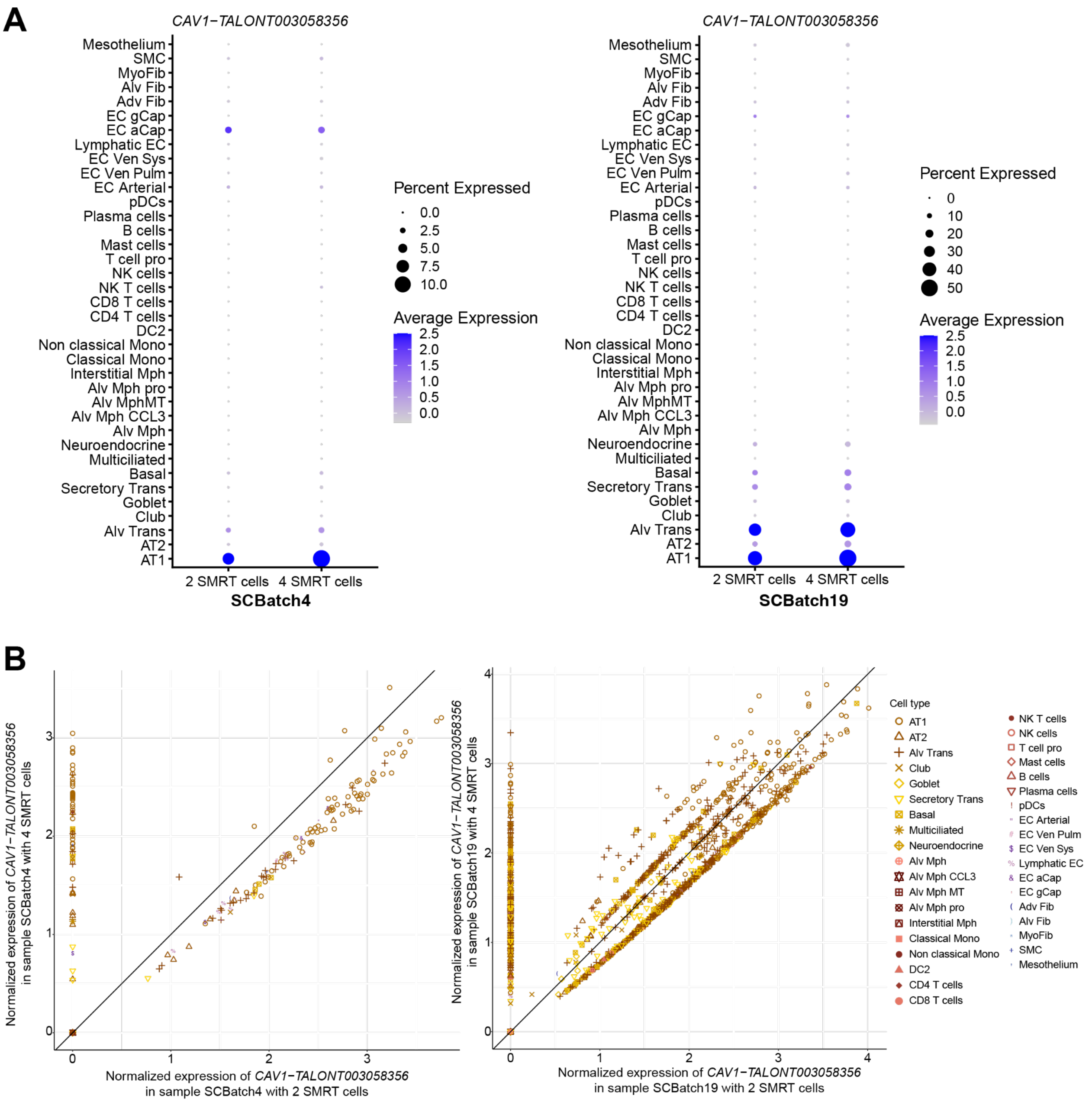


**Supplementary Figure 7. Effect of sequencing depth on AT1 marker gene DEI, *CAV1-TALONT003058356***

(**A**) Dot plot depicts expression levels and percentages of cells expressing *CAV1-TALONT003058356* in batch SCBatch4 (left panel) and SCBatch19 (right panel) using 2 or 4 SMRT cells. The proportion of cells expressing this isoform increased mostly in AT1 and alveolar transitional cells but not in any other cell types, maintaining cell type specificity at a higher coverage. (**B**) Normalized expression of *CAV1-TALONT003058356* from each cell in batch SCBatch4 and SCBatch19 using 2 or 4 SMRT cells. Each point represents a single cell and is colored and shaped by cell type annotation. The cells expressing *CAV1-TALONT003058356* only at the 4 SMRT cell sequencing depth (initially not detected in 2 SMRT cell sequencing depth) are composed of 72.4% AT1 and 61.0% alveolar transitional cells (yellow circle and yellow plus sign, respectively), demonstrating that the cell type specificity of this novel isoform is not mainly due to the sparsity and shallow sequencing depth.

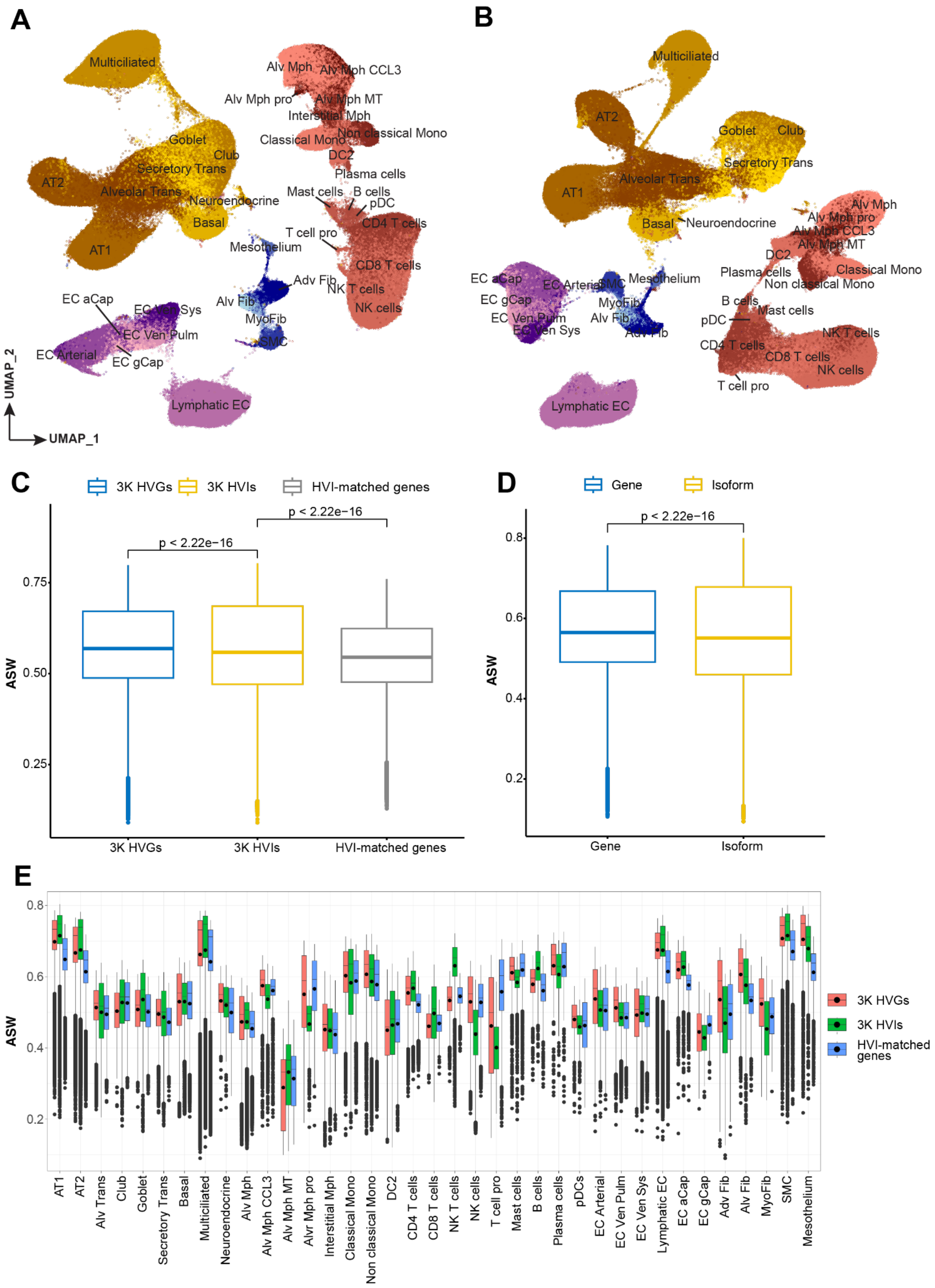
A
B
C
D
E
UMAP_1
UMAP_2
3K HVGs
3K HVIs
HVI-matched genes
Gene
Isoform
p < 2.22e−16
ASW

**Supplementary Figure 8. Benchmarking cell clustering and annotation using gene or isoform expression signatures**

(**A**) UMAP plot depicts the clustering of cells using 3,000 highly variable genes (HVGs) labelled with cell type annotation of iterative clustering (details in Methods). (**B**) UMAP plot depicts the clustering of cells using 3,000 highly variable isoforms (HVIs) labelled with the same cell type annotation of iterative clustering. To assess the performance of clustering contributing to cell type annotation, we calculated the silhouette width of each cell (*i.e.*, relative closeness to the center of its cluster compared to that of the closest neighboring cluster) using the cell type annotation from iterative clustering as ground truth. While comparing gene-level and isoform-level results, a paired T-test was used to calculate *p*-value. Note that the ground truth cell type annotation is from the iterative clustering within each of the 4 lung cell categories, whereas the benchmarking clustering was performed at the whole dataset level. (**C**) Comparison of the average silhouette width (ASW) of cells using different sets of highly variable features. The comparison between HVGs and HVIs was performed using an equal number of unsupervised highly variable features (3,000), where multiple isoforms could come from the same gene. Therefore, 3,000 HVIs had a smaller number of total genes; thus, we benchmarked the performance between 3K HVIs and their matching genes to compare the same gene sets. Using 3,000 HVIs resulted in a significantly higher ASW than 2,532 matched genes, indicating that additional isoforms for the same set of genes could provide distinct cell type information for clustering. (**D**) Comparison of ASW of cells using overlaps between 3,000 HVGs and 3,000 HVIs at the gene-level. 1,090 overlapped genes were selected and used for gene-level clustering, and the top-ranked variable isoforms for these genes (1,090 HVIs) were used for isoform-level clustering. By comparing these two conditions, we evaluated the performance of gene-level and isoform-level clustering using an equal number of variables. Each data point in the box plot represents the ASW for each cell, and *p* values were calculated by paired T test. (**E**) Comparison of the ASW of cells using different sets of highly variable features across cell types. Among them, eight cell types (AT1, AT2, Club, Goblet, Multiciliated, T cells, lymphatic ECs, and SMCs), isoform signatures outperformed gene-level clustering with a significantly higher ASW Center lines show the medians; the boxes indicate the middle of 50% of data, whereas the black dots in the boxes show the means; whiskers extend 1.5 times the interquartile range from the 25th and 75th percentiles; outliers are represented by dots.

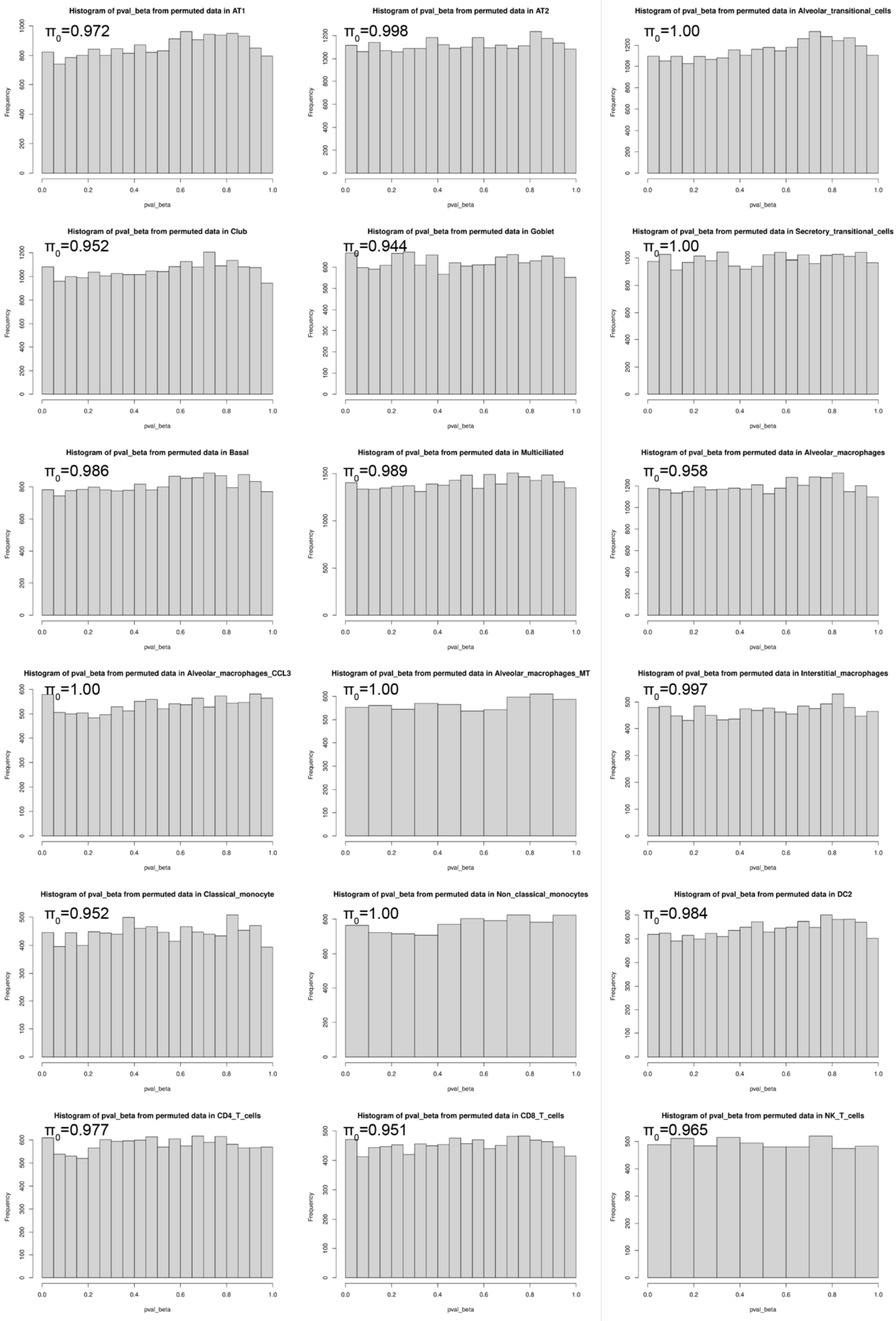
Histogram of pval_beta from permuted data in AT1
π0=0.972
Histogram of pval_beta from permuted data in AT2
π0=0.998
Histogram of pval_beta from permuted data in Alveolar_transitional_cells
π0=1.00
Histogram of pval_beta from permuted data in Club
π0=0.952
Histogram of pval_beta from permuted data in Goblet
π0=0.944
Histogram of pval_beta from permuted data in Secretory_transitional_cells
π0=1.00
Histogram of pval_beta from permuted data in Basal
π0=0.986
Histogram of pval_beta from permuted data in Multiciliated
π0=0.989
Histogram of pval_beta from permuted data in Alveolar_macrophages
π0=0.958
Histogram of pval_beta from permuted data in Alveolar_macrophages_CCL3
π0=1.00
Histogram of pval_beta from permuted data in Alveolar_macrophages_MT
π0=1.00
Histogram of pval_beta from permuted data in Interstitial_macrophages
π0=0.997
Histogram of pval_beta from permuted data in Classical_monocyte
π0=0.952
Histogram of pval_beta from permuted data in Non_classical_monocytes
π0=1.00
Histogram of pval_beta from permuted data in DC2
π0=0.984
Histogram of pval_beta from permuted data in CD4_T_cells
π0=0.977
Histogram of pval_beta from permuted data in CD8_T_cells
π0=0.951
Histogram of pval_beta from permuted data in NK_T_cells
π0=0.965
Frequency
pval_beta

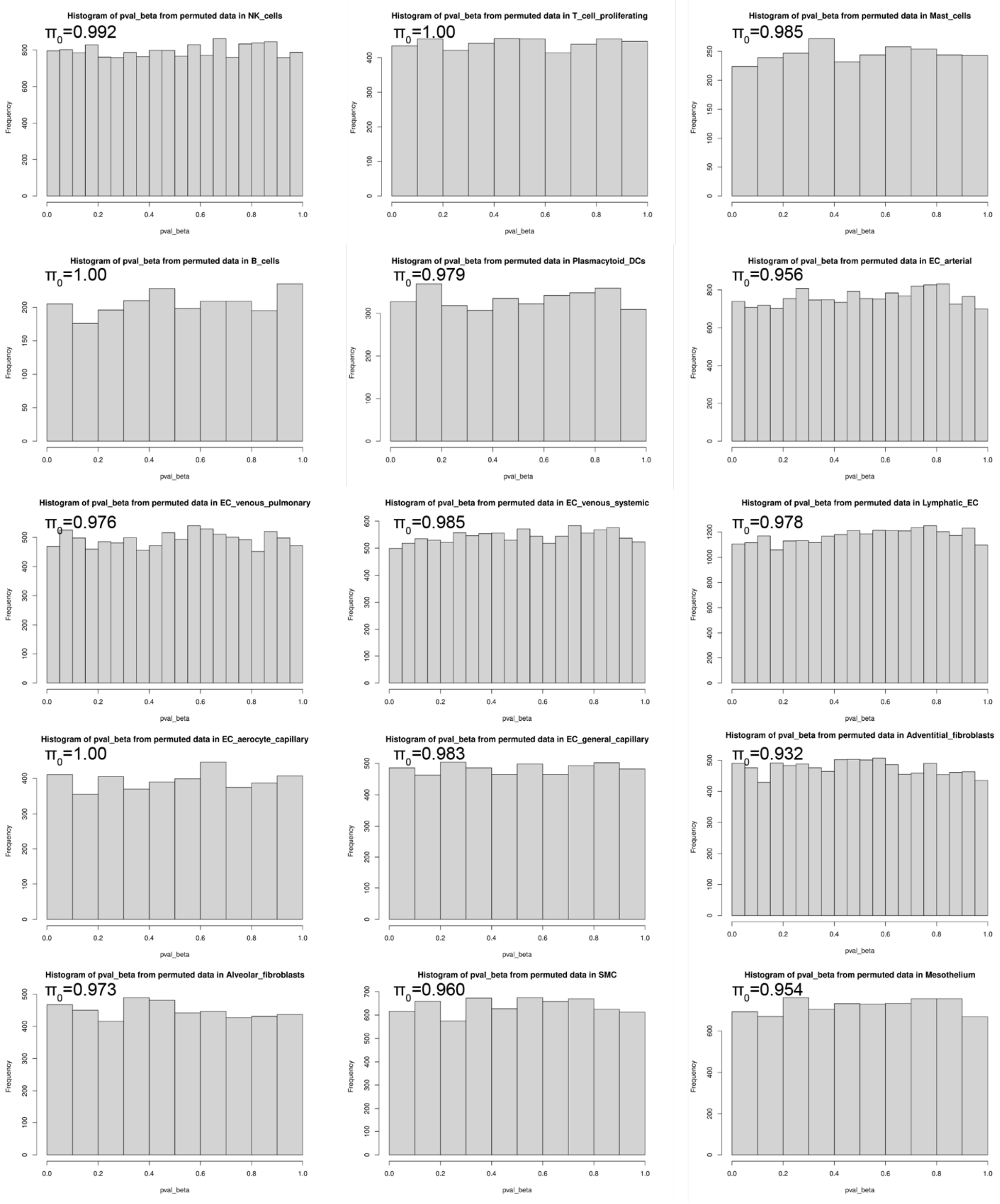


**Supplementary Figure 9. Type I error control for isoQTL detection using negative binomial models**

Histograms of p values adjusted by beta approximation of top variants for each tested gene in permuted data of 33 lung cell types. The permuted data were generated by shuffling the count expression of each isoform to break the association, which is supposed to be null hypotheses. As shown in the histograms, the p values from permuted are distributed uniformly with the proportion of true null hypotheses $\pi_0$ calculated, indicating the p values were well calibrated in negative binomial models.

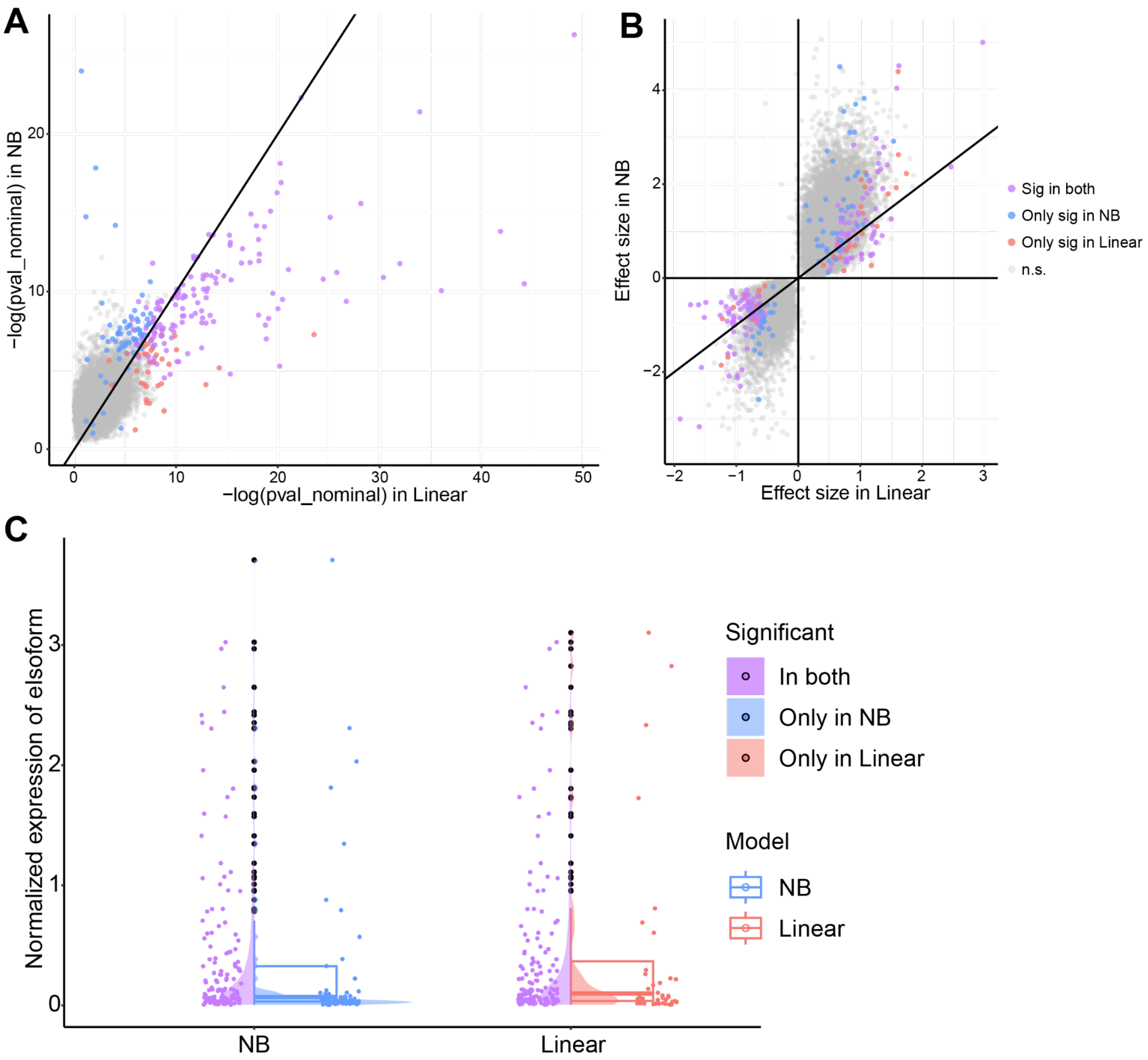


**Supplementary Figure 10. Comparison of isoQTL summary statistics between linear model and negative binomial model for AT2 cells**

Comparison of nominal p values (**A**) and effect sizes (**B**) between the negative binomial (NB) model of jaxQTL and the linear model of TensorQTL in AT2 cells. Each dot represents an isoform with its top SNP color-coded if claimed as significant eIsoforms after multiple test corrections in the two models (blue: only significant in NB model, red: only significant in linear model, purple: significant in both models). (**C**) Distribution of the expression levels of eIsoforms identified by NB model, linear model, and both models. The box plots show the expression of all eIsoforms identified by NB (blue, 210 eIsoforms) and linear (red, 180 eIsoforms), where the median of eIsoform expression is smaller in NB models. The violin plots are split between the eIsoforms detected in both models (left half, purple) and those only detected in the specific model (right half, blue or red). The exclusively identified eIsoforms of NB models have relatively lower expression levels, shown as the blue peak closer to the baseline than the red peak.

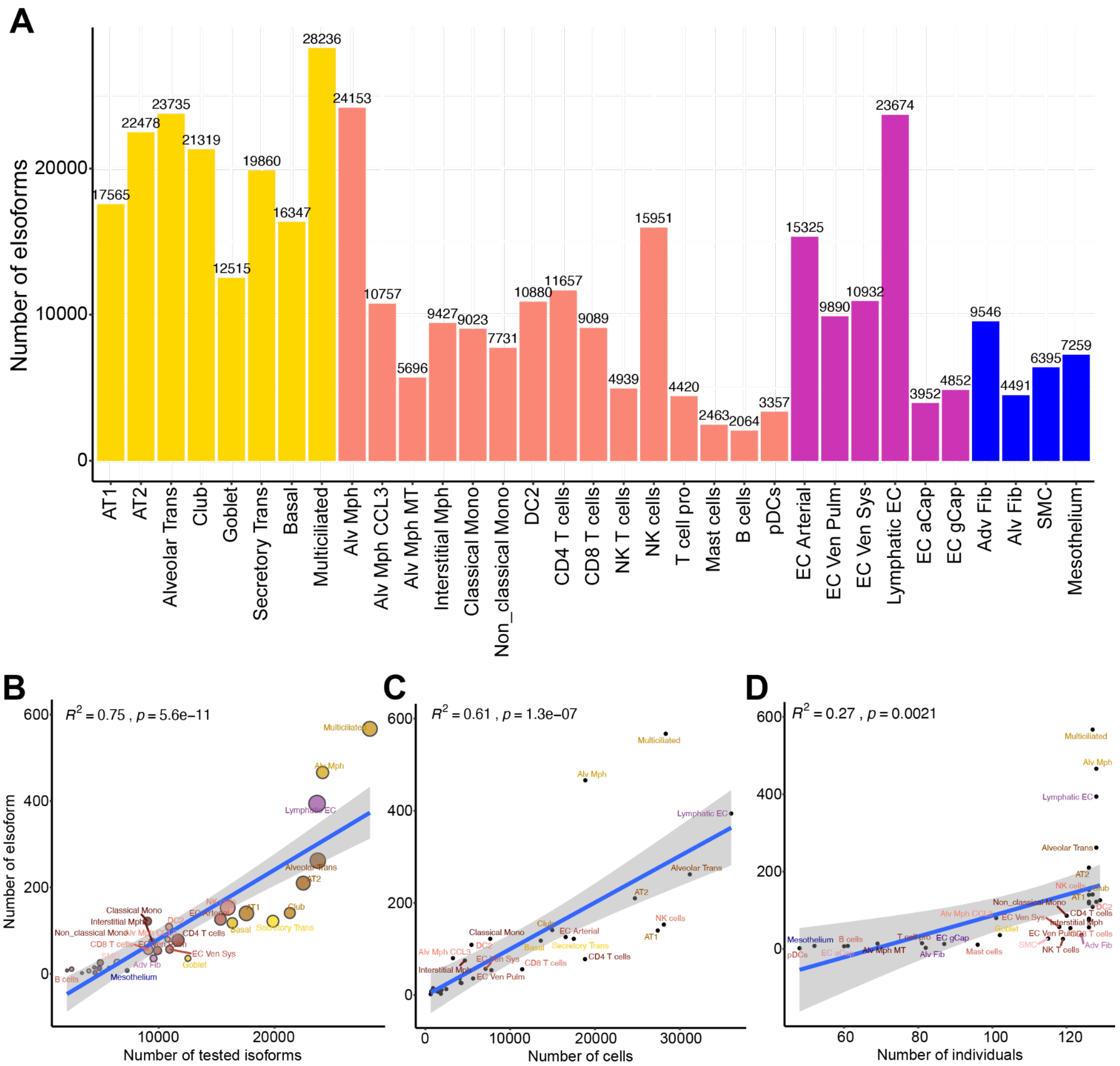


**Supplementary Figure 11. Isoform selection for QTL mapping and isoQTL detection power across cell types**

(**A**) Number of isoforms that were tested in isoQTL analysis across cell types. The bars are colored by cell category (yellow for epithelial, red for immune, purple for endothelial, and blue for stromal). (**B**) Correlation between the number of tested isoforms and the number of eIsoforms across cell types. The bubbles are colored by cell types, and the size represents the number of cells in that cell type. (**C**) Correlation between the numbers of cells and the numbers of eIsoforms across cell types. (**D**) Correlation between the numbers of individuals and the numbers of eIsoforms across cell types. $R^2$ and statistical significance were tested by Pearson correlation.

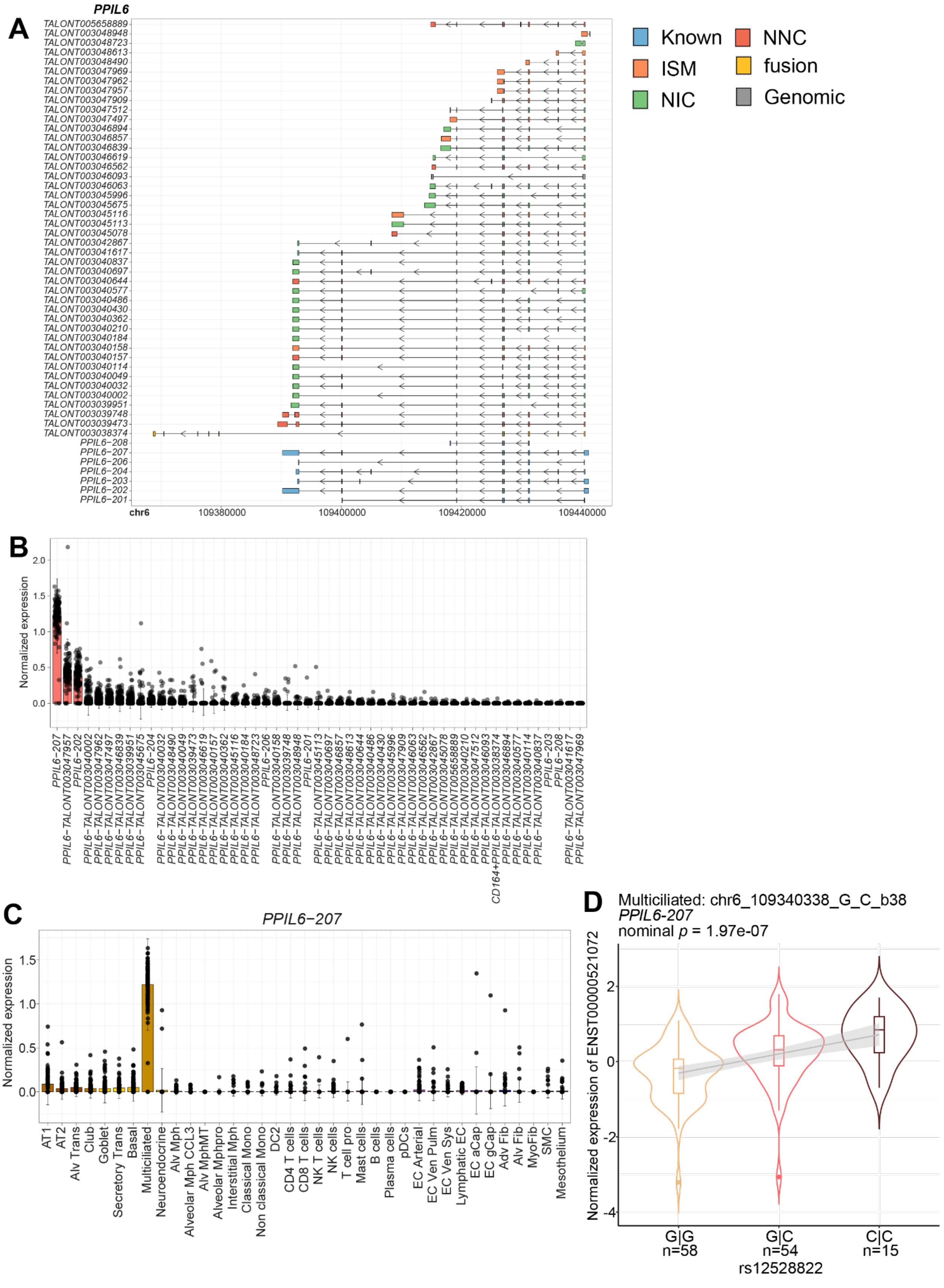

A
PPIL6
TALONT005658889
TALONT003048948
TALONT003048723
TALONT003048613
TALONT003048490
TALONT003047969
TALONT003047962
TALONT003047957
TALONT003047909
TALONT003047512
TALONT003047497
TALONT003046894
TALONT003046857
TALONT003046839
TALONT003046619
TALONT003046562
TALONT003046093
TALONT003046063
TALONT003045996
TALONT003045675
TALONT003045116
TALONT003045113
TALONT003045078
TALONT003042867
TALONT003041617
TALONT003040837
TALONT003040697
TALONT003040644
TALONT003040577
TALONT003040486
TALONT003040430
TALONT003040362
TALONT003040210
TALONT003040184
TALONT003040158
TALONT003040157
TALONT003040114
TALONT003040049
TALONT003040032
TALONT003040002
TALONT003039951
TALONT003039748
TALONT003039473
TALONT003038374
PPIL6-208
PPIL6-207
PPIL6-206
PPIL6-204
PPIL6-203
PPIL6-202
PPIL6-201
chr6
109380000
109400000
109420000
109440000
Known
ISM
NIC
NNC
fusion
Genomic
B
Normalized expression
2.0
1.5
1.0
0.5
0.0
C
PPIL6−207
Normalized expression
1.5
1.0
0.5
0.0
AT1
AT2
Alv Trans
Club
Goblet
Secretory Trans
Basal
Multiciliated
Neuroendocrine
Alv Mph
Alveolar Mph CCL3
Alv MphMT
Alveolar Mphpro
Interstitial Mph
Classical Mono
Non classical Mono
DC2
CD4 T cells
CD8 T cells
NK T cells
NK cells
T cell pro
Mast cells
B cells
Plasma cells
pDCs
EC Arterial
EC Ven Pulm
EC Ven Sys
Lymphatic EC
EC aCap
EC gCap
Adv Fib
Alv Fib
MyoFib
SMC
Mesothelium
D
Multiciliated: chr6_109340338_G_C_b38
PPIL6-207
nominal p = 1.97e-07
Normalized expression of ENST00000521072
2
0
-2
-4
G|G
n=58
G|C
n=54
C|C
n=15
rs12528822

**Supplementary Figure 12. Expression and isoQTL of *PPIL6-207***

(**A**) Structure of all *PPIL6* isoforms based on the genomic coordinates. Arrows indicate the direction of transcription. Isoform categories are color-coded. (**B**) Normalized expressions of all post-QC isoforms for *PPIL6* in multiciliated cells. The isoforms were ordered by average expression level across individuals, indicating 207 was the most abundant isoform for *PPIL6* in multiciliated cells.The bar shows the mean of normalized expression. Each dot represents the isoform expression of each individual within each cell type. The error bars show the mean of normalized expression ± standard deviation. (**C**) The normalized expressions of *PPIL6-207* at the individual level across lung cell types. Each dot represents an individual. The bar shows the mean of normalized expression. Each dot represents the isoform expression of each individual within each cell type. The error bars show the mean of normalized expression ± standard deviation. (**D**) Association between normalized expression of *PPIL6-207* and the genotype of lead isoQTL rs12528822.The violins and boxes are colored by genotype, and the grey line shows the trend of association. The lead isoQTL is shown at the top with chromosome, position, reference allele and alternative allele based on hg38 assembly. Center lines show the medians; the box indicates the middle of 50% of data; whiskers extend 1.5 times the interquartile range from the 25th and 75th percentiles; outliers are represented by dots; density of normalized expression is represented by the width of violin shape.

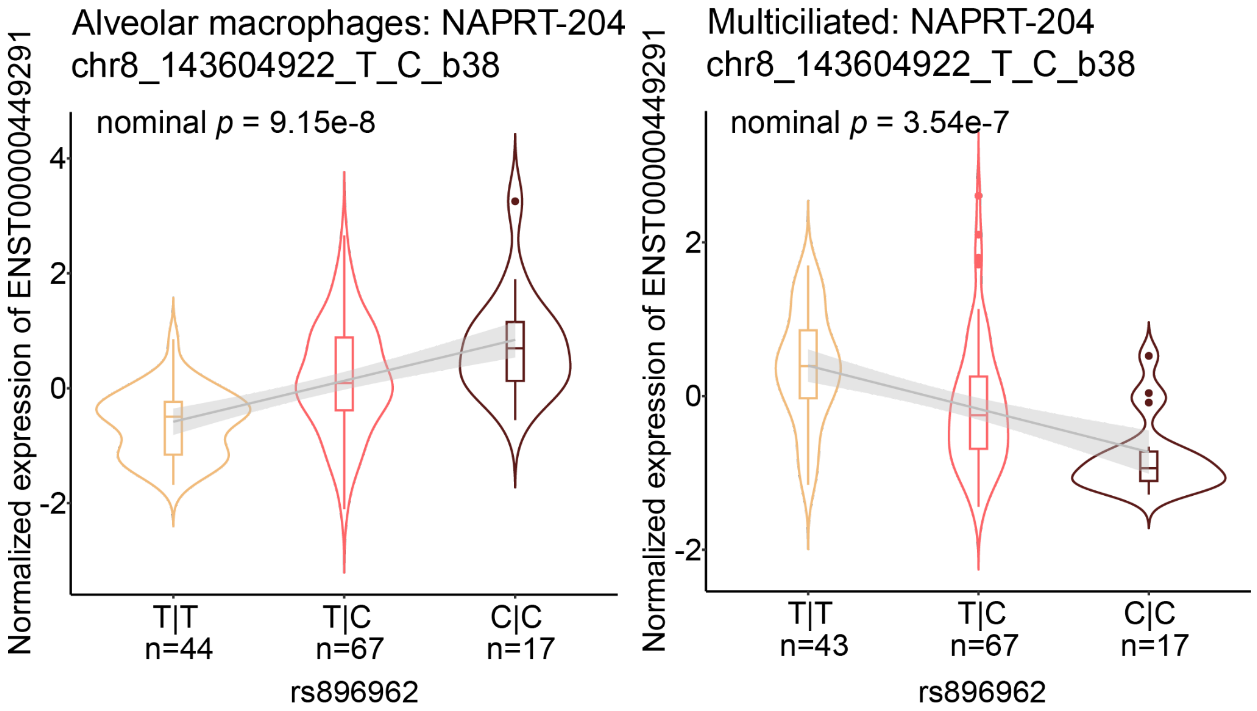


**Supplementary Figure 13. Opposite allelic effects of isoQTLs in different cell types**

Association between normalized expression of *NAPRT-204* and the genotype of lead isoQTL rs896962 in two lung cell types. The violins and boxes are colored by genotype, and the grey line shows the trend of association. The lead isoQTL is shown at the top, with chromosome, position, reference allele, and alternative allele based on hg38 assembly. Center lines show the medians; the box indicates the middle of 50% of data; whiskers extend 1.5 times the interquartile range from the 25th and 75th percentiles; outliers are represented by dots; density of normalized expression is represented by the width of violin shape.

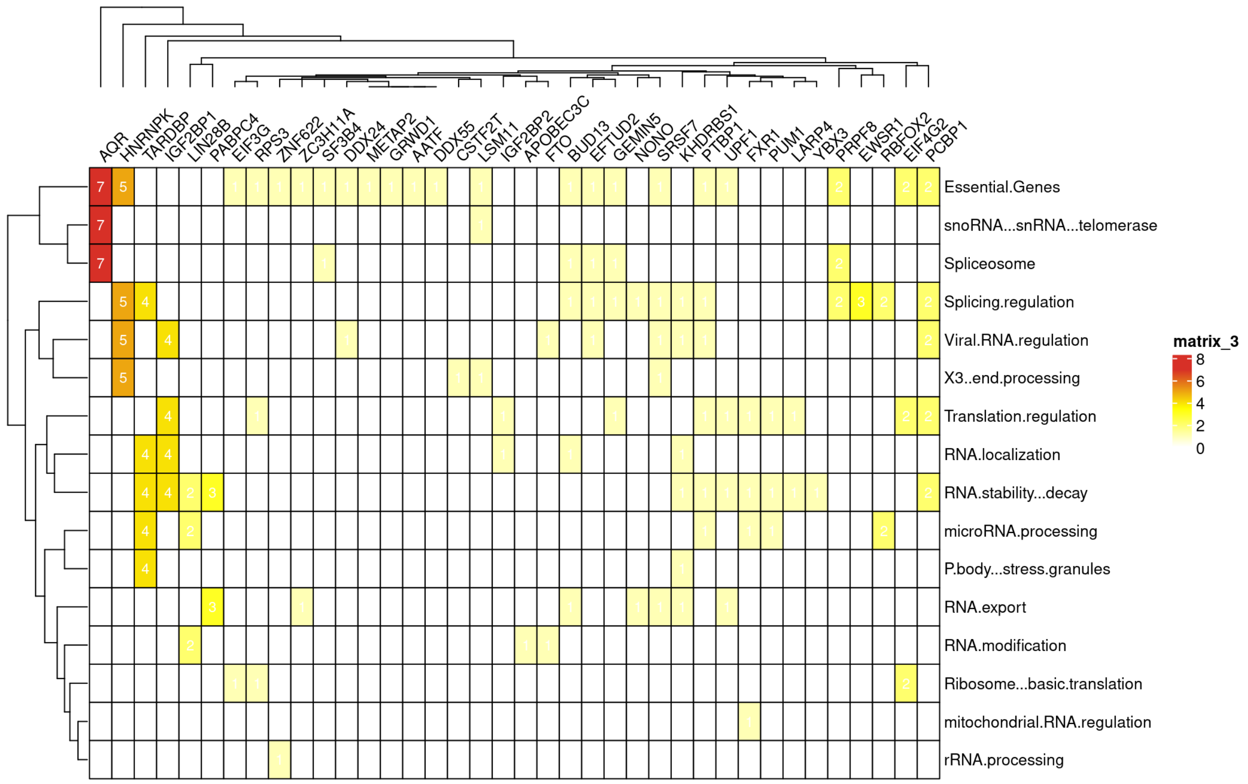


**Supplementary Figure 14. isoQTLs overlapping RBP motifs**
Number of LD-pruned isoQTLs located in the motifs of each RBP in relevant functional annotation categories. The color in the heat map shows the number of LD-pruned isoQTLs.

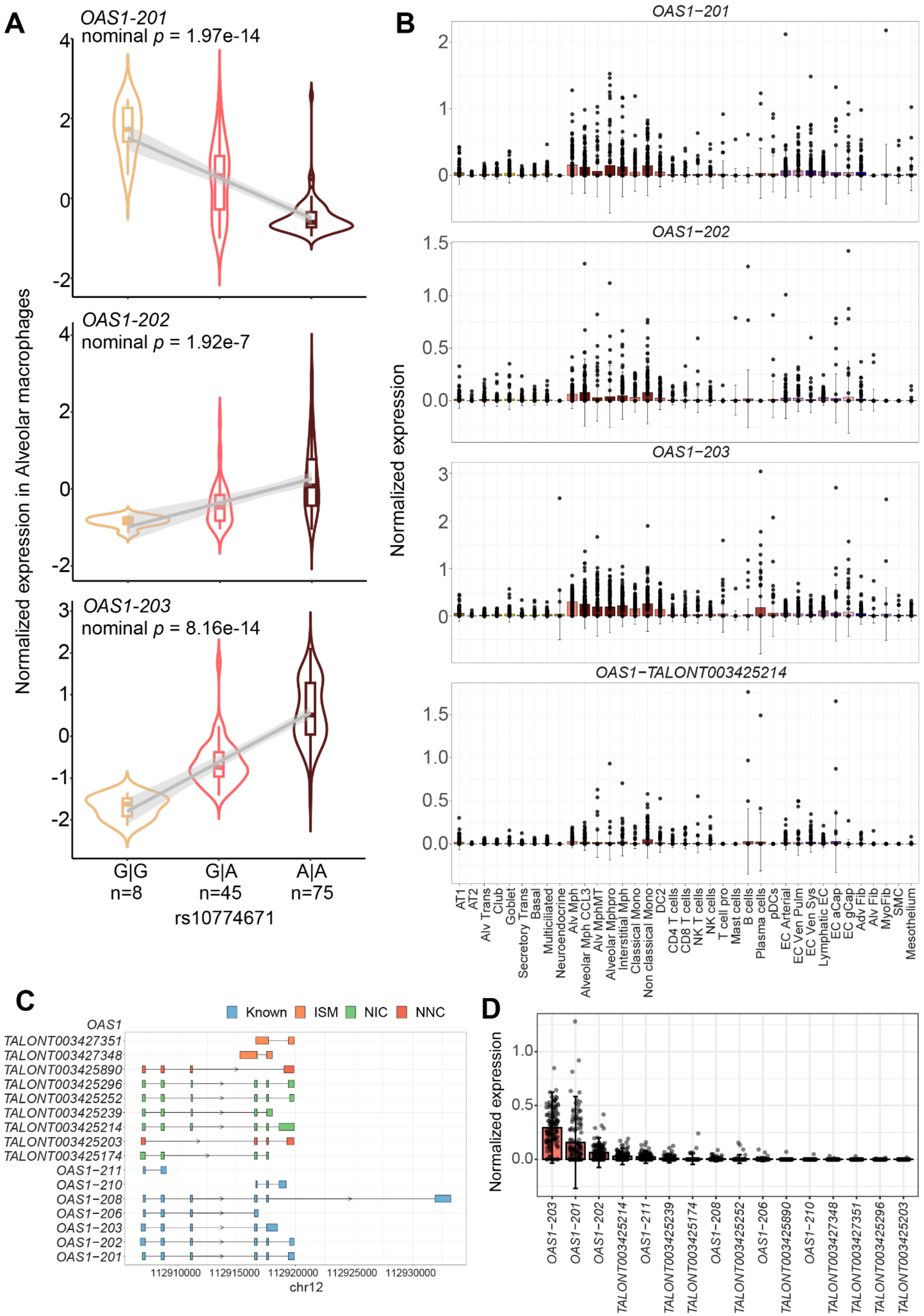
A
B
C
D
OAS1-201
nominal p = 1.97e-14
OAS1-202
nominal p = 1.92e-7
OAS1-203
nominal p = 8.16e-14
Normalized expression in Alveolar macrophages
G|G n=8
G|A n=45
A|A n=75
rs10774671
Normalized expression
OAS1−201
OAS1−202
OAS1−203
OAS1−TALONT003425214
AT1
AT2
Alv Trans
Club
Goblet
Secretory Trans
Basal
Multiciliated
Neuroendocrine
Alv Mph
Alveolar Mph CCL3
Alv MphMT
Alveolar Mphpro
Interstitial Mph
Classical Mono
Non classical Mono
DC2
CD4 T cells
CD8 T cells
NK T cells
NK cells
T cell pro
Mast cells
B cells
Plasma cells
pDCs
EC Arterial
EC Ven Pulm
EC Ven Sys
Lymphatic EC
EC aCap
EC gCap
Adv Fib
Alv Fib
MyoFib
SMC
Mesothelium
Known
ISM
NIC
NNC
OAS1
TALONT003427351
TALONT003427348
TALONT003425890
TALONT003425296
TALONT003425252
TALONT003425239
TALONT003425214
TALONT003425203
TALONT003425174
OAS1-211
OAS1-210
OAS1-208
OAS1-206
OAS1-203
OAS1-202
OAS1-201
112910000
112915000
112920000
112925000
112930000
chr12
OAS1−203
OAS1−201
OAS1−202
TALONT003425214
OAS1−211
TALONT003425239
TALONT003425174
OAS1−208
TALONT003425252
OAS1−206
TALONT003425890
OAS1−210
TALONT003427348
TALONT003427351
TALONT003425296
TALONT003425203

**Supplementary Figure 15. Summary of sjs-isoQTL effects on isoforms of *OAS1***
(**A**) Association between normalized expression of three eIsoforms of *OAS1* and the genotype of lead isoQTL, rs10774671. The violins and boxes are colored by genotype, and the grey line shows the trend of association. Center lines show the medians; the box indicates the middle of 50% of data; whiskers extend 1.5 times the interquartile range from the 25th and 75th percentiles; outliers are represented by dots; density of normalized expression is represented by the width of violin shape. (**B**) The normalized expressions of top 4 most abundant isoforms of *OAS1* at the individual level across lung cell types. Each dot represents an individual. (**C**) Structure of all *OAS1* isoforms based on the genomic coordinates. We validated the isoforms created by the alternative allele of rs1131454 as novel isoforms (*TALONT003425296* and *TALONT003425239* as alternative forms of 201 and 203, respectively). The exons included in each isoform are depicted by the boxes, which are colored by the structural categories. The arrows indicate the strand directions. (**D**) Normalized expressions of all post-QC isoforms for *OAS1* in alveolar macrophages. The bar shows the mean of normalized expression. Each dot represents the isoform expression of each individual within each cell type. The error bars show the mean of normalized expression ± standard deviation

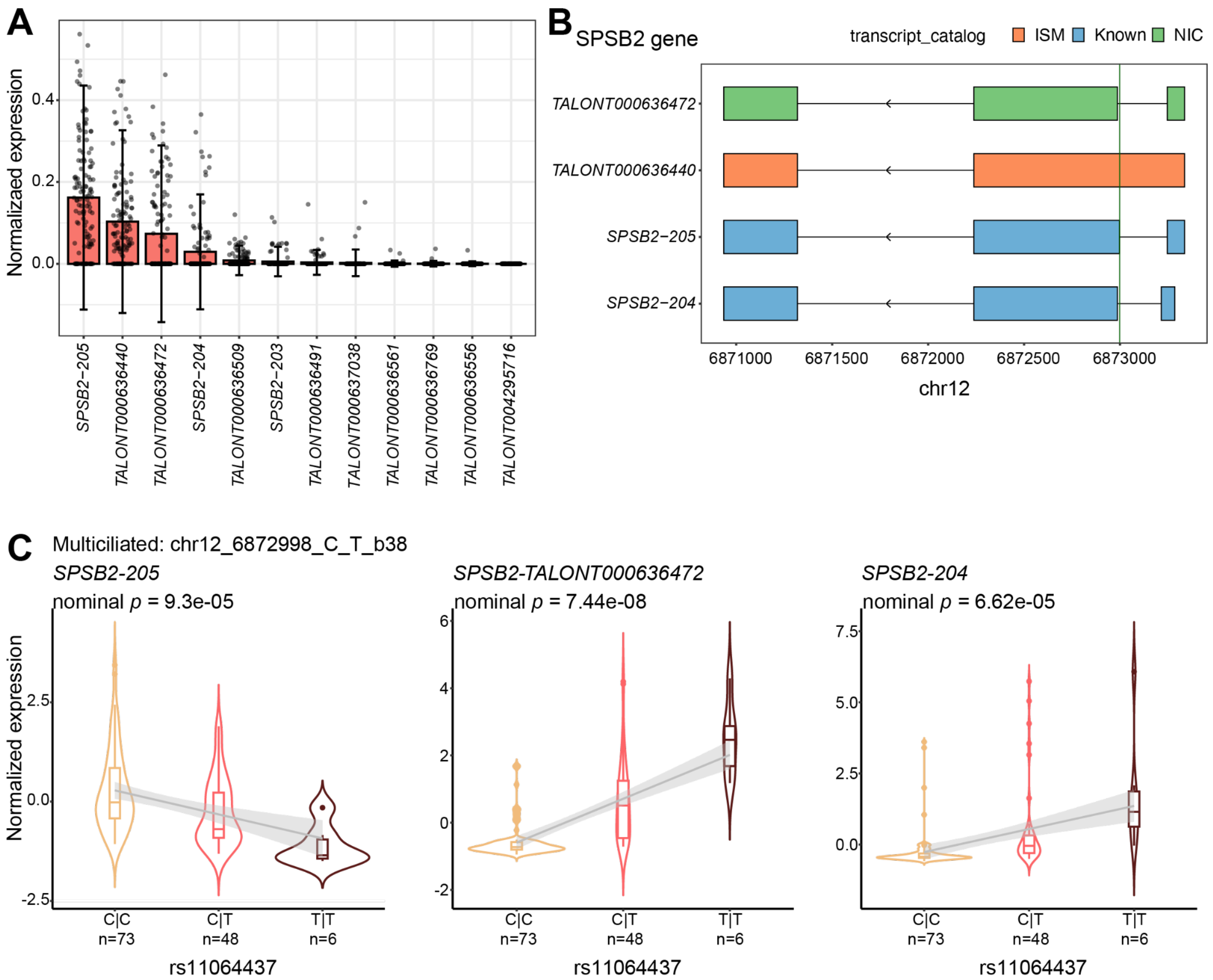


**Supplementary Figure 16. Summary of sjs-isoQTL effects on isoforms of *SPSB2***
(**A**) Normalized expressions of all post-QC isoforms for *SPSB2* in alveolar macrophages. (**B**) Structure of all *SPSB2* isoforms based on the genomic coordinates. The exons included in each isoform are depicted by the boxes, which are colored by the structural categories. The arrows indicate the strand directions. The novel isoform, *TALONT000636472*, is the most abundant isoform of *SPSB2* (**A**) that includes the splice junction altered by sjs-isoQTL, rs11064437 (shown as the green line). (**C**) Association between normalized expression of three *SPSB2* eIsoforms and the genotype of lead isoQTL, rs11064437.The violins and boxes are colored by genotype, and the grey line shows the trend of association. The lead isoQTL is shown at the top, with chromosome, position, reference allele, and alternative allele based on hg38 assembly. Center lines show the medians; the box indicates the middle of 50% of data; whiskers extend 1.5 times the interquartile range from the 25th and 75th percentiles; outliers are represented by dots; density of normalized expression is represented by the width of violin shape.

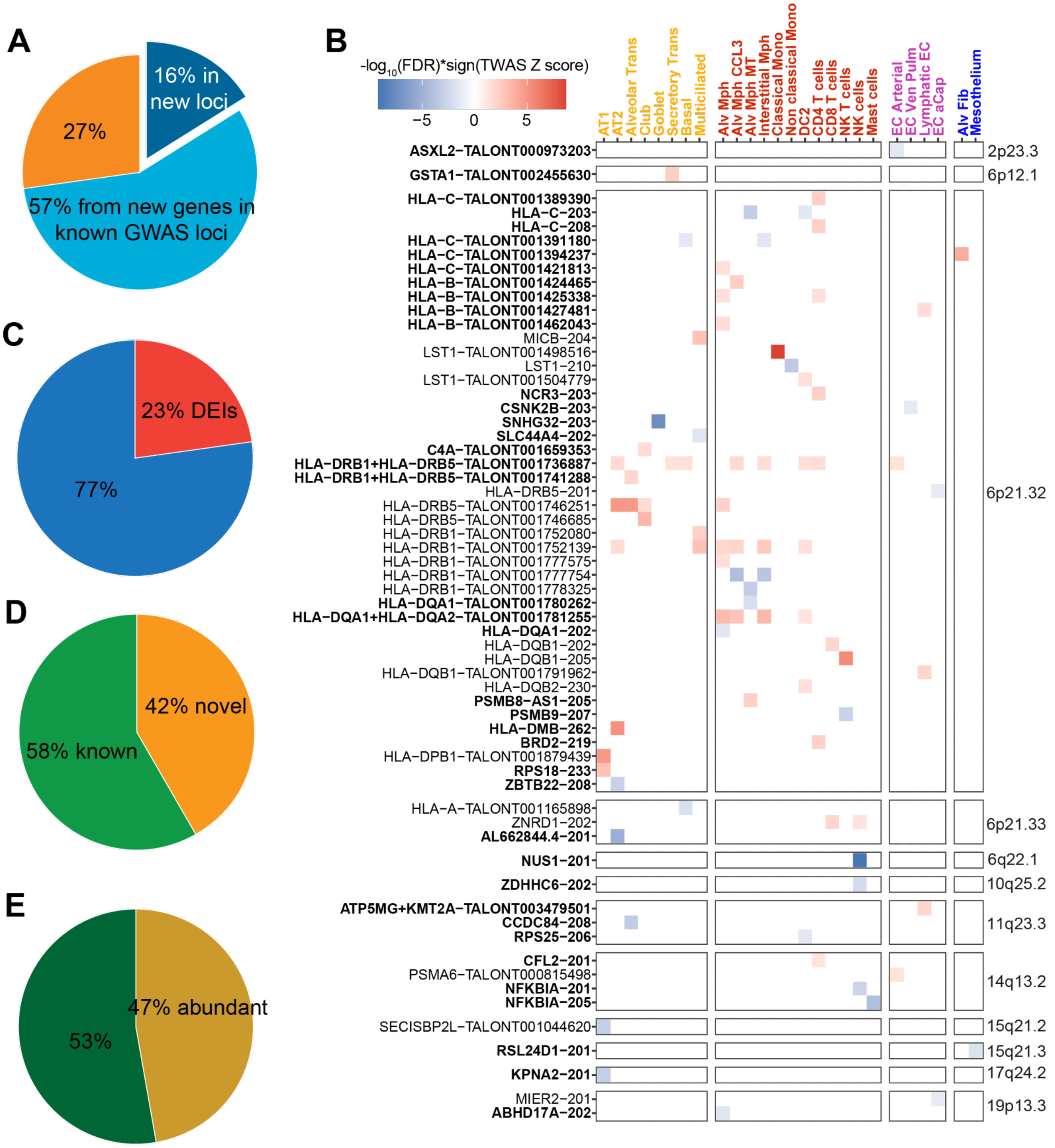


**Supplementary Figure 17. Isoform-level TWAS of LUAD from East Asian population.**

Pie charts present the fractions of TWAS isoforms from replicated or unreported genes or from new loci not significant in the GWAS (**A**), based on cell type specificity (**C**), based on isoform annotation status (known or unannotated in the reference data) (**D**), and based on isoform abundance (**E**). Abundant isoform is defined as a dataset-wide percentage of isoform within a gene larger than the 90th percentile of all isoforms (i.e., 14.4% as threshold). The cell type specificity of susceptibility isoforms was determined by DEI analysis. (**B**) Summary of TWAS results in LUAD GWAS significant loci from East Asian population. The color in the heat map shows the signed -$\log_{10}$FDR from TWAS (red: increased expression correlated with risk, blue: decreased expression correlated with risk). The heat map is ordered based on loci and cell categories. The isoforms from genes that have been reported by previous TWAS studies of lung cancer using bulk lung tissues are in bold.

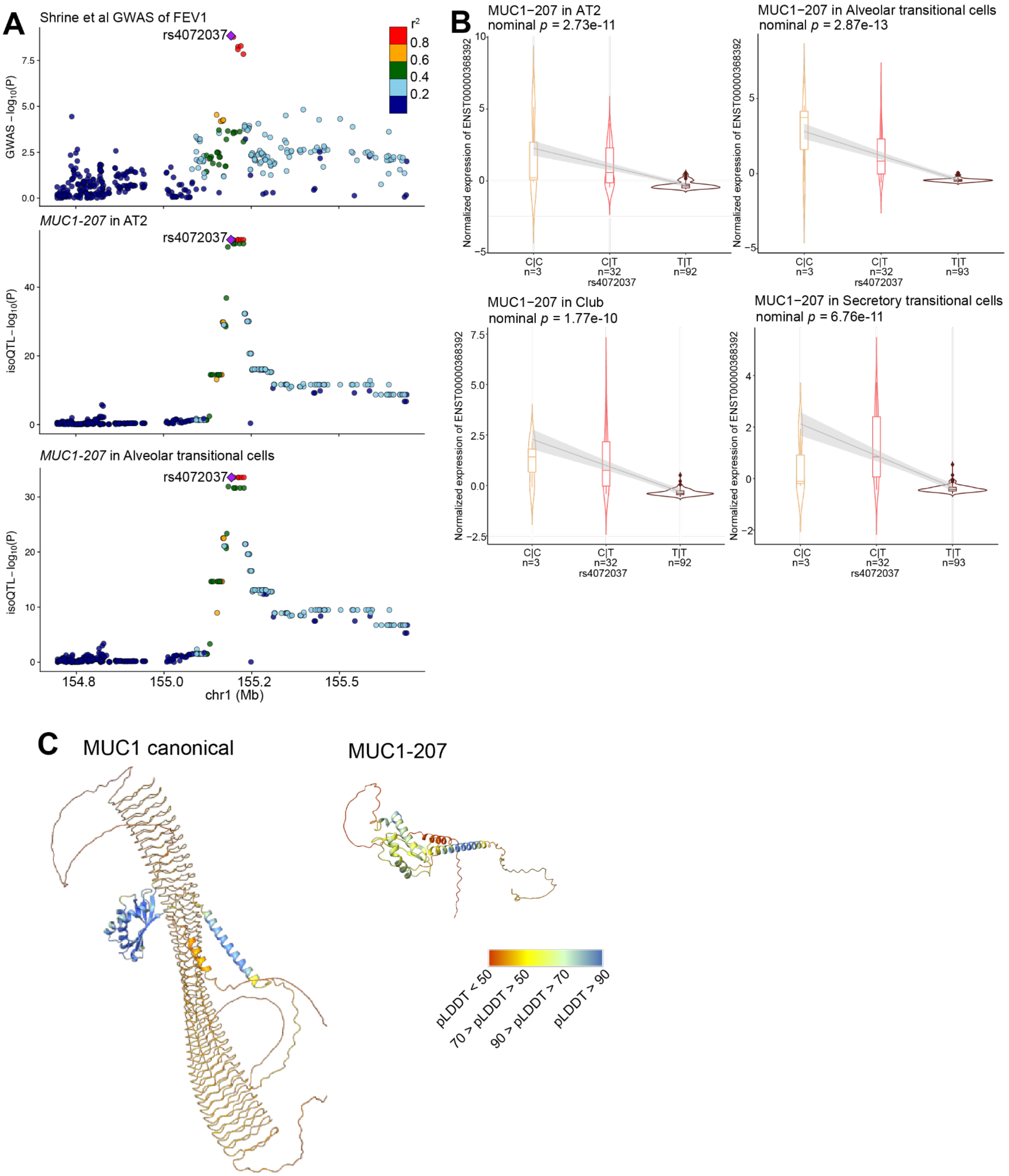


**Supplementary Figure 18. Summary of *MUC1-207* isoQTL**

(**A**) isoQTL colocalization with $FEV_1$ at the *MUC1* locus in AT2 and alveolar transitional cells. SNPs are color-coded based on the LD $R^2$ (1000 Genomes, EAS, phase 3) with the lead isoQTL, rs4072037 (purple diamond). (**B**) Association between the genotype of lead isoQTL rs4072037 and the normalized expression of *MUC1-207* in four epithelial cell types (*i.e.,* AT2, Alveolar transitional cells, club and secretory transitional cells). The violins and boxes are colored by genotype, and the grey line shows the trend of association. (**C**) Protein structures of canonical MUC1 and MUC1-207 predicted by AlphaFold3. The protein structure is colored by the predicted local distance difference test (pLDDT) score, where high confidence residues are in blue and low confidence residues are in red.

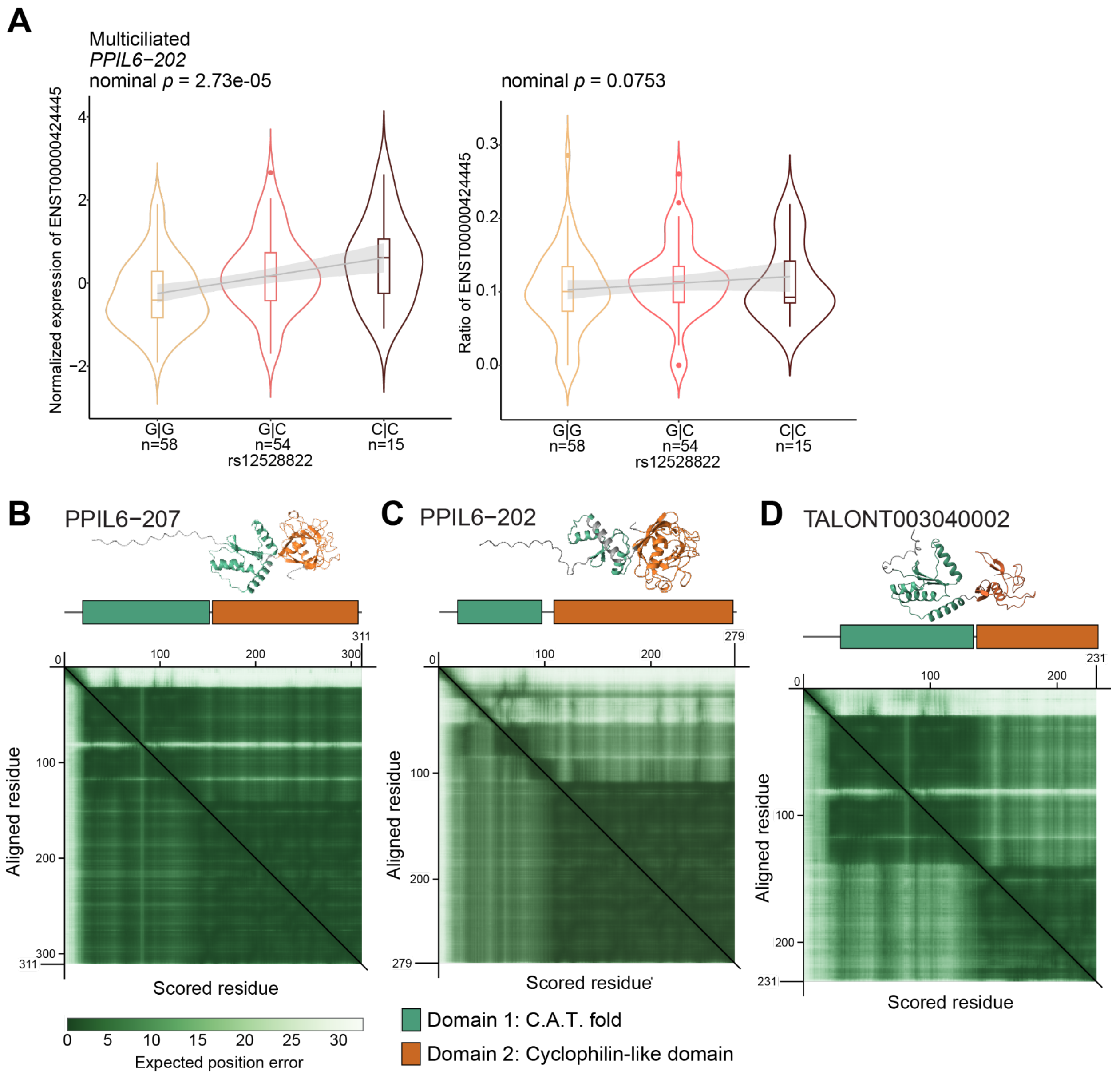


**Supplementary Figure 19. Predicted protein structures of PPIL6 isoforms and reciprocal allelic effects of *PPIL6-207* isoQTL**

(**A**) Association between the genotype of lead isoQTL rs12528822 and the normalized expression (left panel) and ratio of *PPIL6-202* among all *PPIL6* isoforms (right panel). P value is calculated by linear regression with the same covariates used in isoQTL mapping adjusted. Allele C of rs12528822 is the risk-associated allele for lung cancer. The violins and boxes are colored by genotype, and the grey line shows the trend of association. Center lines show the medians; the box indicates the middle of 50% of data; whiskers extend 1.5 times the interquartile range from the 25th and 75th percentiles; outliers are represented by dots; density of normalized expression is represented by the width of violin shape. Protein structures of PPIL6-207 (**B**), PPIL6-202 (**C**), and TALONT003040002 (**D**) predicted by AlphaFold3 with two main domains colored in green and orange. The heat map plots show the confidence in the relative position of the predicted and the true structures. The shade of green indicates the expected distance error in Ångströms. Dark green corresponds to a good prediction (low error), whereas light green indicates a poor prediction (high error).

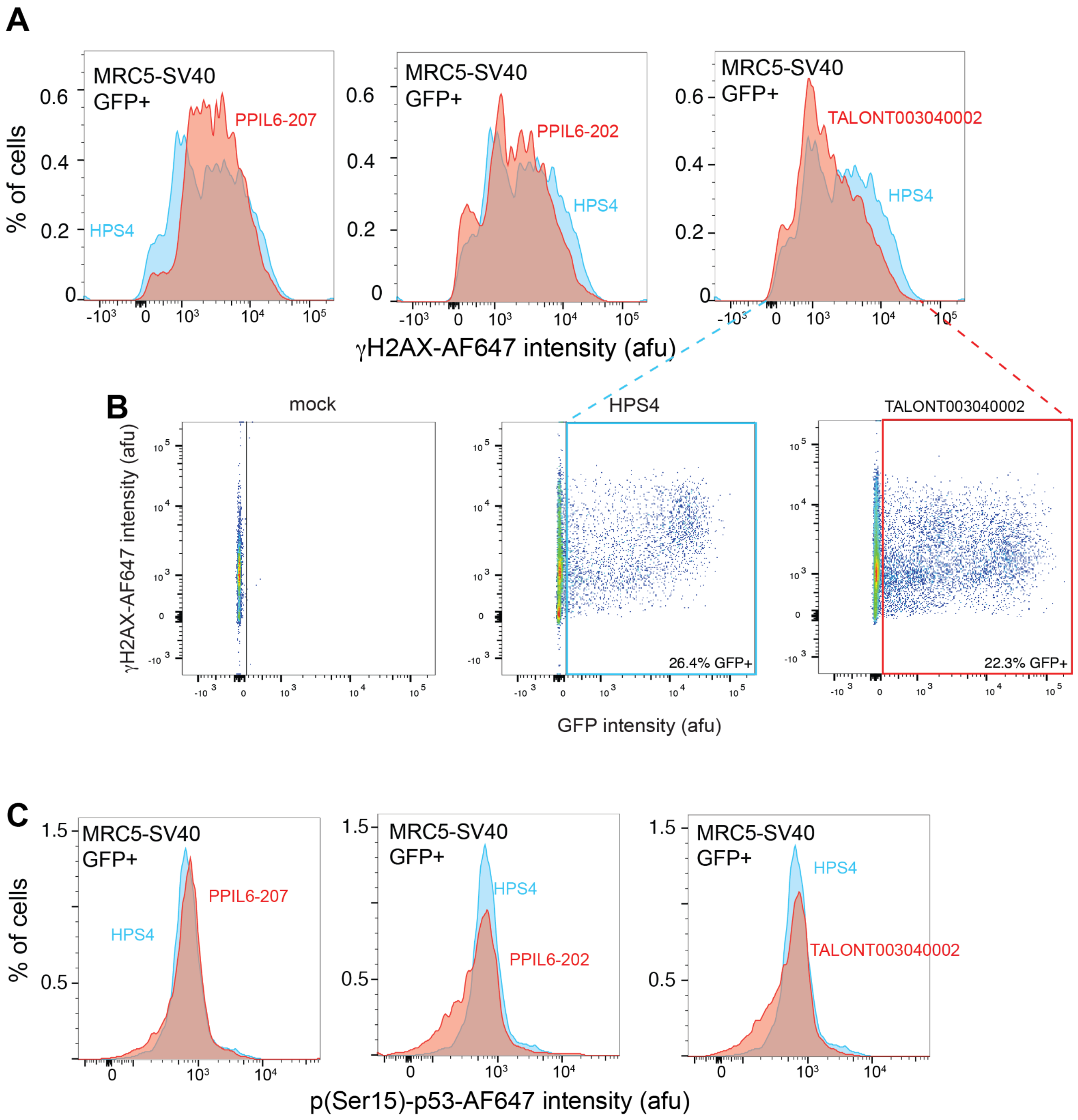


**Supplementary Figure 20. The novel PPIL6 isoform suppresses DNA damage.** **A**,**C**, Representative histograms of DNA damage markers(i.e., γH2AX or phospho-p53 (p-p53)) of each overproduced PPIL6 isoform or HPS4 in GFP$^+$ transfected MRC5-S40 cells from the same experimental batch. (**B**) Gating strategy of HPS4 or TALONT003040002 transfected cells associated with (**A**). Mock represents a transfection reagent without any isoform over-produced.

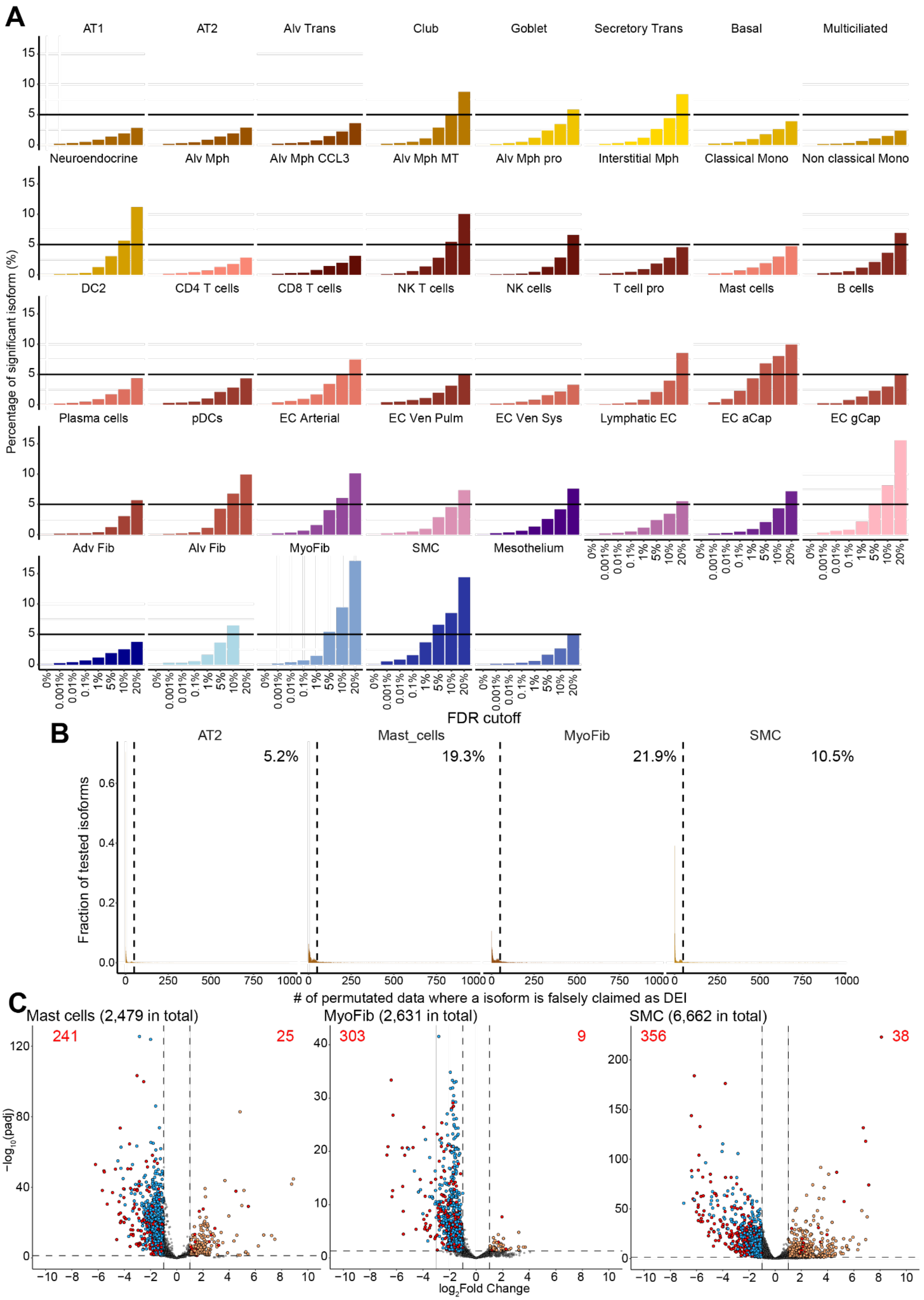
A
AT1
AT2
Alv Trans
Club
Goblet
Secretory Trans
Basal
Multiciliated
Neuroendocrine
Alv Mph
Alv Mph CCL3
Alv Mph MT
Alv Mph pro
Interstitial Mph
Classical Mono
Non classical Mono
DC2
CD4 T cells
CD8 T cells
NK T cells
NK cells
T cell pro
Mast cells
B cells
Plasma cells
pDCs
EC Arterial
EC Ven Pulm
EC Ven Sys
Lymphatic EC
EC aCap
EC gCap
Adv Fib
Alv Fib
MyoFib
SMC
Mesothelium
Percentage of significant isoform (%)
FDR cutoff
0%
0.001%
0.01%
0.1%
1%
5%
10%
20%
B
AT2
Mast_cells
MyoFib
SMC
5.2%
19.3%
21.9%
10.5%
Fraction of tested isoforms
### of permutated data where a isoform is falsely claimed as DEI
C
Mast cells (2,479 in total)
MyoFib (2,631 in total)
SMC (6,662 in total)
241
25
303
9
356
38
−log10(padj)
log2Fold Change

**Supplementary Figure 21. FDR control for differentially expressed isoform analysis**

(**A**) Proportions of isoforms falsely claimed as significant in the permuted data of each cell type while using varied cutoffs of FDR. Cell type names are shown at the top of each plot. (**B**) Fraction of isoforms that were falsely claimed as significant for different times in 1,000 times of permutations for three cell types (i.e., mast cells, myofibroblast, and SMC) that displayed >5% false detection at the 5% FDR cutoff from (**A**). AT2 was shown as a control. (**C**) Volcano plots show the relationship between DEI fold change and adjusted p values from the real data of mast cells, myofibroblasts, and SMC. Each dot represents an isoform. The up-regulated isoforms are colored in orange and down-regulated in blue. The potential false positives (isoforms falsely claimed as significant more than 50 times in 1000-time permutations) are colored in red. The total numbers of false positive isoforms from up-regulated and down-regulated sides are shown on the plots, which indicates that most of the false positives are down-regulated in that cell type and not considered as final DEIs.
